# Public broadly neutralizing antibodies against hepatitis B virus in individuals with elite serologic activity

**DOI:** 10.1101/2020.03.04.976159

**Authors:** Qiao Wang, Eleftherios Michailidis, Yingpu Yu, Zijun Wang, Arlene M. Hurley, Deena A. Oren, Christian T. Mayer, Anna Gazumyan, Zhenmi Liu, Yunjiao Zhou, Till Schoofs, Kai-hui Yao, Jan P. Nieke, Jianbo Wu, Qingling Jiang, Chenhui Zou, Mohanmmad Kabbani, Corrine Quirk, Thiago Oliveira, Kalsang Chhosphel, Qianqian Zhang, William M. Schneider, Cyprien Jahan, Tianlei Ying, Jill Horowitz, Marina Caskey, Mila Jankovic, Davide F. Robbiani, Yumei Wen, Ype P. de Jong, Charles M. Rice, Michel C. Nussenzweig

**Affiliations:** Key Laboratory of Medical Molecular Virology (MOE/NHC/CAMS), School of Basic Medical Sciences, Shanghai Medical College, Fudan University, Shanghai 200032, China; Laboratory of Virology and Infectious Disease, The Rockefeller University, New York, NY 10065, USA; Laboratory of Molecular Immunology, The Rockefeller University, New York, NY 10065, USA; Structural Biology Resource Center, The Rockefeller University, New York, NY 10065, USA; West China School of Public Health, West China Hospital, Sichuan University, Chengdu 610041, China; Howard Hughes Medical Institute, The Rockefeller University, New York, NY 10065, USA; Division of Gastroenterology and Hepatology, Weill Cornell Medicine, New York, NY 10065

**Author notes:** These authors contributed equally (Qiao Wang, Eleftherios Michailidis). These authors contributed equally (Charles M. Rice, Michel C. Nussenzweig). Corresponding authors (Q.W.), (Y.P.J.).

**Keywords:** Hepatitis B infection, elite neutralizing activity, broadly neutralizing antibodies, humanized mice, crystal structure, prophylaxis, passive immunotherapy, escape mutations

## Abstract

**SUMMARY:** Although there is no effective cure for chronic hepatitis B virus (HBV) infection, antibodies are protective and constitute clinical correlates of recovery from infection. To examine the human neutralizing antibody response to HBV in elite neutralizers we screened 144 individuals. The top individuals produced shared clones of broadly neutralizing antibodies (bNAbs) that targeted 3 non-overlapping epitopes on the HBV S antigen (HBsAg). Single bNAbs protected humanized mice against infection, but selected for resistance mutations in mice with established infection. In contrast, infection was controlled by a combination of bNAbs targeting non-overlapping epitopes with complementary sensitivity to mutations that commonly emerge during human infection. The co-crystal structure of one of the bNAbs with a peptide epitope containing residues frequently mutated in human immune escape variants revealed a loop anchored by oppositely charged residues. The structure provides a molecular explanation for why immunotherapy for HBV infection may require combinations of complementary bNAbs.

## INTRODUCTION

Despite the existence of effective vaccines, hepatitis B virus (HBV) infection remains a major global health problem with an estimated 257 million people living with the infection. Whereas 95% of adults and 50-75% of children between the ages of 1 and 5 years spontaneiously control HBV, only 10% of infants recover naturally. The remainder develop a chronic infection that can lead to liver cirrhosis and hepatocellular carcinoma. Although chronic infection can be suppressed with antiviral medications, there is no effective curative therapy (Dienstag, 2008; Revill et al., 2016; Thomas, 2019).

HBV is an enveloped double stranded DNA virus of the *Hepadnaviridae* family. Its genome is the smallest genome among pathogenic human DNA viruses, with only four open reading frames. Infected liver cells produce both infectious HBV virions (Dane particles) and non-infectious subviral particles (Australia antigen) (Dane et al., 1970; Hu and Liu, 2017). The virion is a 42 nm-diameter particle containing the viral genome and HBV core antigen (HBcAg) encapsidated by a lipid membrane containing the hepatitis B surface antigen (HBsAg) (Blumberg, 1964; Venkatakrishnan and Zlotnick, 2016). Subviral particles lack the viral genome.

Antibodies to HBsAg (anti-HBs) are associated with successful vaccination and recovery from acute infection, while antibodies to HBcAg (anti-HBc) are indicative of past or current HBV infection (Ganem, 1982). Indeed, the most significant difference between chronically infected and naturally recovered individuals is a robust antibody response to HBsAg (Ganem, 1982) (Figure S1A). Conversely, the inability to produce these antibodies during acute infection is associated with chronicity (Trepo et al., 2014). Whether these associations reflect an etiologic role for anti-HBs antibodies in protecting from or clearing established infection is not known. However, depletion of antibody-producing B lymphocytes in exposed humans by anti-CD20 therapies (e.g. rituximab) is associated with HBV reactivation, indicating that B cells and/or their antibody products play a significant role in controlling the infection (Loomba and Liang, 2017).

Several human antibodies against HBsAg have been obtained using a variety of methods including: phage display (Kim and Park, 2002; Li et al., 2017; Sankhyan et al., 2016; Wang et al., 2016); humanized mice (Eren et al., 1998); Epstein-Barr virus-induced B cell transformation (Heijtink et al., 2002; Heijtink et al., 1995; Sa’adu et al., 1992); hybridoma technology (Colucci et al., 1986); human B cell cultures (Cerino et al., 2015); and microwell array chips (Jin et al., 2009; Tajiri et al., 2010). However, the donors in these studies were not selected for serum neutralizing activity.

Here, we report on the human humoral immune response to HBsAg in immunized and spontaneously recovered individuals that have been selected for high levels of serum neutralizing activity. We find that these individuals develop closely related bNAbs that target shared non-overlapping epitopes in HBsAg. The crystal structure of one of the antibodies with its peptide target reveals a loop that helps to explain why certain amino acid residues are frequently mutated in naturally arising escape viruses and why combinations of bNAbs may be needed to control infection. *In vivo* experiments in humanized mice demonstrate that the bNAbs are protective and can be therapeutic when used in combination.

## RESULTS

### Serologic Responses Against HBV

To select individuals with outstanding antibody responses to HBsAg we performed ELISA assays on serum obtained from 159 volunteers. These included 15 uninfected and unvaccinated controls (HBsAg^−^, anti-HBs^−^, anti-HBc^−^), 20 infected and spontaneously recovered (HBsAg^−^, anti-HBs^+/−^, anti-HBc^+^), and 124 vaccinated (HBsAg^−^, anti-HBs^+/−^, anti-HBc^−^) volunteers. These individuals displayed a broad spectrum of anti-HBs titers (x-axis in Figure 1A and Figure S1B; Table S1). To determine their neutralizing activity, we tested their ability to block HBV infection in sodium taurocholate co-transporting polypeptide (NTCP)-overexpressing HepG2 cells (Michailidis et al., 2017; Yan et al., 2012) (y-axis in Figure 1A and Figure S1B and S1C; Table S1). Sera or antibodies purified from individuals with high neutralizing titers (elite neutralizers) were then compared across a wide range of dilutions (Figure 1B and 1C). Although anti-HBs ELISA titers positively correlated with neutralizing activity (r_s_=0.492, p<0.001, Spearman’s rank correlation), there were notable exceptions as exemplified by volunteers #99 and #49, whose sera failed to neutralize HBV despite high anti-HBs ELISA titers (Figure 1A). Thus, ELISA titers against HBsAg are not entirely predictive of neutralizing activity *in vitro*.

**Figure 1.**
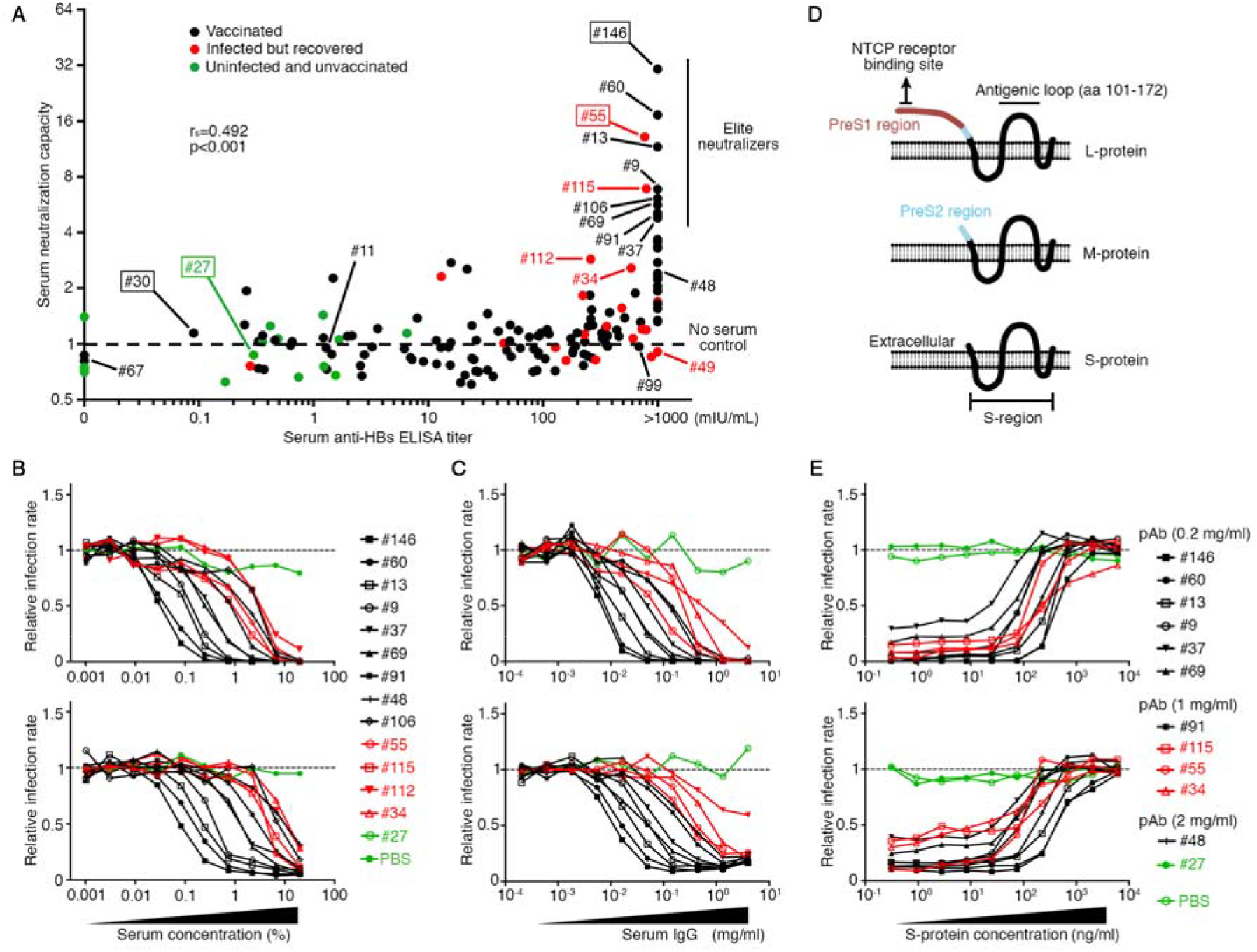
Antibody responses in HBV vaccinated and recovered individuals. (A) Donor screen. Sera from 159 volunteers were evaluated for anti-HBs binding by ELISA (x-axis) and HBV serum neutralization capacity using HepG2-NTCP cells (y-axis). Neutralization tests were performed at 1:5 serum dilution in the final assay volume. Each dot represents an individual donor. Green indicates unvaccinated and unexposed, black indicates vaccinated, and red indicates spontaneously recovered. The dashed line indicates the no serum control. Elite neutralizers are indicated (top right). Boxed are representative samples shown in Figure 2A. Spearman’s rank correlation coefficient (r_s_) and significance value (p). (B and C) Dose-dependent HBV neutralization by serum (B) or by purified IgG (C). Two assays were used to measure infection rates: ELISA to measure HBsAg protein in the medium (upper panels) and immunofluorescent staining for HBcAg in HepG2-NTCP cells (lower panels). Dashed line indicates saturation. (D) Schematic representation illustrating the three forms of the HBV surface protein: L-, M- and S-protein. These three forms of envelope protein all share the same S-region, with PreS1/PreS2 and PreS2 alone as the N-terminal extensions for L- and M-protein, respectively. (E) S-protein produced in Chinese hamster ovary (CHO) cells blocks serum neutralizing activity. Graphs show infection rate as a function of the amount of S-protein added. The concentration of polyclonal IgG antibodies (pAb) is indicated. Upper and lower panels are as in (B) and (C). A representative of at least two experiments is shown.

**Figure 2.**
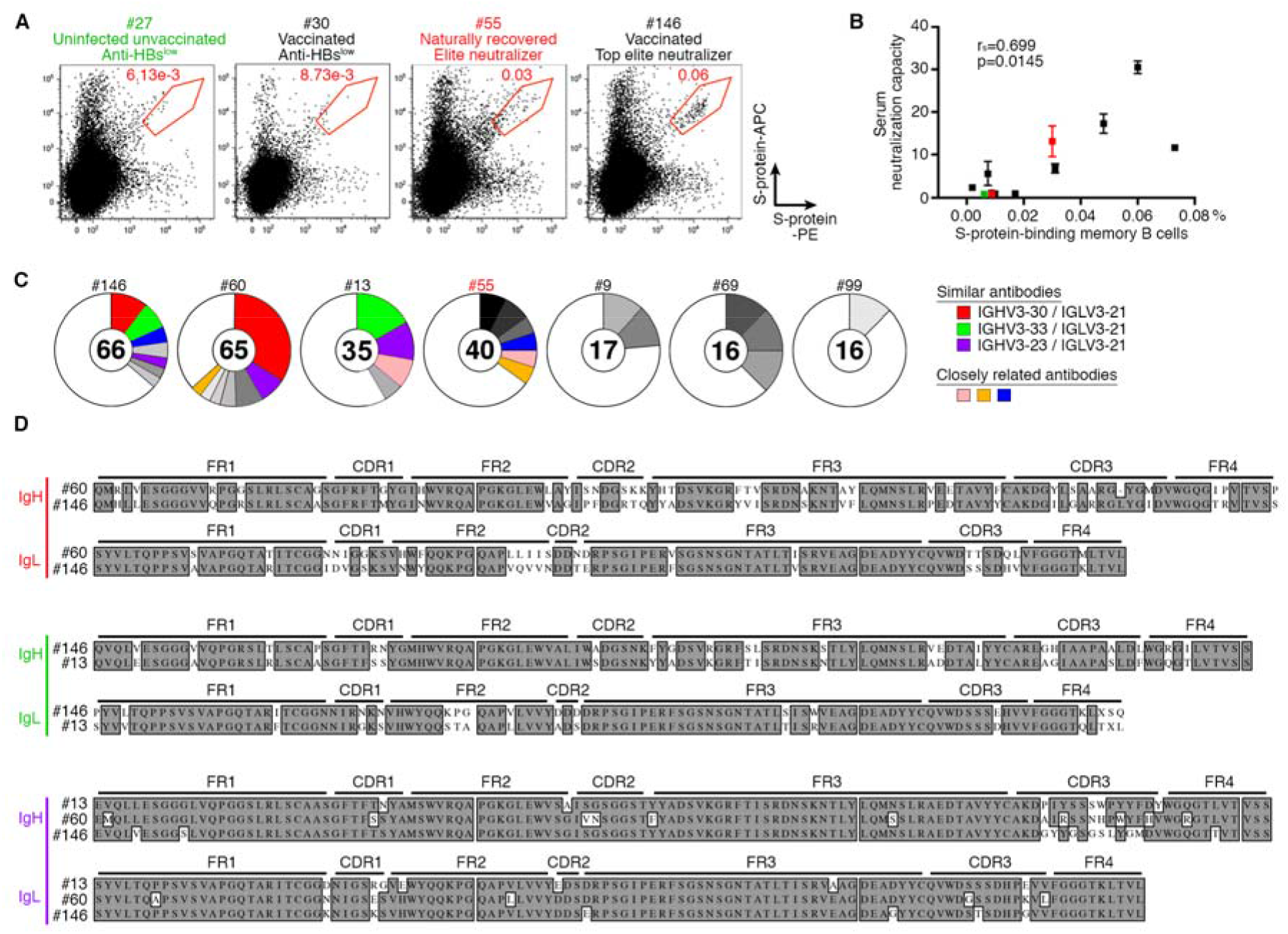
S-protein-specific antibodies. (A) Frequency of S-protein-specific memory B cells.Representative flow cytometry plots displaying the percentage of all IgG^+^ memory B cells that bind to both allophycocyanin- and phycoerythrin-tagged S-protein (S-protein-APC and S-protein-PE). Flow cytometry plots from other individuals are shown in Figure S2A. Experiments were repeated two times. (B) Dot plot showing the correlation between the frequency of S-protein-binding IgG^+^ memory B cells and the serum neutralizing activity. Spearman’s rank correlation coefficient (r_s_) and significance value (p). (C) Each pie chart represents the antibodies from an individual donor, and the total number of sequenced antibodies with paired heavy and light chains is indicated in the center. Antibodies with the same combination of IGH and IGL variable gene sequences and closely related CDR3s are shown as one slice with colors indicating shared sequences between individuals. Grey slices indicate antibodies with closely related sequences that are unique to a single donor. In white are singlets. (D) V(D)J alignments for representative IGHV3-30/IGLV3-21, IGHV3-33/IGLV3-21 and IGHV3-23/IGLV3-21 antibodies from donors #60/#146, #146/#13, and #13/#60/#146 respectively. Boxed grey residues are shared between antibodies.

The HBV surface protein, HBsAg, is a four-transmembrane protein that can be subdivided into PreS1-, PreS2- and S-regions (Figure 1D). To determine which of these regions is the dominant target of the neutralizing response in elite individuals we used S-protein to block neutralizing activity *in vitro*. As expected, the neutralizing activity in volunteers that received the HBV vaccine, which is composed of S-protein, was completely blocked by S-protein (black lines in Figure 1E). The same was true for the spontaneously recovered individuals in our cohort despite a reported ability of this population to produce anti-PreS1 or anti-PreS2 antibodies (Coursaget et al., 1988; Li et al., 2017; Sankhyan et al., 2016) (red lines in Figure 1E). These results suggest that the neutralizing antibody response in elite individuals is directed primarily against the S-protein irrespective of immunization or infection.

### Human Monoclonal Antibodies to HBV

To characterize the antibodies responsible for elite neutralizing activity we purified S-protein binding class-switched memory B cells (Escolano et al., 2019; Scheid et al., 2009a). Unexposed naive controls and vaccinated individuals with low anti-HBs ELISA titers showed background levels of S-protein specific memory B cells (Figure 2A and S2A). In contrast, elite neutralizers displayed a distinct population of S-antigen binding B cells constituting 0.03-0.07% of the IgG^+^ memory compartment (CD19-MicroBeads^+^ CD20-PECy7^+^ IgG-Bv421^+^ S-protein-PE^+^ S-protein-APC^+^ ovalbumin-Alexa Fluor 488^−^) (Figure 2A and S2A). Consistent with the findings in elite HIV-1 neutralizers (Rouers et al., 2017), the fraction of S-protein specific cells was directly correlated to the neutralization titer of the individual (r_s_=0.699, p=0.0145, Spearman’s rank correlation) (Figure 2B).

Immunoglobulin heavy (*IGH*) and light (*IGL* or *IGK*) chain genes were amplified from single memory B cells by PCR (Robbiani et al., 2017; Scheid et al., 2009b; von Boehmer et al., 2016). Overall, we obtained 244 memory B cell antibodies from eight volunteers with high anti-HBs ELISA titers (Figure S2B and S2C; Table S2). Expanded clones composed of cells producing antibodies encoded by the same *Ig* variable gene segments with closely related CDR3s were found in elite neutralizers #146, #60 and #13 (Figure 2C). For example, IGHV3-30/IGLV3-21 was present in #146 and #60; IGHV3-33/IGLV3-21 in #146 and #13; and IGHV3-23/IGLV3-21 in #146, #60 and #13. These antibodies were approximately 80% identical at the amino acid level (Figure 2D). Antibodies with closely related *Ig* heavy and light chains were also identified between volunteer #55 (HBV infected but recovered) and vaccinated individuals (Figure 2C and S2B). We conclude that elite HBV neutralizers produce clones of antigen-binding B cells that express closely related *Ig* heavy and light chains.

### Breadth of Reactivity

Twenty representative antibodies from 5 individuals, designated as H001 to H020, were selected for expression and further testing (Figure S2B). All 20 antibodies showed reactivity to the S-protein antigen used for B cell selection (HBsAg adr CHO) by ELISA with 50% effective concentration (EC_50_) values ranging from 18-350ng/ml (Figure 3A). These antibodies carried somatic hypermutations that enhanced antigen binding as determined by reversion to the inferred germline (Figure 3B). Thus, affinity maturation was essential for their high binding activity.

**Figure 3.**
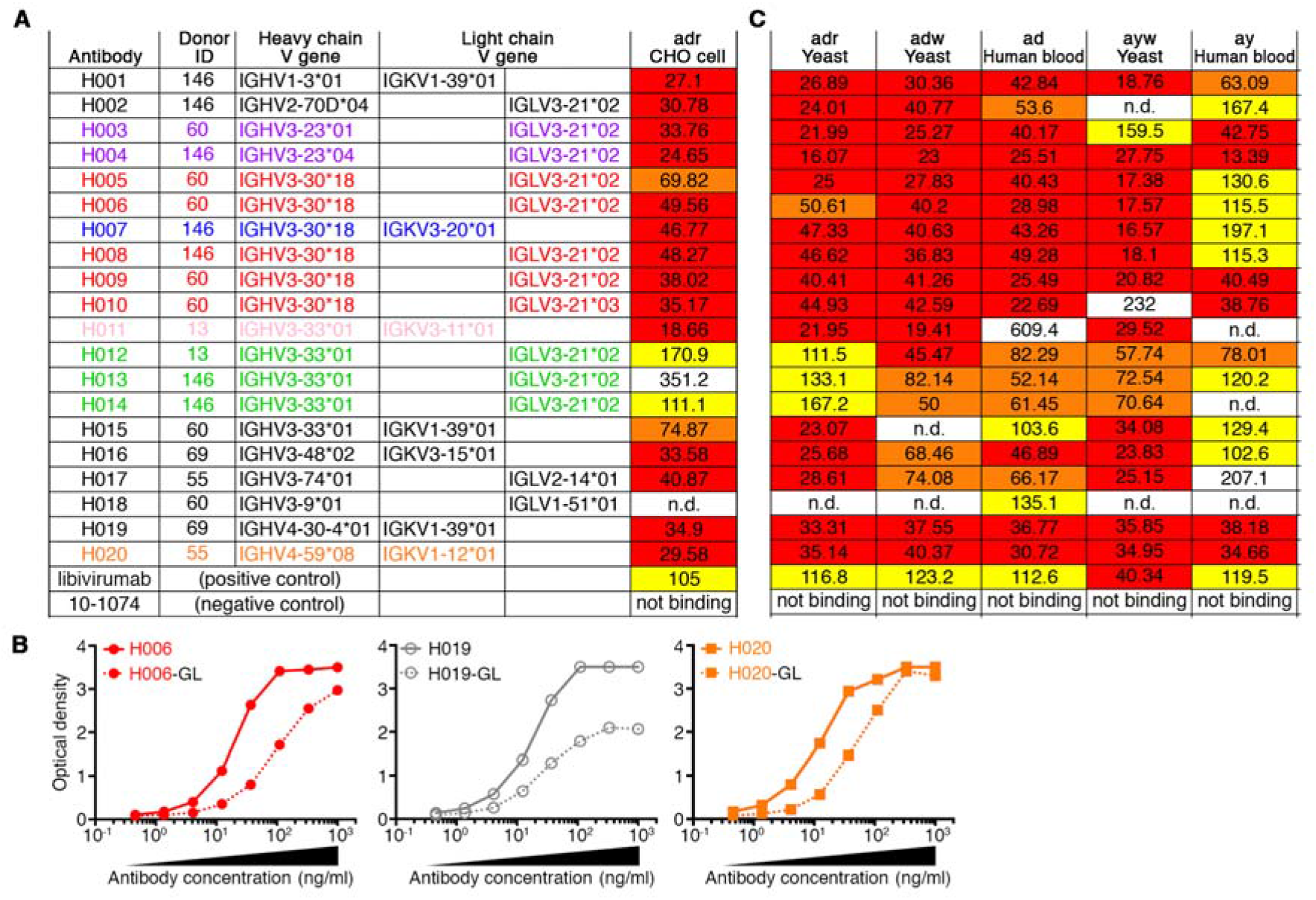
Broad cross-reactivity. (A) Binding to S-protein (adr serotype). 50% effective concentration (EC_50_ in ng/ml) required for binding of the indicated human monoclonal antibodies to the S-protein. Libivirumab (Eren et al., 2000; Eren et al., 1998) and anti-HIV antibody 10-1074 (Mouquet et al., 2012) were used as positive and negative controls, respectively. (B) Comparative binding of the mature and inferred germlines (GL) of antibodies H006, H019, and H020 to S-protein by ELISA. Anti-HBs antibody binding to 5 different serotypes of HBsAg. Similar to panel (A), EC_50_ values are color-coded: red, ≤50 ng/ml; orange, 50 to 100 ng/ml; yellow, 100 to 200 ng/ml; and white, > 200 ng/ml. All experiments were performed at least two times.

Four major serotypes of HBV exist as defined by a constant “a” determinant and two variable and mutually exclusive determinants “d/y” and “w/r” (Bancroft et al., 1972; Le Bouvier, 1971) with a highly statistically significant association between serotypes and genotypes (Kramvis et al., 2008; Norder et al., 2004). To determine whether our antibodies cross-react to different HBsAg serotypes, we performed ELISAs with 5 additional HBsAg antigens: yeast-expressed serotype “adr”, “adw”, and “ayw”, as well as “ad” and “ay” antigen purified from human blood (Figure 3C). Many of the antibodies tested displayed broad cross-reactivity and EC_50_ values lower than libivirumab, a human anti-HBs monoclonal antibody that was isolated from HBV-immunized humanized mice and then tested clinically (Eren et al., 2000; Eren et al., 1998; Galun et al., 2002). These antibodies were not polyreactive or autoreactive (Figure S3A and S3B). We conclude that the antibodies obtained from elite neutralizers are broadly cross-reactive with different HBV serotypes.

### Antigenic Epitopes on S-protein

To determine whether the 20 selected antibodies bind to overlapping or non-overlapping epitopes we performed competition ELISA assays, in which the S-protein was pre-incubated with a selected antibody followed by a second biotinylated antibody. As expected, all antibodies blocked the binding of the autologous biotinylated monoclonal (yellow boxes in Figure 4A), while control human anti-HIV antibody 10-1074 failed to block any of the anti-HBs antibodies. The competition ELISA identified three mutually exclusive groups of monoclonal antibodies, suggesting that there are at least three dominant non-overlapping antigenic sites on HBsAg (red box for Group-I, blue box for Group-II, and H017 in Group-III, Figure 4A). The top 4 elite individuals that had 2 or more antibodies tested in the competition ELISA expressed monoclonal antibodies that targeted 2 of the 3 non-overlapping epitopes (Figure 4A and S2B).

**Figure 4.**
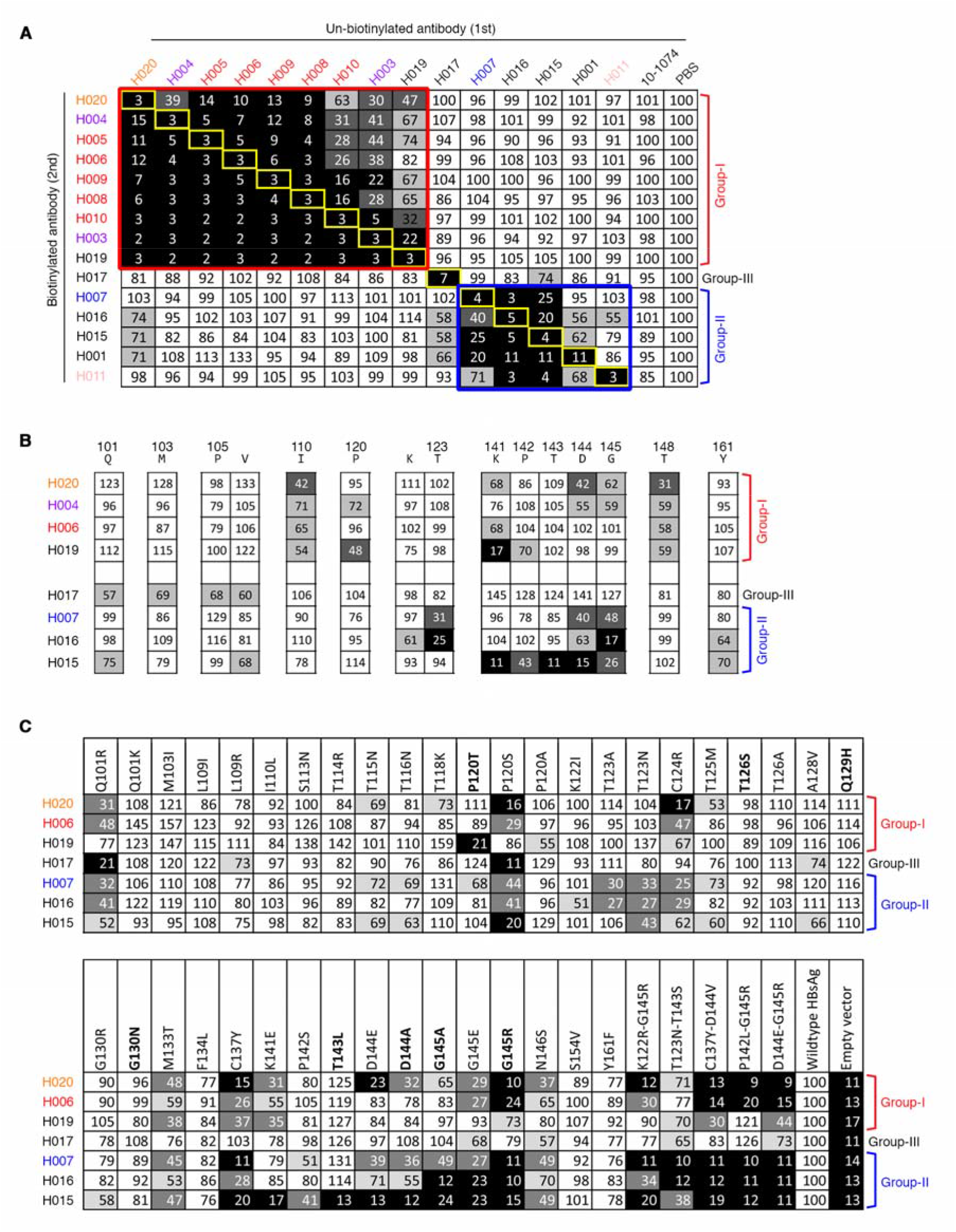
HBsAg epitopes. (A) Competition ELISA defines 3 groups of antibodies. Results of competition ELISA shown as percent of binding by 2nd biotinylated antibodies and illustrated by colors: black, 0-25%; dark grey, 26-50%; light grey, 51-75%; white, >76%. Representative of two experiments. (B) Results of ELISA on alanine scanning mutants of S-protein. Only the amino acids vital for antibody binding are shown. Binding to mutants relative to wild-type S-protein: black, 0-25%; dark grey, 26-50%; light grey, 51-75%; white, >75%. Details are in Figure S4. (C) Results of ELISA on various escape mutations of S-protein. Wild-type S-protein and empty vector serve as a positive and negative controls, respectively. Binding to mutants relative to wild-type S-protein: black, 0-25%; dark grey, 26-50%; light grey, 51-75%; white, >75%. Amino acid mutations in bold represent frequently observed mutations in humans (Ma and Wang, 2012). All experiments were performed at least two times.

To define these epitopes more precisely, we produced a series of alanine mutants spanning most of the predicted extracellular domain of the S-protein with the exception of cysteines, alanines, and amino acid residues critical for S-protein production (Salisse and Sureau, 2009) (Figure 1D and S4A). ELISA assays with the mutant proteins revealed a series of binding patterns corresponding to the three groups defined in the competition assays (Figure 4B and S4B). For example, mutations I110A and T148A interfered with binding by Group-I antibodies exemplified by H004, H006, H019, and H020, but had little measurable effect on Group-II antibodies exemplified by H007, H015, and H016 or Group-III antibody H017 (Figure 4B and S4B). In addition, alanine scanning suggested that residues D144 and G145 are critical antigenic determinants for many of the monoclonals.

In addition to alanine scanning we also produced 44 common naturally occurring escape variants found in chronically infected individuals (Hsu et al., 2015; Ijaz et al., 2012; Ma and Wang, 2012; Salpini et al., 2015). Whereas alanine scanning showed that some of the antibodies in Group-I and -II were resistant to G145A, the corresponding naturally occurring mutations at the same position, G145E and G145R, revealed decreased binding by most antibodies (Figure 4C). Among the antibodies tested, H017 and H019, in Groups-III and -I respectively, showed the greatest resistance to G145 mutations and the greatest breadth and complementarity (Figure 4C). We conclude that human anti-HBs monoclonals obtained from elite individuals recognize distinct epitopes on HBsAg, most of which appear to be non-linear conformational epitopes spanning different regions of the protein.

### *In Vitro* Neutralizing Activity

To determine whether the new monoclonals neutralize HBV *in vitro*, we performed neutralization assays using HepG2-NTCP cells (Figure 5A and 5B). The 50% inhibitory concentration (IC_50_) values were calculated based on HBsAg/HBeAg ELISA or immunofluorescence staining for HBcAg expression (Michailidis et al., 2017) (Figure 5C). Neutralizing activity was further verified by *in vitro* neutralization assays using primary human hepatocytes (Michailidis et al., 2020) (Figure 5C and 5D). Fourteen of the 20 antibodies tested showed neutralizing activity with IC_50_ values as low as 5 ng/ml (Figure 5C). By comparison, libivirumab had an IC_50_ of 35 and 128 ng/ml in the neutralization assays based on ELISA and immunofluorescence assays respectively (Figure 5C). Somatic hypermutations were essential for potent neutralizing activity as illustrated by the reduced activity of inferred germline antibodies (Figure S5A and S5B). In addition, neutralizing activity was dependent on bivalent binding since the IC_50_ values for Fab fragments were 2 orders of magnitude higher than intact antibodies (Figure 5E). Finally, there was no synergy when Group-I, -II, and -III antibodies were combined (Figure S5C). We conclude that half of the new monoclonals were significantly more potent than libivirumab including Group-I H004, H005, H006, H008, H009, H019, and H020 and Group-II H007, H015, and H016 (Figure 5C).

**Figure 5.**
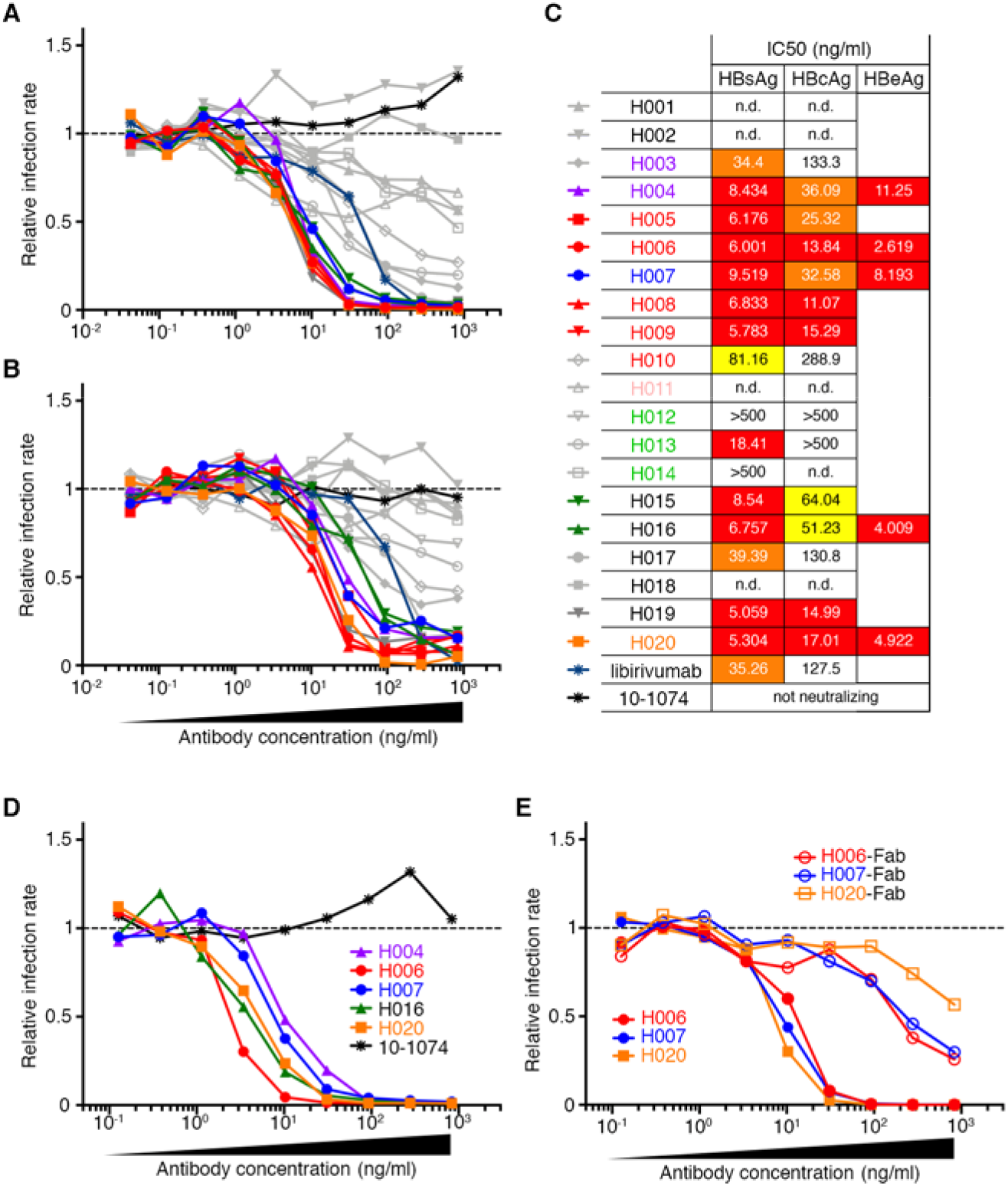
*In vitro* neutralization by the monoclonal antibodies. (A and B) *In vitro* neutralization assays using HepG2-NTCP cells. Percent infection in the presence of the indicated concentrations of bNAbs measured by ELISA of HBsAg in medium (A) and anti-HBcAg immunofluorescence (B). Anti-HIV antibody 10-1074 (Mouquet et al., 2012) and libivirumab (Eren et al., 2000; Eren et al., 1998) were used as negative and positive controls respectively. All experiments were repeated a minimum of two times. (C) bNAb 50% maximal inhibitory concentration (IC_50_) calculated based on HBsAg ELISA (left) or HBcAg immunofluorescence (middle) or HBeAg ELISA (right). *In vitro* neutralization using primary human hepatocytes. The levels of HBeAg in medium were measured by ELISA. The calculated IC_50_ values are shown in the right column of panel (C). Experiments were repeated three times. (E) *In vitro* neutralization assay using HepG2-NTCP cells. IgG antibodies were compared to their corresponding Fab fragments. Concentrations of Fab fragments were adjusted to correspond to IgG. Experiment was performed two times.

### Structure of the H015 Antibody/Peptide Complex

H015 differed from other antibodies in that its binding was inhibited by 5 consecutive alanine mutations spanning positions K141-G145 indicating the existence of a linear epitope. This idea was verified by ELISA against a series of overlapping peptides comprising the predicted extracellular domain of S-protein (Figure 6A and S6A). The data showed that H015 binds to KPSDGN, which is near the center of the putative extracellular domain and contains some of the most frequently mutated amino acids during natural infection.

**Figure 6.**
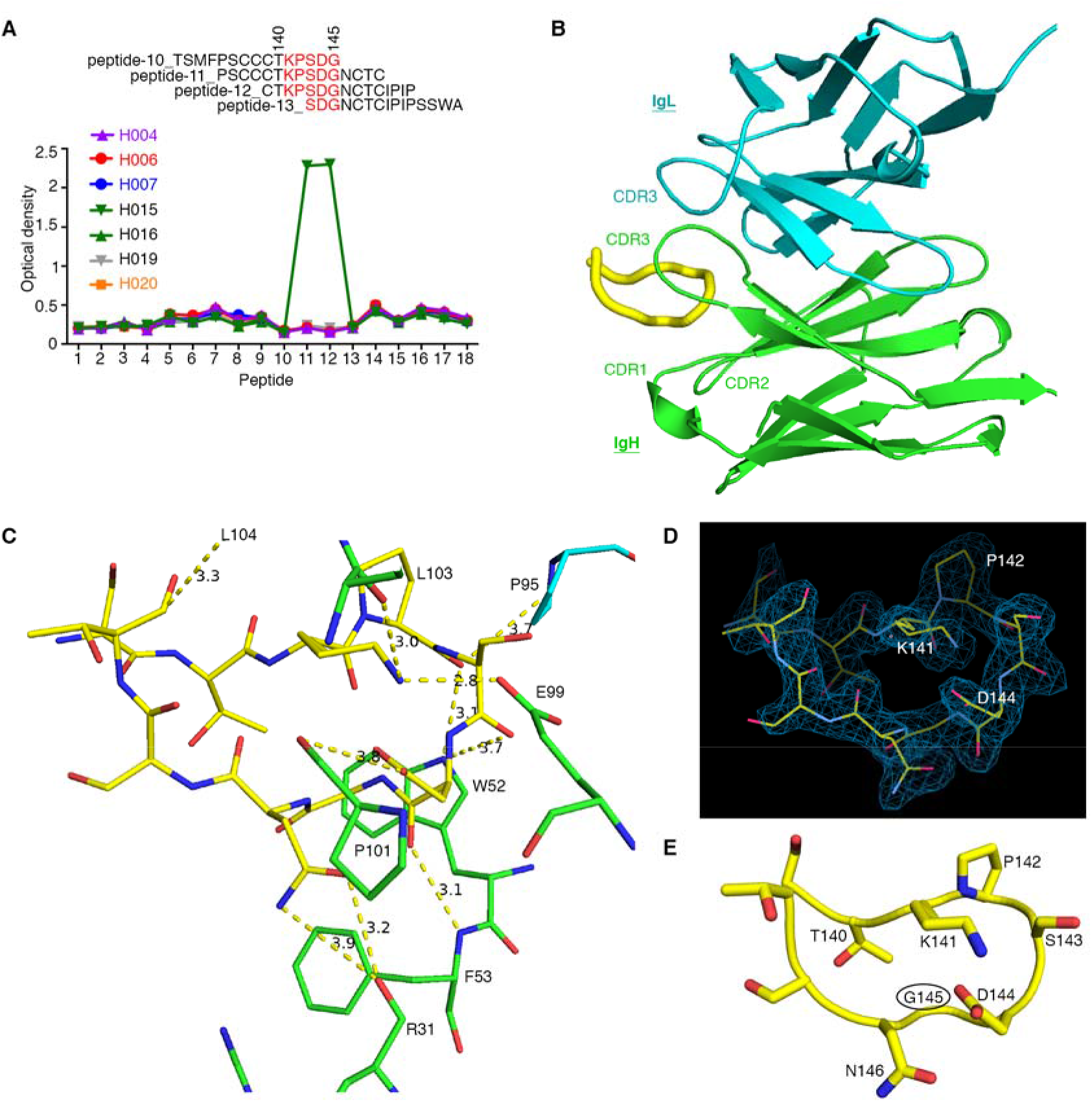
Crystal structure of H015 bound to its recognition motif. A single crystal was used to obtain a high resolution (1.78 Å) structure. (A) Synthetic peptides spanning the antigenic loop region were subjected to ELISA for antibody binding. Among the tested antibodies, only H015 binds peptides-11 and -12. Experiments were performed three times and details are in Figure S6A. (B and C) The peptide binds to CDR1 (R31), CDR2 (W52 and F53) and CDR3 (E99, P101, L103, and L104) of H015 heavy chain (green) and CDR3 (P95) of the light chain (cyan) (B). The interacting residues (C) on the heavy chain (green) are R31 (main chain), W52, F53 (main chain), E99, P101 (main chain), L103 (main chain), L104 (hydrophobic). One contact with the light chain (cyan) is with P95. (D) Electron density map of the bound peptide as seen in the 2Fo-Fc map contoured at 1 RMSD indicating high occupancy (92%). (E) The recognition motif, KPSDGN, adopts a sharp hairpin conformation due to the salt-bridge between lysine141 and aspartic acid 144 and is facilitated by kinks at P142 and G145. Glycine 145 (G145, circled) is the residue that escapes the immune system when mutated to an arginine.

To examine the molecular basis for H015 binding, its Fab fragment was co-crystallized with the target peptide epitope PSSSSTKPSDGNSTS, where all cysteine residues that flank the recognition sequence were substituted with serine to avoid non-physiological cross-linking. The 1.78 Å structure (Figure 6B and S6B) revealed that the peptide is primarily bound to the immunoglobulin heavy chain (Figure 6B and 6C), interacting with residues from CDR1 (R31), CDR2 (W52, F53) and CDR3 (E99, P101, L103, L104) of IgH with only one contact with CDR3 (P95) of IgL. The peptide adopts a sharp hairpin conformation facilitated by P142 and G145 residues that is stabilized by a salt bridge formed between K141 and D144 (Figure 6D). Interestingly, the distance between the Cαs of the two residues flanking the recognition residues is 6.4 Å, which is the distance expected if C139 and C147 form a disulfide bond in the native HBsAg structure (Ito et al., 2010).

The residues that form the hairpin are important for anti-HBs antibody recognition as determined by alanine scanning (Figure 4B and S4). Moreover, each of these residues has been identified as important for immune recognition during natural infection (Ma and Wang, 2012). G145R, the most common naturally occurring S-protein escape mutation substitutes a large positively charged residue for a small neutral residue (circled residue in Figure 6E) potentially altering the antigenic binding surface and the overall hairpin fold due to interference with the internal salt bridge between K141 and D144 and/or the propensity of glycine to loop formation.

### Protection and Therapy in Humanized Mice

HBV infection is limited to humans, chimpanzees, tree shrews, and human liver chimeric mice (Sun and Li, 2017). To determine whether our anti-HBs bNAbs prevent infection *in vivo* we produced human liver chimeric *Fah*^−/−^NOD*Rag*1^−/−^*IL2rg*^null^ (huFNRG) mice (de Jong et al., 2014) and injected them with control or H007 (Group-II) or H020 (Group-I) antibodies before infection with HBV (Figure 7A–7D). Whereas all six control animals in two independent experiments were infected, pre-exposure prophylaxis with either H007 or H020 was fully protective (Figure 7B–7D). We conclude that single anti-HBs bNAbs targeting different epitopes on the major virus surface antigen can prevent infection *in vivo*.

**Figure 7.**
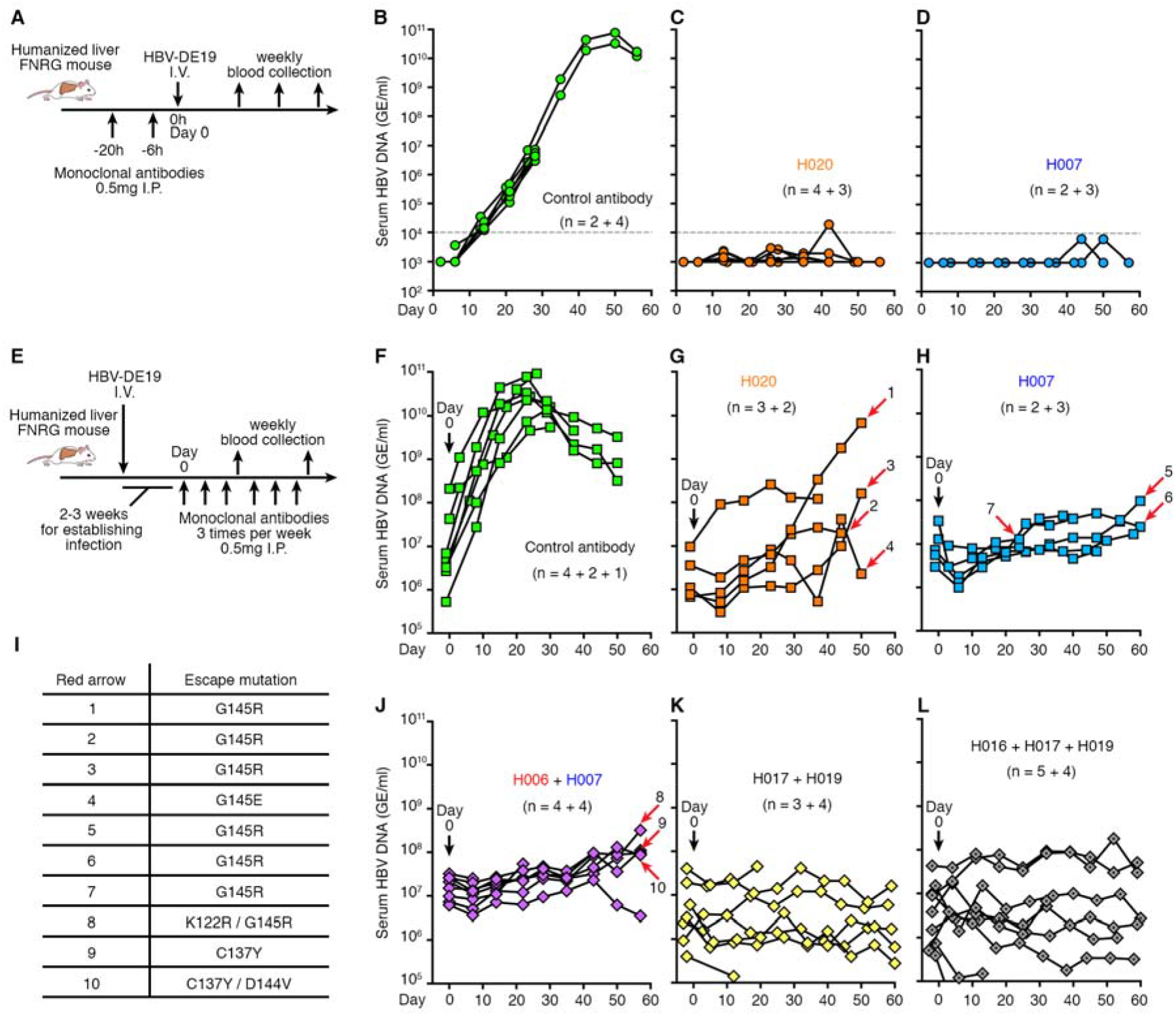
Anti-HBs bNAbs are protective and therapeutic *in vivo*. (A and E) Diagram of the prophylaxis and treatment protocols, respectively. (B) Prophylaxis with isotype control antibody 10-1074 (Mouquet et al., 2012). (C and D) Prophylaxis with H020 and H007 respectively. The dashed line in (B-D) indicates the detection limit. (F) Treatment of viremic huFNRG mice with control antibody 10-1074. (G and H) Treatment of viremic huFNRG mice with H020 alone or H007 alone, respectively. HBV DNA levels in serum were monitored on a weekly basis. Two independent experiments comprising a total of 5 to 8 mice were combined and displayed. (I) Mutations in the S-protein sequence from the indicated mice (red arrows) in (G), (H) and (J). S-protein sequence chromotograms are shown in Figure S7. (J-L) Treatment of viremic huFNRG mice with combination of anti-HBs bNAb H006 + H007 (J), or H017 + H019 (K), or H016 + H017 + H019 (L), respectively. The mice without red arrows bear no escape mutations at the last time point.

To determine whether bNAbs can also control established infections, we infused control antibody or bNAb H020 or H007 shortly after HBV-infected huFNRG mice reached ~10^7^ copies of virus per ml of serum (Figure 7E–7H and Figure S7A). Animals that received the control antibodies further increased viremia to as high as ~10^11^ DNA copies/ml (Figure 7F). In contrast, the 5 mice that received H020 maintained stable levels of viremia for around 30 days (Figure 7G), after which time 2 mice showed increased viremia (arrow-1/3 in Figure 7G). A similar result was observed in the 5 mice that received H007 (Figure 7H), where only one showed a slight increase viremia at around day 50 (arrow-5 in Figure 7H).

To determine whether the animals that showed increased HBV DNA levels during antibody monotherapy developed escape mutations, we sequenced the viral DNA recovered from mouse blood. All three mice that escaped H020 or H007 monotherapy developed viruses that carried a G145R mutation in the S-protein (arrow-1/3 in Figure 7G, arrow-5 in Figure 7H, Figure 7I, and Figure S7). This mutation represents a major immune escape mutation in humans (Zanetti et al., 1988). Furthermore, mutations at the same position in the S-protein were also identified in mice that maintained low level viremia (arrow-2/4 in Figure 7G, arrow-6/7 in Figure 7H, Figure 7I, and Figure S7), but not in control animals (Figure S7). These results show that anti-HBs bNAb monotherapy leads to the emergence of escape mutations that are consistent with bNAb binding properties *in vitro* (Figure 4C).

To determine whether a combination of bNAbs targeting 2 separate epitopes would interfere with the emergence of resistant strains, we co-administered H006 + H007 (Group-I and -II, respectively) to 8 HBV-infected huFNRG mice (Figure 7J). Similar to H007 monotherapy, there was only a slight increase in viremia in animals treated with the H006 + H007 anti-HBs bNAb combination during the 60-day observation period (Figure 7J and S7A). However, sequence analysis revealed that 3 of the mice developed resistance mutations including K122R/G145R, C137Y, and C137Y/D144V (arrow-8/9/10 in Figure 7J, Figure 7I, and Figure S7). These mutations confer loss of binding to both H006 and H007 (Figure 4C). Thus, the combination of 2 anti-HBs bNAbs targeting separate epitopes but susceptible to the same clinical escape variants is not sufficient to inhibit emergence of escape mutations.

In contrast, none of 7 mice treated with the combination of H017 + H019 (Group-III and -I, respectively) bNAbs that displayed complementary sensitivity to commonly occuring natural mutations (Figure 4C), showed increased viremia or escape mutations (Figure 7K and S7A). Similar effects were also observed in the 9 animals treated with the H016, H017 and H019 triple antibody combination (Figure 7L and S7A). Altogether, these findings suggest that control of HBV infection by bNAbs requires a combination of antibodies targeting non-overlapping groups of common escape mutations.

## DISCUSSION

Previous studies have identified several anti-HBs neutralizing antibodies from a small number of otherwise unselected spontaneously recovered or vaccinated individuals (Cerino et al., 2015; Colucci et al., 1986; Eren et al., 1998; Heijtink et al., 2002; Heijtink et al., 1995; Jin et al., 2009; Kim and Park, 2002; Li et al., 2017; Sa’adu et al., 1992; Sankhyan et al., 2016; Tajiri et al., 2007; Tokimitsu et al., 2007; Wang et al., 2016). In contrast, we screened sera from 144 exposed volunteers to identify elite neutralizers. Serologic activity varied greatly among the donors with a small number of individuals demonstrating high levels of neutralizing activity. To understand this activity, we isolated 244 anti-HBs antibodies from single B cells obtained from the top donors. Each of the elite donors tested showed expanded clones of memory B cells expressing bNAbs that targeted 3 non-overlapping sites on the S-protein. Moreover, the amino acid sequence of several of the bNAbs was highly similar in different individuals, and as might be expected these closely related antibodies target the same epitope.

The near identity of clones of HBV bNAbs in unrelated elite individuals is akin to reports for elite responders to HIV-1 (Scheid et al., 2011; West et al., 2012), influenza (Laursen and Wilson, 2013; Pappas et al., 2014; Wrammert et al., 2011), Zika (Robbiani et al., 2017), and malaria (Tan et al., 2018). However, none of the elite anti-HBs bNAbs shares both IgH and IgL with previously reported HBV neutralizing antibodies, the best of which have been tested in the clinic but are less potent than some of the bNAbs reported here (libivirumab IC_50_: 35 ng/ml, tuvirumab IC_50_: ~100 ng/ml) (Galun et al., 2002; Heijtink et al., 2001; van Nunen et al., 2001).

Our alanine scanning and competition binding analyses are consistent with the existence of at least 3 domains that can be recognized concomitantly by bNAbs (Gao et al., 2017; Tajiri et al., 2010; Zhang et al., 2016). However, the domains do not appear to be limited to either of two previously defined circular peptide epitopes, 123-137 and 139-148 (Tajiri et al., 2010; Zhang et al., 2016). Instead, residues spanning most of the external domain can contribute to binding by both Group-I and -II antibodies. For example, alanine scanning indicates that Group-I H020 binding is dependent on I110, K141, D144, G145 and T148, while Group-II H016 binding depends on T123, D144, and G145. Thus, despite having non-overlapping binding sites some of the essential residues are shared by Group-I and II suggesting that the epitopes are conformational. Moreover, the antibody epitopes on S-protein identified using mouse and human antibodies may be distinct (Chen et al., 1996; Ijaz et al., 2003; Paulij et al., 1999; Zhang et al., 2019; Zhang et al., 2016). Finally, G145, a residue that is frequently mutated in infected humans (Ma and Wang, 2012; Tong et al., 2013), is essential for binding by all the Group-II but not all Group-I or -III antibodies tested.

Although there is no high-resolution structural information available for HBsAg, crystallization of the Group-II bNAb H015 and its linear epitope revealed a loop that includes P142, S/T143, D144, and G145, all of which are frequently mutated during natural infection to produce well-documented immune escape variants (Hsu et al., 2015; Ijaz et al., 2012; Ma and Wang, 2012; Salpini et al., 2015). In addition to immune escape, the residues that form this structure are also essential for infectivity, possibly by facilitating virus interactions with cell surface glycosaminoglycans (Sureau and Salisse, 2013). Mutations in K141, P142 as well as C139 and C147, all of which contribute to the stability of the structure, decrease viral infectivity (Salisse and Sureau, 2009). We speculate that drugs that destabilize the newly elucidated H015-peptide loop structure may also interfere with infectivity.

The G145R mutation, which is among the most frequent immune escape variants, replaces a small neutral residue with a bulky charged residue that would likely interfere with antigenicity by destroying the salt bridge between K141 and D144 that anchors the peptide loop. However, this drastic structural change does not alter infectivity (Salisse and Sureau, 2009), possibly because the additional charge compensates for otherwise altered interactions between HBV and cell surface glycosaminoglycans (Sureau and Salisse, 2013). Thus, the additional charge may allow G145R to function as a dominant immune escape variant while preserving infectivity.

Our analysis is limited to antibodies directed at S-protein antigen in part because this is the antigen used in the currently FDA-approved vaccines, and because purified S-protein blocked nearly all of the neutralizing activity in the serum of the elite neutralizers irrespective of whether they were vaccinated or spontaneously recovered. Nevertheless, individuals who recover from infection naturally also produce antibodies to the PreS1 domain of HBsAg (Li et al., 2017; Sankhyan et al., 2016). The PreS1 domain is essential for the virus to interact with the entry factor NCTP on hepatocytes and potent neutralizing antibodies to PreS1 have been described (Li et al., 2017). However, these are not naturally occurring antibodies but rather randomly paired IgH and IgL chains derived from phage libraries obtained from unexposed or vaccinated healthy donors (Li et al., 2017). Moreover, the phage antibodies required further engineering to enhance their neutralizing activity (Li et al., 2017). Thus, whether the human immune system also produces potent anti-PreS1 bNAbs has yet to be determined.

Chronic HBV infection remains a major global public health problem in need of an effective curative strategy (Graber-Stiehl, 2018; Lazarus et al., 2018; Revill et al., 2016). Chronically infected individuals produce an overwhelming amount of HBsAg that is postulated to incapacitate the immune system. Consequently, immune cells, which might normally clear the virus, are unable to react to antigen, a phenomenon referred to as exhaustion or anergy (Ye et al., 2015). The appearance of anti-HBs antibodies is associated with spontaneous recovery from the disease, perhaps because they can clear the antigen and facilitate the emergence of a productive immune response (Celis and Chang, 1984; Zhang et al., 2016; Zhu et al., 2016). These findings led to the hypothesis that passively administered antibodies might be used in conjunction with antiviral drugs to further decrease the antigenic burden while enhancing immune responses that maintain long-term control of the disease. Our observations in huFNRG mice infected with HBV indicate that antibody monotherapy with a potent bNAb can lead to the emergence of the very same escape mutations commonly found in chronically infected individuals. Moreover, not all bNAb combinations are effective in preventing escape by mutation. Combinations that target separate epitopes but have overlapping sensitivity to commonly occurring escape mutations such as H006 and H007 are ineffective. In contrast, combinations with complementary sensitivity to common escape mutations prevent the emergence of escape mutations in huFNRG mice infected with HBV. Thus, immunotherapy for HBV infection may require combinations of antibodies with complementary activity to avert this potential problem.

## SUPPLEMENTAL INFORMATION

Supplemental Information includes seven figures and three tables.

## AUTHOR CONTRIBUTIONS

Q.W. conceived the study, conducted the experiments, supervised and designed the experiments, interpreted experimental results, and wrote the paper. E.M., Y.Y. and Z.W. conducted experiments, interpreted experimental results, and edited the manuscript. A.M.H. coordinated cohort studies and blood collection and edited the manuscript. D.A.O. solved and analyzed crystal structure and wrote structure-related parts of the paper. C.T.M., A.G., Y.Z., K.Y., J.P.N., J.W., Q.Z., and C.J. conducted experiments. Z.L. and Q.J. performed statistical analysis. T.O. performed computational analysis. K.C. performed single-cell sorting. T.S. and W.M.S. designed experiments and edited the manuscript. D.F.R., M.J., J.H., M.C., T.Y. and Y.W. interpreted experimental results and edited the manuscript. Y.P.J. performed mouse experiments with C.Z., M.K. and C.Q., supervised, designed, and interpreted experimental results and edited the manuscript. C.M.R. supervised, designed, and interpreted experimental results and edited the manuscript. M.C.N. conceived the study, supervised, designed, and interpreted experiments and wrote the paper.

## ACKNOWLEDGMENTS

We would especially like to thank all the volunteers who agreed to participate in this study, as well as the staff of the Rockefeller Hospital and Clinical Lab for their assistance with blood collection, processing, and storage. We thank members of the Nussenzweig, Rice, and Wen labs for helpful discussion and suggestions. A particular thank you to Rui Gong, Yixiao Zhang, Xiaochun Li, Kai Liu, Amelia Escolano and Irina Shimeliovich for sharing expertise, protocols, and reagents. Peptide synthesis was performed by Henry Zebroski at the Proteomics Resource Center of The Rockefeller University. This work was supported by National Natural Science Foundation of China 31872730 (to Q.W.), the Thousand Talents Plan Youth Program (to Q.W.), and The Program for Professor of Special Appointment (Eastern Scholar) at Shanghai Institutions of Higher Learning TP2017010 (to Q.W.). The project was also supported in part by grant #UL1TR001866 (to Q.W.) from the National Center for Advancing Translational Sciences (NCATS), National Institutes of Health (NIH) Clinical and Translational Science Award (CTSA) program; NIH grants R01AI143295 and R01DK085713 (to C.M.R.), 2U19AI111825-06 (to MCN), K08DK090576 (to Y.P.J.), NIH fellowship F32DK107164 (to E.M.). The project was co-sponsored by the Center for Basic and Translational Research on Disorders of the Digestive System through the generosity of the Leona M. and Harry B. Helmsley Charitable trust (to E.M.). The use of the instruments in the Rockefeller University Structural Biology Resource Center was made possible by grant number 1S10RR027037-01 from the National Center for Research Resources of the NIH. We thank the staff at APS for their support of remote data collection. This work is based upon research conducted at the Northeastern Collaborative Access Team beamlines, which are funded by the National Institute of General Medical Sciences from NIH (P30 GM124165). The Eiger 16M detector on 24-ID-E beam line is funded by a NIH-ORIP HEI grant (S10OD021527). This research used resources of the Advanced Photon Source, a U.S. Department of Energy (DOE) Office of Science User Facility operated for the DOE Office of Science by Argonne National Laboratory under Contract No. DE-AC02-06CH11357. Support was also provided by the Robertson Therapeutic Development Fund (to Q.W., E.M. and M.C.N.). M.C.N. is an HHMI Investigator. In connection with this work, Q.W. and M.C.N. have a provisional patent application with the U.S. Patent and Trademark Office (62898735). The content is solely the responsibility of the authors and does not necessarily represent the official views of any of the funding agencies or individuals.

