## Supplemental Table 1 for "Public broadly neutralizing antibodies against hepatitis B virus in individuals with elite serologic activity"

Table S1. Detailed information of donors including their anti-HBs ELISA titer (x-axis in Figure 1A and S1B) and relative infection rate determined by *in vitro* neutralization assay (y-axis in Figure 1A and S1B).

| **DONOR ID** | **ANTI-HBs ELISA TITER** | **STATUE** | **BIG BLOOD DRAW** | **INFECTION RATE (1:5 SERUM) PERCENTAGE OF HBcAg+ CELLS** | | | **AVERAGE** | **NEUTRA CAPACITY** | **INFECTION RATE (1:50 SERUM) PERCENTAGE OF HBcAg+ CELLS** | | | **AVERAGE** | **NEUTRA CAPACITY** |
| --- | --- | --- | --- | --- | --- | --- | --- | --- | --- | --- | --- | --- | --- |
| 1 | 9.08 | Vaccinated |  | 114.00 | 127.12 | 133.64 | 124.92 | 0.80 | 118.06 | 128.90 | 122.19 | 123.05 | 0.81 |
| 2 | 516.12 | Vaccinated |  | 65.28 | 77.58 | 75.13 | 72.66 | 1.38 | 131.60 | 137.04 | 133.44 | 134.03 | 0.75 |
| 3 | 0 | Vaccinated |  | 108.59 | 118.31 | 118.53 | 115.14 | 0.87 | 132.81 | 136.65 | 132.22 | 133.89 | 0.75 |
| 4 | 267.1 | Vaccinated |  | 113.39 | 121.91 | 121.67 | 118.99 | 0.84 | 142.69 | 146.69 | 153.24 | 147.54 | 0.68 |
| 5 | 113.01 | Vaccinated |  | 82.91 | 87.48 | 95.39 | 88.59 | 1.13 | 148.24 | 149.36 | 146.75 | 148.12 | 0.68 |
| 6 | 325.77 | Vaccinated |  | 81.19 | 85.97 | 96.43 | 87.86 | 1.14 | 117.48 | 116.83 | 119.74 | 118.02 | 0.85 |
| 7 | 63.54 | Vaccinated |  | 100.37 | 105.53 | 110.73 | 105.54 | 0.95 | 134.11 | 135.15 | 138.51 | 135.92 | 0.74 |
| 8 | 21.8 | Vaccinated |  | 40.86 | 35.60 | 42.82 | 39.76 | 2.52 | 78.64 | 84.09 | 84.79 | 82.51 | 1.21 |
| 9 | >1000 | Vaccinated | BIG BLOOD DRAW | 12.84 | 14.39 | 17.20 | 14.81 | 6.75 | 34.66 | 38.81 | 33.48 | 35.65 | 2.81 |
| 10 | 0.33 | Vaccinated |  | 93.33 | 93.42 | 104.91 | 97.22 | 1.03 | 105.41 | 116.72 | 115.58 | 112.57 | 0.89 |
| 11 | 1.29 | Vaccinated | BIG BLOOD DRAW | 109.40 | 113.87 | 94.10 | 105.79 | 0.95 | 95.79 | 93.25 | 81.95 | 90.33 | 1.11 |
| 12 | 12.87 | Vaccinated |  | 102.04 | 102.94 | 108.81 | 104.60 | 0.96 | 142.95 | 137.95 | 125.12 | 135.34 | 0.74 |
| 13 | >1000 | Vaccinated | BIG BLOOD DRAW | 9.24 | 8.31 | 8.33 | 8.63 | 11.59 | 19.95 | 20.81 | 17.29 | 19.35 | 5.17 |
| 14 | 248.31 | Vaccinated |  | 80.90 | 80.65 | 79.88 | 80.48 | 1.24 | 117.58 | 120.17 | 115.37 | 117.71 | 0.85 |
| 15 | 76.1 | Vaccinated |  | 97.26 | 91.48 | 86.95 | 91.90 | 1.09 | 112.79 | 109.82 | 101.19 | 107.93 | 0.93 |
| 16 | 0.36 | Non-Vaccinated |  | 100.22 | 96.77 | 95.84 | 97.61 | 1.02 | 118.04 | 120.05 | 117.34 | 118.48 | 0.84 |
| 17 | 0.63 | Vaccinated |  | 100.30 | 102.98 | 101.55 | 101.61 | 0.98 | 122.27 | 119.42 | 110.93 | 117.54 | 0.85 |
| 18 | 3.24 | Vaccinated |  | 101.65 | 113.38 | 96.98 | 104.00 | 0.96 | 118.18 | 121.26 | 107.05 | 115.50 | 0.87 |
| 19 | >1000 | Vaccinated |  | 47.62 | 43.35 | 40.11 | 43.69 | 2.29 | 105.86 | 100.46 | 89.54 | 98.62 | 1.01 |
| 20 | 1.21 | Non-Vaccinated |  | 77.13 | 75.25 | 60.04 | 70.81 | 1.41 | 116.28 | 113.72 | 108.65 | 112.88 | 0.89 |
| 21 | 15.81 | Vaccinated |  | 33.85 | 41.39 | 34.98 | 36.74 | 2.72 | 89.74 | 89.87 | 88.60 | 89.40 | 1.12 |
| 22 | 2.17 | Vaccinated |  | 83.52 | 94.22 | 93.88 | 90.54 | 1.10 | 100.08 | 119.88 | 110.19 | 110.05 | 0.91 |
| 23 | 183.37 | Vaccinated |  | 93.15 | 119.57 | 116.08 | 109.60 | 0.91 | 113.43 | 117.91 | 103.45 | 111.60 | 0.90 |
| 24 | 256.85 | Vaccinated |  | 55.70 | 60.03 | 49.13 | 54.95 | 1.82 | 104.68 | 116.21 | 105.17 | 108.69 | 0.92 |
| 25 | 709.29 | Vaccinated |  | 76.18 | 77.38 | 75.01 | 76.19 | 1.31 | 100.58 | 105.57 | 116.82 | 107.66 | 0.93 |
| 26 | 0.49 | Non-Vaccinated |  | 83.02 | 102.42 | 96.98 | 94.14 | 1.06 | 104.24 | 117.09 | 105.73 | 109.02 | 0.92 |
| 27 | 0.3 | Non-Vaccinated | BIG BLOOD DRAW | 103.51 | 128.79 | 114.82 | 115.71 | 0.86 | 105.91 | 126.68 | 117.03 | 116.54 | 0.86 |
| 28 | 6.51 | Non-Vaccinated |  | 84.03 | 90.88 | 87.77 | 87.56 | 1.14 | 107.23 | 117.89 | 108.21 | 111.11 | 0.90 |
| 29 | 1.67 | Non-Vaccinated |  | 103.72 | 98.68 | 83.74 | 95.38 | 1.05 | 96.06 | 102.25 | 100.20 | 99.50 | 1.00 |
| 30 | 0.09 | Vaccinated | BIG BLOOD DRAW | 86.06 | 84.44 | 91.73 | 87.41 | 1.14 | 111.74 | 115.73 | 102.26 | 109.91 | 0.91 |
| 31 | 488.3 | Core Ab + |  | 71.30 | 66.21 | 56.71 | 64.74 | 1.54 | 94.99 | 102.87 | 94.27 | 97.38 | 1.03 |
| 32 | >1000 | Vaccinated |  | 71.12 | 67.87 | 56.53 | 65.17 | 1.53 | 97.45 | 107.81 | 106.59 | 103.95 | 0.96 |
| 33 | 330.93 | Vaccinated |  | 99.61 | 93.60 | 87.25 | 93.49 | 1.07 | 106.28 | 116.65 | 108.45 | 110.46 | 0.91 |
| 34 | 588.26 | Core Ab + |  | 40.92 | 41.19 | 35.30 | 39.14 | 2.56 | 99.54 | 111.16 | 105.28 | 105.32 | 0.95 |
| 35 | 365.7 | Vaccinated |  | 101.52 | 113.83 | 94.76 | 103.37 | 0.97 | 129.77 | 131.60 | 121.19 | 127.52 | 0.78 |
| 36 | 108.83 | Vaccinated |  | 87.55 | 85.69 | 77.65 | 83.63 | 1.20 | 122.28 | 122.03 | 107.69 | 117.33 | 0.85 |
| 37 | >1000 | Vaccinated |  | 23.79 | 21.09 | 18.46 | 21.11 | 4.74 | 66.02 | 59.97 | 52.64 | 59.54 | 1.68 |
| 38 | 11.76 | Vaccinated |  | 94.89 | 95.53 | 96.87 | 95.76 | 1.04 | 119.79 | 125.09 | 115.27 | 120.05 | 0.83 |
| 39 | 230.99 | Vaccinated |  | 102.08 | 99.23 | 91.54 | 97.62 | 1.02 | 113.01 | 115.04 | 98.95 | 109.00 | 0.92 |
| 40 | 0.26 | Vaccinated |  | 54.12 | 57.38 | 45.32 | 52.27 | 1.91 | 102.43 | 107.89 | 95.37 | 101.90 | 0.98 |
| 41 | 0.42 | Non-Vaccinated |  | 77.09 | 78.29 | 84.91 | 80.09 | 1.25 | 87.59 | 101.25 | 101.19 | 96.68 | 1.03 |
| 42 | 128.75 | Core Ab + |  | 97.03 | 106.51 | 110.26 | 104.60 | 0.96 | 114.07 | 149.01 | 121.98 | 128.35 | 0.78 |
| 43 | 229.75 | Core Ab + |  | 78.56 | 94.35 | 95.71 | 89.54 | 1.12 | 95.18 | 144.23 | 123.82 | 121.08 | 0.83 |
| 44 | 222.48 | Core Ab + |  | 45.85 | 56.61 | 65.34 | 55.93 | 1.79 | 94.00 | 133.96 | 109.86 | 112.60 | 0.89 |
| 45 | >1000 | Core Ab + |  | 51.82 | 66.02 | 61.55 | 59.80 | 1.67 | 104.36 | 122.03 | 120.96 | 115.78 | 0.86 |
| 46 | 350.93 | Vaccinated |  | 89.57 | 99.40 | 111.55 | 100.17 | 1.00 | 100.98 | 129.50 | 111.79 | 114.09 | 0.88 |
| 47 | 0.33 | Vaccinated |  | 126.88 | 138.30 | 142.37 | 135.85 | 0.74 | 112.35 | 156.77 | 127.72 | 132.28 | 0.76 |
| 48 | >1000 | Vaccinated | BIG BLOOD DRAW | 35.94 | 44.44 | 46.02 | 42.14 | 2.37 | 83.08 | 109.89 | 94.60 | 95.86 | 1.04 |
| 49 | >1000 | Core Ab + | BIG BLOOD DRAW | 94.21 | 120.65 | 119.57 | 111.48 | 0.90 | 113.33 | 139.10 | 126.42 | 126.28 | 0.79 |
| 50 | 6.2 | Vaccinated |  | 93.10 | 114.26 | 113.58 | 106.98 | 0.93 | 93.34 | 121.07 | 112.91 | 109.10 | 0.92 |
| 51 | 228.35 | Vaccinated |  | 111.78 | 112.69 | 115.97 | 113.48 | 0.88 | 104.71 | 102.64 | 83.60 | 96.98 | 1.03 |
| 52 | 2.77 | Vaccinated |  | 116.53 | 115.59 | 110.83 | 114.32 | 0.87 | 113.48 | 114.25 | 125.60 | 117.78 | 0.85 |
| 53 | 48.25 | Vaccinated |  | 114.90 | 112.78 | 98.40 | 108.69 | 0.92 | 105.04 | 116.61 | 119.01 | 113.55 | 0.88 |
| 54 | 24.06 | Vaccinated |  | 121.56 | 125.13 | 111.71 | 119.47 | 0.84 | 99.64 | 118.99 | 122.11 | 113.58 | 0.88 |
| 55 | 772.43 | Core Ab + | BIG BLOOD DRAW | 8.79 | 9.33 | 5.78 | 7.97 | 12.55 | 82.33 | 95.93 | 87.80 | 88.69 | 1.13 |
| 56 | 441.12 | Vaccinated |  | 86.57 | 86.67 | 79.10 | 84.12 | 1.19 | 98.81 | 120.31 | 116.13 | 111.75 | 0.89 |
| 57 | 91.37 | Vaccinated |  | 120.46 | 113.18 | 110.80 | 114.82 | 0.87 | 111.45 | 107.08 | 116.80 | 111.78 | 0.89 |
| 58 | >1000 | Vaccinated |  | 27.66 | 27.74 | 26.37 | 27.26 | 3.67 | 82.86 | 83.73 | 74.74 | 80.44 | 1.24 |
| 59 | 83.28 | Vaccinated |  | 110.69 | 109.95 | 99.34 | 106.66 | 0.94 | 115.21 | 115.69 | 99.96 | 110.29 | 0.91 |
| 60 | >1000 | Vaccinated | BIG BLOOD DRAW | 5.68 | 6.72 | 5.17 | 5.86 | 17.08 | 25.39 | 26.65 | 24.93 | 25.66 | 3.90 |
| 61 | 104.78 | Vaccinated |  | 109.98 | 120.25 | 131.62 | 120.61 | 0.83 | 107.59 | 134.01 | 124.63 | 122.08 | 0.82 |
| 62 | 2.56 | Vaccinated |  | 113.16 | 140.19 | 143.78 | 132.38 | 0.76 | 120.58 | 139.98 | 135.32 | 131.96 | 0.76 |
| 63 | 246.16 | Vaccinated |  | 101.93 | 112.01 | 141.39 | 118.44 | 0.84 | 114.71 | 134.13 | 143.18 | 130.67 | 0.77 |
| 64 | Not Available | Vaccinated |  | 128.17 | 114.29 | 147.09 | 129.85 | 0.77 | 133.50 | 133.31 | 133.23 | 133.35 | 0.75 |
| 65 | 0.28 | Core Ab + |  | 112.39 | 133.12 | 155.43 | 133.65 | 0.75 | 131.76 | 134.78 | 150.74 | 139.09 | 0.72 |
| 66 | 265.6 | Vaccinated |  | 50.87 | 68.37 | 86.59 | 68.61 | 1.46 | 129.24 | 133.43 | 119.16 | 127.28 | 0.79 |
| 67 | 0 | Vaccinated |  | 106.36 | 120.57 | 150.56 | 125.83 | 0.79 | 122.57 | 142.42 | 139.51 | 134.84 | 0.74 |
| 68 | 0.25 | Vaccinated |  | 58.26 | 87.26 | 105.25 | 83.59 | 1.20 | 123.65 | 128.77 | 124.58 | 125.67 | 0.80 |
| 69 | >1000 | Vaccinated | BIG BLOOD DRAW | 11.42 | 27.20 | 22.32 | 20.31 | 4.92 | 50.80 | 55.44 | 64.27 | 56.84 | 1.76 |
| 70 | 7.01 | Vaccinated |  | 107.39 | 113.96 | 138.16 | 119.84 | 0.83 | 118.06 | 116.97 | 118.95 | 117.99 | 0.85 |
| 71 | 638.24 | Vaccinated |  | 56.39 | 48.55 | 55.42 | 53.45 | 1.87 | 94.83 | 111.19 | 117.23 | 107.75 | 0.93 |
| 72 | 15.55 | Vaccinated |  | 128.74 | 135.08 | 126.34 | 130.05 | 0.77 | 137.00 | 132.93 | 122.94 | 130.96 | 0.76 |
| 73 | 225.34 | Vaccinated |  | 131.87 | 122.33 | 137.50 | 130.57 | 0.77 | 137.43 | 128.35 | 124.59 | 130.12 | 0.77 |
| 74 | 159.52 | Core Ab + |  | 120.51 | 119.04 | 128.75 | 122.76 | 0.81 | 140.98 | 130.18 | 124.00 | 131.72 | 0.76 |
| 75 | 0.37 | Vaccinated |  | 131.78 | 125.05 | 159.51 | 138.78 | 0.72 | 133.08 | 135.29 | 117.74 | 128.70 | 0.78 |
| 76 | 27.37 | Vaccinated |  | 137.32 | 145.89 |  | 141.60 | 0.71 | 132.06 |  |  | 132.06 | 0.76 |
| 77 | 52.77 | Vaccinated |  | 112.72 | 121.48 |  | 117.10 | 0.85 | 137.12 |  |  | 137.12 | 0.73 |
| 78 | 181.6 | Vaccinated |  | 116.16 | 107.29 |  | 111.72 | 0.90 | 138.30 | 131.27 |  | 134.79 | 0.74 |
| 79 | >1000 | Vaccinated |  | 54.73 | 59.28 | 71.14 | 61.72 | 1.62 | 130.53 | 128.66 | 118.32 | 125.84 | 0.79 |
| 80 | >1000 | Vaccinated |  | 62.38 | 53.55 | 69.34 | 61.76 | 1.62 | 122.37 | 106.42 |  | 114.40 | 0.87 |
| 81 | 7.23 | Vaccinated |  | 113.81 | 111.55 | 142.59 | 122.65 | 0.82 | 109.38 | 105.63 | 116.78 | 110.60 | 0.90 |
| 82 | 24.98 | Vaccinated |  | 120.69 | 124.58 | 118.10 | 121.12 | 0.83 | 106.45 | 118.55 | 117.79 | 114.26 | 0.88 |
| 83 | 90.5 | Vaccinated |  | 84.79 | 87.62 | 99.65 | 90.69 | 1.10 | 91.86 | 99.93 | 105.67 | 99.15 | 1.01 |
| 84 | 880.92 | Core Ab + |  | 112.81 | 125.10 | 114.29 | 117.40 | 0.85 | 106.78 | 117.78 | 147.46 | 124.01 | 0.81 |
| 85 | 82.25 | Vaccinated |  | 120.45 | 148.83 | 159.96 | 143.08 | 0.70 | 114.14 | 114.33 | 136.45 | 121.64 | 0.82 |
| 86 | 19.16 | Vaccinated |  | 147.29 | 173.59 | 166.75 | 162.55 | 0.62 | 110.03 | 128.28 | 120.46 | 119.59 | 0.84 |
| 87 | 88.6 | Vaccinated |  | 126.58 | 146.46 | 145.74 | 139.59 | 0.72 | 105.86 | 119.12 | 119.22 | 114.74 | 0.87 |
| 88 | 288.42 | Core Ab + |  | 108.25 | 120.39 | 140.09 | 122.91 | 0.81 | 90.55 | 101.28 | 115.15 | 102.33 | 0.98 |
| 89 | 15.74 | Vaccinated |  | 101.12 | 94.87 | 98.43 | 98.14 | 1.02 | 75.70 | 96.65 | 111.21 | 94.52 | 1.06 |
| 90 | >1000 | Vaccinated |  | 54.59 | 47.73 | 53.63 | 51.99 | 1.92 | 71.09 | 87.31 | 99.22 | 85.87 | 1.16 |
| 91 | >1000 | Vaccinated |  | 18.28 | 23.05 | 19.85 | 20.39 | 4.90 | 48.17 | 59.50 | 53.15 | 53.61 | 1.87 |
| 93 | 2.64 | Vaccinated |  | 155.82 | 147.87 | 143.25 | 148.98 | 0.67 | 126.83 | 113.67 | 124.04 | 121.51 | 0.82 |
| 94 | 127.58 | Vaccinated |  | 160.76 | 145.63 | 119.61 | 142.00 | 0.70 | 105.84 | 110.91 | 114.14 | 110.30 | 0.91 |
| 95 | 23.74 | Vaccinated |  | 176.74 | 160.65 | 159.27 | 165.55 | 0.60 | 110.83 | 103.36 | 112.17 | 108.78 | 0.92 |
| 96 | >1000 | Vaccinated |  | 51.24 | 50.60 | 36.36 | 46.07 | 2.17 | 109.95 | 99.89 | 100.76 | 103.54 | 0.97 |
| 97 | 25.97 | Vaccinated |  | 151.86 | 144.37 | 127.26 | 141.16 | 0.71 | 109.63 | 109.86 | 98.50 | 106.00 | 0.94 |
| 98 | >1000 | Vaccinated |  | 73.82 | 83.70 | 70.82 | 76.11 | 1.31 | 88.55 | 95.98 | 100.28 | 94.94 | 1.05 |
| 99 | 688.28 | Vaccinated | BIG BLOOD DRAW | 110.48 | 112.47 | 90.05 | 104.33 | 0.96 | 100.29 | 103.59 | 94.65 | 99.51 | 1.00 |
| 100 | 366.05 | Vaccinated |  | 85.68 | 99.38 | 86.23 | 90.43 | 1.11 | 97.85 | 101.09 | 85.62 | 94.85 | 1.05 |
| 101 | 1.22 | Vaccinated |  | 88.54 | 96.48 | 92.77 | 92.60 | 1.08 | 107.51 | 104.47 | 106.69 | 106.22 | 0.94 |
| 102 | 0.65 | Vaccinated |  | 96.90 | 94.97 | 99.88 | 97.25 | 1.03 | 109.63 | 101.73 | 102.43 | 104.60 | 0.96 |
| 103 | 45.53 | Core Ab + |  | 100.38 | 104.13 | 93.37 | 99.29 | 1.01 | 103.40 | 95.77 | 93.83 | 97.67 | 1.02 |
| 104 | 95.76 | Vaccinated |  | 95.99 | 87.67 | 79.38 | 87.68 | 1.14 | 111.24 | 105.95 | 113.01 | 110.07 | 0.91 |
| 105 | 1.48 | Vaccinated |  | 42.47 | 43.05 | 47.61 | 44.38 | 2.25 | 102.15 | 99.45 | 98.24 | 99.95 | 1.00 |
| 106 | >1000 | Vaccinated |  | 16.40 | 16.66 | 16.15 | 16.40 | 6.10 | 64.77 | 68.47 | 66.59 | 66.61 | 1.50 |
| 107 | 32.84 | Vaccinated |  | 148.16 | 141.65 | 127.35 | 139.05 | 0.72 | 123.79 | 119.06 | 104.81 | 115.89 | 0.86 |
| 108 | 352.99 | Vaccinated |  | 78.23 | 81.50 | 84.50 | 81.41 | 1.23 | 109.31 | 116.46 | 103.38 | 109.72 | 0.91 |
| 109 | 15.04 | Vaccinated |  | 112.45 | 103.15 | 97.70 | 104.43 | 0.96 | 110.85 | 121.08 | 117.28 | 116.40 | 0.86 |
| 110 | 26.27 | Vaccinated |  | 86.31 | 102.24 | 99.68 | 96.08 | 1.04 | 115.57 | 130.13 | 116.61 | 120.77 | 0.83 |
| 111 | 40.79 | Vaccinated |  | 81.44 | 86.98 | 87.80 | 85.40 | 1.17 | 109.93 | 98.54 | 91.77 | 100.08 | 1.00 |
| 112 | 260.82 | Core Ab + |  | 35.62 | 36.63 | 32.67 | 34.97 | 2.86 | 69.44 | 65.66 | 65.06 | 66.72 | 1.50 |
| 113 | 1.97 | Vaccinated |  | 93.25 | 98.77 | 81.76 | 91.26 | 1.10 | 101.75 | 104.27 | 100.12 | 102.05 | 0.98 |
| 114 | >1000 | Vaccinated |  | 32.49 | 27.64 | 25.33 | 28.49 | 3.51 | 80.93 | 61.56 | 71.59 | 71.36 | 1.40 |
| 115 | 798.12 | Core Ab + |  | 13.86 | 14.67 | 15.02 | 14.52 | 6.89 | 45.62 | 46.82 | 42.16 | 44.87 | 2.23 |
| 116 | 459.84 | Vaccinated |  | 99.96 | 93.67 | 81.32 | 91.65 | 1.09 | 107.55 | 108.08 | 109.31 | 108.31 | 0.92 |
| 117 | 0.46 | Vaccinated |  | 98.06 | 92.72 | 95.42 | 95.40 | 1.05 | 106.12 | 104.66 | 111.74 | 107.51 | 0.93 |
| 118 | 20.87 | Vaccinated |  | 151.82 | 146.04 | 144.65 | 147.50 | 0.68 | 112.36 | 109.15 | 124.29 | 115.26 | 0.87 |
| 119 | 91.47 | Vaccinated |  | 114.36 | 109.63 | 108.87 | 110.96 | 0.90 | 108.91 | 112.79 | 104.89 | 108.86 | 0.92 |
| 120 | 13 | Core Ab + |  | 44.97 | 45.06 | 40.30 | 43.44 | 2.30 | 102.35 | 110.61 | 97.70 | 103.55 | 0.97 |
| 121 | 175.21 | Vaccinated |  | 91.44 | 91.91 | 96.95 | 93.43 | 1.07 | 166.31 | 164.27 | 144.97 | 158.52 | 0.63 |
| 122 | 51.51 | Vaccinated |  | 95.47 | 92.45 | 100.97 | 96.30 | 1.04 | 180.79 | 174.92 | 143.73 | 166.48 | 0.60 |
| 123 | 36.81 | Vaccinated |  | 145.77 | 156.75 | 146.88 | 149.80 | 0.67 | 180.84 | 176.71 | 135.18 | 164.24 | 0.61 |
| 124 | 1.3 | Vaccinated |  | 147.11 | 133.31 | 131.87 | 137.43 | 0.73 | 170.89 | 169.84 | 127.57 | 156.10 | 0.64 |
| 125 | 1.44 | Vaccinated |  | 110.99 | 111.85 | 118.19 | 113.68 | 0.88 | 178.09 | 178.48 | 149.45 | 168.67 | 0.59 |
| 126 | 359.06 | Core Ab + |  | 79.16 | 76.31 | 85.67 | 80.38 | 1.24 | 167.98 | 161.85 | 154.57 | 161.47 | 0.62 |
| 127 | 1.23 | Non-Vaccinated |  | 128.75 | 125.40 | 143.54 | 132.56 | 0.75 | 162.01 | 157.22 | 139.08 | 152.77 | 0.65 |
| 128 | >1000 | Vaccinated |  | 30.63 | 25.86 | 36.31 | 30.94 | 3.23 | 69.73 | 82.09 | 84.44 | 78.76 | 1.27 |
| 129 | 0.17 | Non-Vaccinated |  | 166.56 | 156.86 | 157.89 | 160.44 | 0.62 | 162.24 | 168.18 | 154.52 | 161.65 | 0.62 |
| 131 | 11.81 | Vaccinated |  | 143.14 | 129.67 | 114.98 | 129.26 | 0.77 | 156.94 | 144.37 | 119.26 | 140.19 | 0.71 |
| 132 | 0 | Non-Vaccinated |  | 143.37 | 148.51 | 134.41 | 142.09 | 0.70 | 170.72 | 146.03 | 126.56 | 147.77 | 0.68 |
| 133 | 731.27 | Core Ab + |  | 86.48 | 85.37 | 78.16 | 83.34 | 1.20 | 157.64 | 137.87 | 131.07 | 142.20 | 0.70 |
| 134 | >1000 | Vaccinated |  | 31.62 | 30.52 | 27.60 | 29.92 | 3.34 | 144.36 | 137.82 | 115.14 | 132.44 | 0.76 |
| 135 | 796.93 | Core Ab + |  | 81.56 | 81.84 | 88.25 | 83.88 | 1.19 | 157.58 | 142.42 | 116.84 | 138.95 | 0.72 |
| 136 | 0 | Non-Vaccinated |  | 128.19 | 132.95 | 139.91 | 133.68 | 0.75 | 165.85 | 140.56 | 137.58 | 148.00 | 0.68 |
| 137 | 1.56 | Non-Vaccinated |  | 155.09 | 151.93 | 137.05 | 148.02 | 0.68 | 175.51 | 155.22 | 128.75 | 153.16 | 0.65 |
| 138 | 0.74 | Non-Vaccinated |  | 160.76 | 159.87 | 135.61 | 152.08 | 0.66 | 165.30 | 137.96 | 119.52 | 140.93 | 0.71 |
| 139 | 0 | Non-Vaccinated |  | 143.96 | 145.71 | 123.98 | 137.88 | 0.73 | 143.82 | 125.60 | 114.22 | 127.88 | 0.78 |
| 140 | 609.46 | Core Ab + |  | 93.61 | 96.21 | 89.99 | 93.27 | 1.07 | 125.42 | 135.92 | 106.58 | 122.64 | 0.82 |
| 141 | >1000 | Vaccinated |  | 35.14 | 37.74 | 36.31 | 36.40 | 2.75 | 90.32 | 87.71 | 83.80 | 87.28 | 1.15 |
| 142 | 0.01 | Non-Vaccinated |  | 72.39 | 72.79 | 69.24 | 71.47 | 1.40 | 85.86 | 89.49 | 80.98 | 85.44 | 1.17 |
| 143 | 51.9 | Vaccinated |  | 83.85 | 74.96 | 78.42 | 79.07 | 1.26 | 89.70 | 83.02 | 81.24 | 84.65 | 1.18 |
| 144 | 192.41 | Vaccinated |  | 99.03 | 103.75 | 104.69 | 102.49 | 0.98 | 88.00 | 80.96 | 100.21 | 89.73 | 1.11 |
| 145 | >1000 | Vaccinated |  | 35.23 | 36.75 | 36.99 | 36.32 | 2.75 | 91.40 | 83.04 | 84.37 | 86.27 | 1.16 |
| 146 | >1000 | Vaccinated | BIG BLOOD DRAW | 3.21 | 3.17 | 3.47 | 3.28 | 30.45 | 4.66 | 4.45 | 3.82 | 4.31 | 23.18 |
| 147 | >1000 | Vaccinated |  | 74.42 | 79.63 | 70.02 | 74.69 | 1.34 | 96.43 | 89.82 | 90.93 | 92.39 | 1.08 |
| 149 | >1000 | Vaccinated |  | 48.09 | 53.38 | 53.47 | 51.65 | 1.94 | 85.58 | 89.74 | 84.49 | 86.60 | 1.15 |
| 150 | >1000 | Vaccinated |  | 45.84 | 50.91 | 49.78 | 48.84 | 2.05 | 97.27 | 93.55 | 92.90 | 94.57 | 1.06 |
| 151 | 24.03 | Vaccinated |  | 76.16 | 78.93 | 83.53 | 79.54 | 1.26 | 104.49 | 103.12 | 97.68 | 101.76 | 0.98 |
| 152 | 32.89 | Vaccinated |  | 76.50 | 74.31 | 58.42 | 69.75 | 1.43 | 80.82 | 81.72 | 78.43 | 80.33 | 1.24 |
| 153 | 0.36 | Vaccinated |  | 92.38 | 95.06 | 83.56 | 90.33 | 1.11 | 81.13 | 82.29 | 70.79 | 78.07 | 1.28 |
| 154 | 8.02 | Vaccinated |  | 81.20 | 78.04 | 61.49 | 73.58 | 1.36 | 89.89 | 80.29 | 72.54 | 80.90 | 1.24 |
| 155 | 271.68 | Vaccinated |  | 83.98 | 86.99 | 69.06 | 80.01 | 1.25 | 79.14 | 80.93 | 74.47 | 78.18 | 1.28 |
| 156 | >1000 | Vaccinated |  | 23.65 | 20.82 | 15.97 | 20.15 | 4.96 | 78.25 | 75.10 | 58.66 | 70.67 | 1.42 |
| 157 | 93.05 | Vaccinated |  | 93.75 | 95.02 | 83.23 | 90.67 | 1.10 | 77.02 | 80.62 | 70.27 | 75.97 | 1.32 |
| 158 | 261.09 | Vaccinated |  | 78.75 | 78.50 | 64.50 | 73.92 | 1.35 | 88.22 | 86.79 | 71.09 | 82.03 | 1.22 |
| 159 | 61.12 | Vaccinated |  | 87.14 | 91.02 | 76.00 | 84.72 | 1.18 | 80.77 | 86.19 | 74.40 | 80.45 | 1.24 |
| 160 | 381.69 | Vaccinated |  | 72.70 | 68.79 | 62.46 | 67.98 | 1.47 | 81.13 | 74.64 | 66.11 | 73.96 | 1.35 |
| 161 | 3.62 | Vaccinated |  | 91.93 | 85.42 | 72.87 | 83.40 | 1.20 | 85.76 | 80.35 | 74.62 | 80.24 | 1.25 |
| 162 | 198.58 | Vaccinated |  | 91.35 | 92.25 | 79.20 | 87.60 | 1.14 | 112.63 | 106.14 | 95.29 | 104.69 | 0.96 |
