## Supplemental Table 2 for "Public broadly neutralizing antibodies against hepatitis B virus in individuals with elite serologic activity"

Table S2. Detailed information about cloned antibodies with paired heavy and light chains. Variable (V), diversity (D) and joining (J) genes, mutation on the variable gene (V MUT), and CDR3 amino acid sequences of cloned immunoglobulin heavy, kappa light and lambda light chains are listed. These antibodies are grouped by their IGHV genes, with our selected H001-H020 antibodies indicated.

|  | **DONOR ID** | **HEAVY CHAIN** | | | | | **KAPPA LIGHT CHAIN** | | | | **LAMBDA LIGHT CHAIN** | | | |
| --- | --- | --- | --- | --- | --- | --- | --- | --- | --- | --- | --- | --- | --- | --- |
|  |  | **V** | **D** | **J** | **V MUT** | **CDR3** | **V** | **J** | **V MUT** | **CDR3** | **V** | **J** | **V MUT** | **CDR3** |
|  | 9 | IGHV1-18*01 | IGHD6-13*01 | IGHJ6*02 | 5 | ARNGYSSSWHGGTHYYYYALDF | IGKV1-5*03 | IGKJ1*01 | 5 | QQYYSYPWT |  |  |  |  |
|  | 55 | IGHV1-18*01 | IGHD2-15*01 | IGHJ4*02 | 12 | ARDSVSWWNLLFKSLEKLTLDY |  |  |  |  | IGLV2-23*01 | IGLJ3*02 | 55 | CSYSGSTTCV |
|  | 55 | IGHV1-18*01 | IGHD2-15*01 | IGHJ4*02 | 12 | ARDSVSWWNLLFKSLEKLTLDY | IGKV3-20*01 | IGKJ4*01 | 4 | QQFGSSPLT | IGLV2-8*01 | IGLJ1*01 | 7 | NSYAGNNNFV |
|  | 55 | IGHV1-18*01 | IGHD2-15*01 | IGHJ4*02 | 12 | ARDSVSWWNLLFKSLEKLTLDY | IGKV3-11*01 | IGKJ1*01 | 1 | QHRSNSWT |  |  |  |  |
|  | 55 | IGHV1-18*01 | IGHD4-17*01 | IGHJ4*02 | 10 | ARDPDFGDYGSDIVDY |  |  |  |  | IGLV3-25*02 | IGLJ3*02 | 7 | QSADSTATYWV |
|  | 99 | IGHV1-18*01 | IGHD3-16*01 | IGHJ6*03 | 15 | ARWGVGMTFSYHYHYMDV |  |  |  |  | IGLV3-19*01 | IGLJ3*02 | 16 | GSRDNSGYS |
|  | 99 | IGHV1-18*01 | IGHD6-13*01 | IGHJ4*02 | 9 | ARGRGDSSTWYSLY | IGKV3-11*01 | IGKJ4*01 | 9 | QQYNDWLT |  |  |  |  |
|  | 146 | IGHV1-18*01 | IGHD1-1*01 | IGHJ1*01 | 21 | AGDTTSSAALTF | IGKV4-1*01 | IGKJ3*01 | 14 | QQYYTTPS |  |  |  |  |
|  | 146 | IGHV1-18*01 | IGHD1-26*01 | IGHJ6*02 | 16 | ARFFGGATMTVYFYGLDV | IGKV1-5*01 | IGKJ1*01 | 7 | QQYNSYSGWT |  |  |  |  |
|  | 146 | IGHV1-18*01 | IGHD6-19*01 | IGHJ4*02 | 7 | TRSEQWRSRGEY |  |  |  |  | IGLV1-44*01 | IGLJ3*02 | 13 | ASWDDSLSGSWV |
|  | 146 | IGHV1-18*01 | IGHD1-26*01 | IGHJ5*02 | 8 | GRDDSGSYPMSP | IGKV3-20*01 | IGKJ4*01 | 15 | QQYXSSPLA |  |  |  |  |
|  | 146 | IGHV1-18*01 | IGHD1-20*01 | IGHJ4*02 | 19 | VRDINFIFDY |  |  |  |  | IGLV7-43*01 | IGLJ3*02 | 10 | LLYXWSSSALG |
|  | 69 | IGHV1-18*04 | IGHD3-10*01 | IGHJ6*03 | 10 | AREGWFGEFRRNYNYNYYMDV | IGKV3-15*01 | IGKJ1*01 | 7 | QQYHEWPRT |  |  |  |  |
|  | 49 | IGHV1-24*01 | IGHD4-11*01 | IGHJ3*01 | 28 | TLVTRVDAFDV | IGKV1-39*01 | IGKJ2*01 | 6 | QQSYFAPYT |  |  |  |  |
|  | 49 | IGHV1-24*01 | IGHD4-11*01 | IGHJ3*01 | 13 | TLVTRVDAFEV | IGKV1-39*01 | IGKJ2*01 | 7 | QQSYFAPYT |  |  |  |  |
|  | 49 | IGHV1-24*01 | IGHD2-21*02 | IGHJ3*01 | 17 | TLVTGVDAFAV | IGKV1-39*01 | IGKJ2*01 | 11 | QQSYFAPYT |  |  |  |  |
|  | 49 | IGHV1-24*01 | IGHD3-16*01 | IGHJ3*01 | 16 | TSVIKADAFEV | IGKV1-39*01 | IGKJ2*01 | 12 | QQSYFAPYT |  |  |  |  |
|  | 13 | IGHV1-3*01 | IGHD2-21*01 | IGHJ6*02 | 21 | ARGGWIIQNGGARYYHGMDV | IGKV3-20*01 | IGKJ4*01 | 10 | QHYNNPVA | IGLV3-21*02 | IGLJ2*01 | 13 | QVWDGGSYHVI |
|  | 13 | IGHV1-3*01 | IGHD3-10*01 | IGHJ4*02 | 20 | ARKDYYGSGSYEFDN |  |  |  |  | IGLV2-11*01 | IGLJ2*01 | 6 | CSYAGNYILV |
|  | 69 | IGHV1-3*01 | IGHD2-2*01 | IGHJ3*01 | 27 | AKGRSSHDLYDPFDF | IGKV1-5*03 | IGKJ2*01 | 15 | QQYNSYSRYT |  |  |  |  |
|  | 99 | IGHV1-3*01 | IGHD3-10*01 | IGHJ4*02 | 10 | ARKNYYASGSYHFDL |  |  |  |  | IGLV2-11*01 | IGLJ3*02 | 6 | SSYAGKYTLV |
|  | 146 | IGHV1-3*01 | IGHD1-1*01 | IGHJ3*02 | 14 | ARVGIPLRGAGGSPFDI | IGKV1-39*01 | IGKJ4*01 | 15 | QQSSSVPLT |  |  |  |  |
| **H001** | 146 | IGHV1-3*01 | IGHD3-16*01 | IGHJ3*02 | 8 | ARVGILVRGAGGSPFDI | IGKV1-39*01 | IGKJ4*01 | 7 | QQSYSTPLT |  |  |  |  |
|  | 146 | IGHV1-3*01 | IGHD3-16*01 | IGHJ3*02 | 8 | ARVGILVRGAGGSPFDI | IGKV1-39*01 | IGKJ4*01 | 5 | QQSYSTPLT |  |  |  |  |
|  | 146 | IGHV1-3*01 | IGHD4-11*01 | IGHJ3*01 | 10 | AREDYTGNYYDAFDF | IGKV1-5*03 | IGKJ4*01 | 7 | QLYNSYSGTT |  |  |  |  |
|  | 146 | IGHV1-3*01 | IGHD6-13*01 | IGHJ6*02 | 13 | ARDGVKEQLVYYYFGMDV |  |  |  |  | IGLV1-40*01 | IGLJ2*01 | 11 | QSYDSNLSGSVV |
|  | 146 | IGHV1-3*01 | IGHD3-10*01 | IGHJ4*02 | 12 | ARGALLWFRDDFDF | IGKV1-27*01 | IGKJ3*01 | 4 | QKYDSAPYT | IGLV1-51*01 | IGLJ1*01 | 5 | GSWDSSLYSFYV |
|  | 99 | IGHV1-46*01 | IGHD2-2*01 | IGHJ1*01 | 10 | ASRLDAIPFQV | IGKV3-20*01 | IGKJ2*01 | 11 | QQYGGLPFT |  |  |  |  |
|  | 99 | IGHV1-46*01 | IGHD4-23*01 | IGHJ6*04 | 29 | ARKGNYGSRYDWYFDV |  |  |  |  | IGLV2-14*01 | IGLJ3*02 | 6 | SSYTSSSTGV |
|  | 55 | IGHV1-46*03 | IGHD3-9*01 | IGHJ3*01 | 21 | TRDPILRYFDWQSRDAFDV | IGKV1-17*01 | IGKJ3*01 | 6 | LQHNGYPIT |  |  |  |  |
|  | 60 | IGHV1-69*01 | IGHD3-10*01 | IGHJ6*01 | 11 | ARVPSVATCNFGCYSAMDV | IGKV1-39*01 | IGKJ3*01 | 5 | QQSYSTLFS | IGLV2-14*01 | IGLJ1*01 | 4 | SSYTGSSTRYV |
|  | 146 | IGHV1-69*06 | IGHD3-3*01 | IGHJ4*02 | 13 | ARASFGDLWSGYPNQFFDH | IGKV1-17*03 | IGKJ1*01 | 6 | LQHNTYPWT |  |  |  |  |
|  | 13 | IGHV2-5*02 | IGHD3-9*01 | IGHJ5*02 | 10 | AHRLLTAYYDH | IGKV1-5*03 | IGKJ1*01 | 5 | QQYNSYSWA |  |  |  |  |
|  | 55 | IGHV2-5*02 | IGHD6-13*01 | IGHJ5*02 | 1 | AHTVAAAATFWFDP |  |  |  |  | IGLV1-44*01 | IGLJ2*01 | 2 | ATWDDSLNGLV |
|  | 69 | IGHV2-5*02 | IGHD2-21*02 | IGHJ3*02 | 6 | AHSSYFDCGGDCSDVAFDI |  |  |  |  | IGLV3-10*01 | IGLJ3*02 | 11 | YSTDSSGDPV |
|  | 146 | IGHV2-5*02 | IGHD2-15*01 | IGHJ3*01 | 9 | ARSYCRGGNCYSTAFNV |  |  |  |  | IGLV2-11*01 | IGLJ2*01 | 7 | CSYAGAYTYVA |
| **H002** | 146 | IGHV2-70D*04 | IGHD7-27*01 | IGHJ4*02 | 4 | ARSNHWGSHFDY |  |  |  |  | IGLV3-21*02 | IGLJ1*01 | 6 | QVWDTSGDHLYV |
|  | 55 | IGHV3-11*01 | IGHD2-21*01 | IGHJ3*01 | 9 | ARDLPGDEYLDAFDL | IGKV4-1*01 | IGKJ4*01 | 7 | QQFYTAPLT |  |  |  |  |
|  | 60 | IGHV3-11*01 | IGHD6-13*01 | IGHJ6*02 | 11 | ASGAAVPYFYYGVDV |  |  |  |  | IGLV3-1*01 | IGLJ2*01 | 11 | QAWGSSPAKV |
|  | 99 | IGHV3-13*01 | IGHD3-22*01 | IGHJ4*02 | 9 | ARAVHYYDSSGHYSGYYFDY |  |  |  |  | IGLV1-44*01 | IGLJ2*01 | 6 | ATWDASLKGVV |
|  | 55 | IGHV3-15*01 | IGHD1-1*01 | IGHJ4*02 | 6 | TTQNAFES |  |  |  |  | IGLV3-21*01 | IGLJ1*01 | 0 | QVWDSSSDHYV |
|  | 55 | IGHV3-15*01 | IGHD2/OR15-2a*01 | IGHJ6*02 | 7 | HTLSTTHYYGMDV | IGKV3-20*01 | IGKJ4*01 | 5 | QQYINSPLT |  |  |  |  |
|  | 146 | IGHV3-15*01 | IGHD6-25*01 | IGHJ4*02 | 12 | AAHNRAAY | IGKV1-5*01 | IGKJ2*01 | 8 | QQYYSYPLT |  |  |  |  |
|  | 9 | IGHV3-21*01 | IGHD3-3*02 | IGHJ4*02 | 9 | ARARPPGTAFGFDH |  |  |  |  | IGLV1-40*01 | IGLJ2*01 | 7 | QSYDSSLSGAL |
|  | 13 | IGHV3-21*01 | IGHD3-16*01 | IGHJ4*02 | 10 | VRTFYFDY | IGKV3-15*01 | IGKJ1*01 | 7 | QQYDIWPPRT |  |  |  |  |
|  | 55 | IGHV3-21*01 | IGHD4-17*01 | IGHJ1*01 | 7 | VRDMTTVTTCXXQH |  |  |  |  | IGLV1-44*01 | IGLJ2*01 | 3 | AAWDDSLNGLV |
|  | 55 | IGHV3-21*01 | IGHD4-17*01 | IGHJ1*01 | 7 | VRDMTTVTTCYLQH |  |  |  |  | IGLV1-44*01 | IGLJ2*01 | 4 | AAWDDSLNGLV |
|  | 146 | IGHV3-21*01 | IGHD4-17*01 | IGHJ4*02 | 12 | ARGRTYGDSN | IGKV3-15*01 | IGKJ1*01 | 7 | QQYDNWHT |  |  |  |  |
|  | 146 | IGHV3-21*01 | IGHD3-3*01 | IGHJ4*02 | 10 | ARVPILLAQGVPTFDL | IGKV3-15*01 | IGKJ2*01 | 7 | QQYSDWPRYT |  |  |  |  |
|  | 9 | IGHV3-23*01 | IGHD2-2*02 | IGHJ6*03 | 14 | AKAAILGNYNYYMDV |  |  |  |  | IGLV3-21*02 | IGLJ2*01 | 8 | QVWDSSADHLVV |
|  | 13 | IGHV3-23*01 | IGHD1-1*01 | IGHJ4*01 | 14 | VKDAIYMSNWPWYFDY |  |  |  |  | IGLV3-21*02 | IGLJ2*01 | 7 | XXXX |
|  | 13 | IGHV3-23*01 | IGHD6-13*01 | IGHJ4*02 | 2 | AKDPIYSSSWPYYFDY |  |  |  |  | IGLV3-21*02 | IGLJ2*01 | 6 | QVWDSSSDHPEVV |
|  | 13 | IGHV3-23*01 | IGHD6-13*01 | IGHJ4*02 | 2 | AKDPIYTSRWPYYFDY | IGKV1-39*01 | IGKJ2*03 | 0 | QQSYSLYS | IGLV3-21*02 | IGLJ2*01 | 1 | QVWHSSSDHSEVI |
|  | 13 | IGHV3-23*01 | IGHD3-10*01 | IGHJ4*02 | 9 | AKDGVLGSYHQYYFQY |  |  |  |  | IGLV3-21*02 | IGLJ2*01 | 3 | QVWDNSSDHPGVV |
|  | 13 | IGHV3-23*01 | IGHD5-12*01 | IGHJ4*02 | 1 | AKGSRNGPYIVATLHFDY | IGKV3-15*01 | IGKJ4*01 | 3 | QHYNHWSLT |  |  |  |  |
|  | 55 | IGHV3-23*01 | IGHD6-13*01 | IGHJ4*02 | 6 | VLSSSWMDNPFDF |  |  |  |  | IGLV5-45*02 | IGLJ3*02 | 0 | MIWHSSAWV |
|  | 55 | IGHV3-23*01 | IGHD1-26*01 | IGHJ4*02 | 9 | AGFPSGTHFFDY | IGKV3-11*01 | IGKJ1*01 | 3 | QQRSNWWT |  |  |  |  |
|  | 55 | IGHV3-23*01 | IGHD1-26*01 | IGHJ4*02 | 9 | AGFPSGTHFFDY | IGKV3-20*01 | IGKJ4*01 | 4 | QQFGSSPLT | IGLV2-8*01 | IGLJ1*01 | 7 | NSYAGNNNFV |
|  | 55 | IGHV3-23*01 | IGHD2-2*01 | IGHJ6*02 | 4 | ARPDALHCSSITSCSLYGLAYYYGMDV |  |  |  |  | IGLV1-44*01 | IGLJ3*02 | 4 | AAWDDRLIGWV |
|  | 55 | IGHV3-23*01 | IGHD1-26*01 | IGHJ4*02 | 9 | AKRMVEATNRYFDY | IGKV3-20*01 | IGKJ2*01 | 2 | QQYGSSPPYT |  |  |  |  |
|  | 55 | IGHV3-23*01 | IGHD1-26*01 | IGHJ4*02 | 13 | ARDSSEWVLGIDF |  |  |  |  | IGLV2-11*01 | IGLJ2*01 | 4 | CSYAGSYTVV |
|  | 60 | IGHV3-23*01 | IGHD4/OR15-4a*01 | IGHJ2*01 | 2 | AKDAVRSANHAWYFDF |  |  |  |  | IGLV3-21*02 | IGLJ2*01 | 3 | QVWDSNSDHPKVV |
|  | 60 | IGHV3-23*01 | IGHD5-12*01 | IGHJ4*02 | 22 | AKGYGLFDS | IGKV3-15*01 | IGKJ1*01 | 15 | QQYINWPPWS |  |  |  |  |
| **H003** | 60 | IGHV3-23*01 | IGHD2-21*01 | IGHJ2*01 | 11 | AKDAIRNSNHAWYFDV |  |  |  |  | IGLV3-21*02 | IGLJ2*01 | 8 | QVWDPTSDQV |
|  | 60 | IGHV3-23*01 | IGHD2-8*01 | IGHJ4*02 | 13 | AKSRVTNSGSIDH | IGKV1-39*01 | IGKJ5*01 | 6 | QQSYSTSIT |  |  |  |  |
|  | 60 | IGHV3-23*01 | IGHD4/OR15-4a*01 | IGHJ2*01 | 12 | AKDAILSANHPWYFDF | IGKV4-1*01 | IGKJ4*01 | 0 | QQYYSTPLT | IGLV3-21*02 | IGLJ3*02 | 7 | QVWDSSSDHPKVV |
|  | 60 | IGHV3-23*01 | IGHD2-21*01 | IGHJ2*01 | 5 | AKDAIRSSNHPWYFHV |  |  |  |  | IGLV3-21*02 | IGLJ2*01 | 4 | QVWDGSSDHPKVL |
|  | 60 | IGHV3-23*01 | IGHD4/OR15-4a*01 | IGHJ2*01 | 14 | AKDAILSANHPWYFDF |  |  |  |  | IGLV3-21*02 | IGLJ3*02 | 12 | QVWDRSSDQSKVV |
|  | 146 | IGHV3-23*01 | IGHD3/OR15-3a*01 | IGHJ4*02 | 18 | AKGYGLFDF | IGKV3-15*01 | IGKJ1*01 | 12 | QQYINRPPWT |  |  |  |  |
|  | 9 | IGHV3-23*03 | IGHD1-1*01 | IGHJ4*02 | 8 | AKVLLGGWNGVFDH | IGKV3-11*01 | IGKJ4*01 | 6 | QQRSTWPPS |  |  |  |  |
| **H004** | 146 | IGHV3-23*04 | IGHD3-10*01 | IGHJ6*02 | 21 | AKDGYFGSGSLYGIDV |  |  |  |  | IGLV3-21*02 | IGLJ2*01 | 10 | QVWDSNHDHPGVV |
|  | 146 | IGHV3-23*04 | IGHD3-10*01 | IGHJ6*02 | 3 | AKDGYYGSGSLYGMDV |  |  |  |  | IGLV3-21*02 | IGLJ2*01 | 4 | QVWDSTSDHPGVV |
|  | 146 | IGHV3-23*04 | IGHD3-9*01 | IGHJ6*02 | 8 | AKVIQYPRGFWFYGMDV |  |  |  |  | IGLV1-51*01 | IGLJ3*02 | 1 | GTWDSSLNNCV |
|  | 60 | IGHV3-30-3*01 | IGHD3-10*01 | IGHJ6*02 | 9 | ARDPGVPYYHYAMDV |  |  |  |  | IGLV2-8*01 | IGLJ2*01 | 2 | SSYAGSNNYIL |
|  | 146 | IGHV3-30-3*01 | IGHD3-3*01 | IGHJ4*02 | 17 | VRDETDWEIGVVVATPEFDY | IGKV1-17*01 | IGKJ2*01 | 0 | LQHNSYPRT | IGLV1-44*01 | IGLJ2*01 | 10 | STWDDSLNGVV |
|  | 146 | IGHV3-30-3*02 | IGHD4-23*01 | IGHJ4*02 | 16 | VTGIRARDYGGSTFDL |  |  |  |  | IGLV10-54*01 | IGLJ2*01 | 11 | AAWDSSLSAMI |
|  | 9 | IGHV3-30*02 | IGHD3-10*01 | IGHJ3*01 | 13 | AKDGRWFGESGGFDV |  |  |  |  | IGLV3-21*02 | IGLJ2*01 | 9 | QVWESSTDPVV |
|  | 9 | IGHV3-30*03 | IGHD2-2*01 | IGHJ2*01 | 11 | ARDYCSRTNCINWIFDL | IGKV3-20*01 | IGKJ4*01 | 5 | QQYGSSPLT |  |  |  |  |
|  | 60 | IGHV3-30*03 | IGHD2-8*02 | IGHJ6*02 | 20 | AKDGYLSAARGYGMDV |  |  |  |  | IGLV3-21*02 | IGLJ2*01 | 10 | QVWETTSDQLV |
|  | 146 | IGHV3-30*03 | IGHD4-17*01 | IGHJ4*02 | 9 | ARDTFGDYYFDY |  |  |  |  | IGLV1-40*01 | IGLJ1*01 | 3 | QSYDSRLSVPYV |
|  | 9 | IGHV3-30*04 | IGHD5-18*01 | IGHJ6*02 | 11 | ARGGGYTYGSYYYSMDV | IGKV4-1*01 | IGKJ3*01 | 2 | QQYYSTPFT |  |  |  |  |
|  | 9 | IGHV3-30*04 | IGHD5-18*01 | IGHJ6*02 | 6 | ARGGGYTYGSYYYAMDV | IGKV4-1*01 | IGKJ3*01 | 2 | QQYYSTPFT |  |  |  |  |
|  | 55 | IGHV3-30*04 | IGHD1-26*01 | IGHJ4*02 | 12 | ARALLSVVGSKSYYFDF | IGKV3-11*01 | IGKJ1*01 | 1 | QHRSNSWT |  |  |  |  |
|  | 55 | IGHV3-30*04 | IGHD1-26*01 | IGHJ4*02 | 12 | ARALLSVVGSKSYYFDF | IGKV3-11*01 | IGKJ1*01 | 1 | QHRSNSWT |  |  |  |  |
|  | 55 | IGHV3-30*04 | IGHD1-26*01 | IGHJ6*02 | 11 | ARDEKYSGLYSGRTGDYYYGMDV |  |  |  |  | IGLV1-51*01 | IGLJ2*01 | 6 | GTWDSSLSLGV |
|  | 55 | IGHV3-30*04 | IGHD3-22*01 | IGHJ6*02 | 5 | ARDGKLGRTYHDSRQSYFYIMDV |  |  |  |  | IGLV2-14*01 | IGLJ2*01 | 6 | SSYTSSTSLV |
|  | 13 | IGHV3-30*18 | IGHD3-10*01 | IGHJ6*02 | 12 | AKDAYYFASGSFFGMDV |  |  |  |  | IGLV3-21*02 | IGLJ2*01 | 7 | QVWGSGGVI |
|  | 13 | IGHV3-30*18 | IGHD3-10*01 | IGHJ6*02 | 1 | AKDAYYYGSGYGMDV | IGKV1-5*03 | IGKJ4*01 | 6 | QQYTSFST | IGLV1-47*01 | IGLJ1*01 | 2 | AAWDDRLSGYV |
|  | 60 | IGHV3-30*18 | IGHD3-10*01 | IGHJ6*02 | 9 | AKDGYVVSGSGYGMDV |  |  |  |  | IGLV3-21*02 | IGLJ2*01 | 6 | QVWDSSSDHVV |
| **H009** | 60 | IGHV3-30*18 | IGHD1-26*01 | IGHJ6*02 | 15 | AKTDIKWGATNYGMDV |  |  |  |  | IGLV3-21*02 | IGLJ2*01 | 10 | QVWDGTRDHLVV |
|  | 60 | IGHV3-30*18 | IGHD5-12*01 | IGHJ4*02 | 9 | AKDSAGYGLHY | IGKV1-17*01 | IGKJ1*01 | 6 | LQHNSYPWT | IGLV3-19*01 | IGLJ2*01 | 5 | NSRDSIGNHVV |
|  | 60 | IGHV3-30*18 | IGHD3-22*01 | IGHJ6*02 | 21 | AKDGYLSAARGYGMDV |  |  |  |  | IGLV3-21*02 | IGLJ2*01 | 9 | QVWDTTTDQLV |
|  | 60 | IGHV3-30*18 | IGHD3-22*01 | IGHJ6*02 | 21 | AKDGYLSAARGYGMDV |  |  |  |  | IGLV3-21*02 | IGLJ2*01 | 10 | QVWDTTSDQLV |
|  | 60 | IGHV3-30*18 | IGHD3-22*01 | IGHJ6*02 | 21 | AKDGYLSAARGYGMDV |  |  |  |  | IGLV3-21*02 | IGLJ2*01 | 11 | QVWDTSSDQLV |
| **H005** | 60 | IGHV3-30*18 | IGHD3-16*01 | IGHJ6*02 | 20 | AKDAYLSAARGYGMHV |  |  |  |  | IGLV3-21*02 | IGLJ2*01 | 11 | QVWDTASDQLV |
|  | 60 | IGHV3-30*18 | IGHD3-22*01 | IGHJ6*02 | 18 | AKDGYLSAARGYGMDV |  |  |  |  | IGLV3-21*02 | IGLJ2*01 | 9 | QVWDTTSDQLV |
|  | 60 | IGHV3-30*18 | IGHD3-22*01 | IGHJ6*02 | 21 | AKDGYLSAARGYGMDV |  |  |  |  | IGLV3-21*02 | IGLJ2*01 | 13 | QVWDTTSDQLV |
|  | 60 | IGHV3-30*18 | IGHD3-22*01 | IGHJ6*02 | 18 | AKDGYLSAARGYGMDV |  |  |  |  | IGLV3-21*02 | IGLJ2*01 | 8 | QVWDTTSDQLV |
|  | 60 | IGHV3-30*18 | IGHD1-26*01 | IGHJ6*02 | 6 | AKTDIRWGATNYGMDV |  |  |  |  | IGLV3-21*02 | IGLJ2*01 | 5 | QVWDGSSDHLVV |
| **H010** | 60 | IGHV3-30*18 | IGHD3-3*01 | IGHJ4*02 | 12 | AKGPLFGLFSFDQ |  |  |  |  | IGLV3-21*03 | IGLJ2*01 | 9 | QVWDNSRNRGI |
|  | 60 | IGHV3-30*18 | IGHD3-22*01 | IGHJ6*02 | 18 | AKDGYLSAARGYGMDV |  |  |  |  | IGLV3-21*02 | IGLJ2*01 | 9 | QVWDTTSDQLV |
|  | 60 | IGHV3-30*18 | IGHD3-22*01 | IGHJ6*02 | 24 | AKDGYLSAARGYGMDV | IGKV2-30*01 | IGKJ1*01 | 8 | TQVTLWPPWT |  |  |  |  |
| **H006** | 60 | IGHV3-30*18 | IGHD3-22*01 | IGHJ6*02 | 21 | AKDGYLSAARGYGMDV |  |  |  |  | IGLV3-21*02 | IGLJ2*01 | 11 | QVWDTTSDQLV |
|  | 60 | IGHV3-30*18 | IGHD3-22*01 | IGHJ6*02 | 19 | AKDGYLSAARGYGMDV |  |  |  |  | IGLV2-18*02 | IGLJ2*01 | 0 | SSYTSSSTLV |
|  | 60 | IGHV3-30*18 | IGHD2-21*02 | IGHJ4*02 | 9 | AKDPIKVSANGWGFDY |  |  |  |  | IGLV3-21*03 | IGLJ2*01 | 7 | QVWDSNSDHVV |
|  | 60 | IGHV3-30*18 | IGHD3-22*01 | IGHJ6*02 | 19 | AKDGYLSAARGFGMDV |  |  |  |  | IGLV3-21*02 | IGLJ2*01 | 13 | QVWDTTSDQLV |
|  | 60 | IGHV3-30*18 | IGHD2-15*01 | IGHJ6*02 | 13 | AKTDIMWRAVNYGMDV |  |  |  |  | IGLV3-21*02 | IGLJ2*01 | 12 | QVWDDSRDHLVI |
|  | 60 | IGHV3-30*18 | IGHD2-15*01 | IGHJ6*02 | 13 | AKTDIMWRAVNYGMDV |  |  |  |  | IGLV3-21*02 | IGLJ2*01 | 12 | QVWDDSRDHLVI |
|  | 60 | IGHV3-30*18 | IGHD2-21*01 | IGHJ6*02 | 8 | AKTDIMWQAVNYGMDV |  |  |  |  | IGLV3-21*02 | IGLJ2*01 | 9 | QVWDDSRDHLVI |
|  | 60 | IGHV3-30*18 | IGHD3-22*01 | IGHJ6*02 | 18 | AKDGYLSAARGYGMDV |  |  |  |  | IGLV3-21*02 | IGLJ2*01 | 9 | QVWDTTSDQLV |
|  | 60 | IGHV3-30*18 | IGHD2-15*01 | IGHJ6*02 | 12 | AKTDIMWRAVNYGMDV |  |  |  |  | IGLV3-21*02 | IGLJ2*01 | 12 | QVWDESRDHLVI |
|  | 60 | IGHV3-30*18 | IGHD3-22*01 | IGHJ6*02 | 18 | AKDGYLSAARGYGMDV |  |  |  |  | IGLV3-21*02 | IGLJ2*01 | 9 | QVWDTTSDQLV |
| **H008** | 146 | IGHV3-30*18 | IGHD1-26*01 | IGHJ6*02 | 19 | AKDGILGARRGLYGIDV |  |  |  |  | IGLV3-21*02 | IGLJ2*01 | 10 | QVWDSSSDHVV |
| **H007** | 146 | IGHV3-30*18 | IGHD3-3*01 | IGHJ6*02 | 9 | AKEIGGFDFRSGSQRSYYYYGVDV | IGKV3-20*01 | IGKJ2*01 | 10 | QQYSSSPPGYT |  |  |  |  |
|  | 146 | IGHV3-30*18 | IGHD3-3*01 | IGHJ6*02 | 7 | AKEIGGFDFRSGDQLTYYYYGMDV | IGKV3-20*01 | IGKJ2*01 | 9 | HQYVTSPPGYT |  |  |  |  |
|  | 146 | IGHV3-30*18 | IGHD5-18*01 | IGHJ6*02 | 9 | AKDAYIYARGSYYGMDV |  |  |  |  | IGLV3-21*02 | IGLJ2*01 | 6 | QVWDSSSNDPVV |
|  | 146 | IGHV3-30*18 | IGHD1-26*01 | IGHJ6*02 | 15 | AKDGILGARRGXYGIDV |  |  |  |  | IGLV3-21*02 | IGLJ2*01 | 5 | QVWDSYSDHVV |
|  | 146 | IGHV3-30*18 | IGHD3-3*01 | IGHJ6*02 | 10 | AKEIGGFDFRSGKQRSYYYYGVDV | IGKV3-20*01 | IGKJ2*01 | 9 | QQYGNSPPGYT |  |  |  |  |
|  | 146 | IGHV3-30*18 | IGHD3-10*01 | IGHJ4*02 | 12 | AKDAYYYGSGSHNNPDY |  |  |  |  | IGLV3-21*02 | IGLJ3*02 | 9 | QVWDSSSMGV |
|  | 146 | IGHV3-30*18 | IGHD3-16*02 | IGHJ4*02 | 7 | AKGYDYIWGTYRPRPDLDS |  |  |  |  | IGLV1-44*01 | IGLJ3*02 | 13 | ASWDDSLSGSWV |
|  | 146 | IGHV3-30*18 | IGHD6-13*01 | IGHJ4*02 | 10 | AKDPVQRSNWYYFDY |  |  |  |  | IGLV3-21*02 | IGLJ2*01 | 10 | QVWHSTTEPVV |
|  | 146 | IGHV3-30*18 | IGHD1-20*01 | IGHJ4*02 | 10 | AKDPVHRSNWFYFDH |  |  |  |  | IGLV3-21*02 | IGLJ2*01 | 12 | QVWYSNSEPVV |
|  | 146 | IGHV3-30*18 | IGHD3-3*01 | IGHJ6*02 | 18 | AKGSPIIRFLMMDV | IGKV1-5*03 | IGKJ1*01 | 11 | QYYSVYST | IGLV3-21*02 | IGLJ3*02 | 6 | QVWDNNSDHVV |
|  | 13 | IGHV3-33*01 | IGHD6-13*01 | IGHJ4*02 | 6 | AREAGIAAPASLDF |  |  |  |  | IGLV3-21*02 | IGLJ2*01 | 9 | QVWDSSSDHVV |
|  | 13 | IGHV3-33*01 | IGHD6-13*01 | IGHJ4*02 | 8 | TREAGIAAPAALDY |  |  |  |  | IGLV3-21*02 | IGLJ2*01 | 6 | QVWDSSSYHVV |
| **H011** | 13 | IGHV3-33*01 | IGHD1-7*01 | IGHJ3*01 | 15 | VRDNWSYNAFDV | IGKV3-11*01 | IGKJ2*03 | 13 | QHRNSWPYS |  |  |  |  |
|  | 13 | IGHV3-33*01 | IGHD6-19*01 | IGHJ4*02 | 10 | VRENSGWYYFDY | IGKV1-17*01 | IGKJ4*01 | 5 | LQHDSYPFT |  |  |  |  |
| **H012** | 13 | IGHV3-33*01 | IGHD6-13*01 | IGHJ4*02 | 12 | AREGAIAAPASLDV |  |  |  |  | IGLV3-21*02 | IGLJ2*01 | 10 | QVWDSGTDHVI |
|  | 13 | IGHV3-33*01 | IGHD6-13*01 | IGHJ4*02 | 9 | AREGGIAAPAALDF |  |  |  |  | IGLV3-21*02 | IGLJ2*01 | 12 | QVWDGGSYHVI |
|  | 13 | IGHV3-33*01 |  | IGHJ4*02 | 8 | ARESKAYPYYFDY | IGKV1-39*01 | IGKJ2*03 | 23 | QHSSFPPQDS |  |  |  |  |
|  | 13 | IGHV3-33*01 | IGHD1-1*01 | IGHJ4*02 | 11 | ARESNGFGSDF | IGKV1-17*01 | IGKJ1*01 | 5 | LQHNSFPRT |  |  |  |  |
|  | 13 | IGHV3-33*01 | IGHD3-10*01 | IGHJ3*02 | 8 | VRDNWSYNAFDI | IGKV3-11*01 | IGKJ2*03 | 10 | QHRNSWPYS |  |  |  |  |
|  | 13 | IGHV3-33*01 | IGHD1-20*01 | IGHJ3*02 | 5 | ARDNWKYNAFDI | IGKV3-11*01 | IGKJ2*03 | 3 | QHRSNWPYS |  |  |  |  |
|  | 13 | IGHV3-33*01 | IGHD6-19*01 | IGHJ4*02 | 7 | ARESSGWYYFDY |  |  |  |  | IGLV2-23*01 | IGLJ1*01 | 7 | CLYAGSSISYV |
|  | 13 | IGHV3-33*01 | IGHD6-13*01 | IGHJ5*02 | 7 | AREERIAAPASLDL |  |  |  |  | IGLV3-21*02 | IGLJ2*01 | 10 | QVWDSSTYHVV |
|  | 13 | IGHV3-33*01 | IGHD6-25*01 | IGHJ4*02 | 11 | ARELRIAAPAALDY |  |  |  |  | IGLV3-21*02 | IGLJ2*01 | 13 | QVWDSGSDHVL |
|  | 49 | IGHV3-33*01 | IGHD3-16*01 | IGHJ4*02 | 6 | ARQMFTGHFDY | IGKV3-15*01 | IGKJ1*01 | 7 | QHYNNWPRT |  |  |  |  |
|  | 60 | IGHV3-33*01 | IGHD2-2*01 | IGHJ6*02 | 10 | AREGRGQLLFHGMDV | IGKV3-20*01 | IGKJ3*01 | 3 | QQYGRSQGFT |  |  |  |  |
|  | 60 | IGHV3-33*01 | IGHD2-21*02 | IGHJ4*02 | 18 | ARDGHCDGGCYSALYDY |  |  |  |  | IGLV2-23*01 | IGLJ3*02 | 6 | CSFAGSRWV |
| **H015** | 60 | IGHV3-33*01 | IGHD6-6*01 | IGHJ4*02 | 13 | AREDPHLLIATLDL | IGKV1-39*01 | IGKJ4*01 | 15 | QQSYGTPALA |  |  |  |  |
|  | 60 | IGHV3-33*01 | IGHD4-11*01 | IGHJ2*01 | 9 | ARETTIFNWYFDL | IGKV1-39*01 | IGKJ4*01 | 15 | QQSFSIPPT |  |  |  |  |
|  | 60 | IGHV3-33*01 | IGHD5-18*01 | IGHJ4*02 | 6 | ARERRGFSYGLDDN |  |  |  |  | IGLV2-11*01 | IGLJ2*01 | 4 | CSYAGRYTFVV |
|  | 60 | IGHV3-33*01 | IGHD6-19*01 | IGHJ3*01 | 12 | AREQAEIAVASFDF | IGKV1-39*01 | IGKJ4*01 | 18 | QQAYNAPPLT |  |  |  |  |
|  | 60 | IGHV3-33*01 | IGHD6-19*01 | IGHJ4*02 | 10 | ARESHYSAWYVLDY |  |  |  |  | IGLV1-51*01 | IGLJ3*02 | 13 | GSWDGSLSVGV |
|  | 60 | IGHV3-33*01 | IGHD2-8*01 | IGHJ4*02 | 4 | AREDPNVFIATLDL | IGKV1-39*01 | IGKJ4*01 | 10 | QQSDSTPALA |  |  |  |  |
|  | 60 | IGHV3-33*01 | IGHD3-16*01 | IGHJ4*02 | 10 | AREDPYVFMATLDS |  |  |  |  | IGLV2-11*01 | IGLJ3*02 | 0 | CSYAGSYTWV |
|  | 60 | IGHV3-33*01 | IGHD2-21*01 | IGHJ2*01 | 15 | ARETTIFQWYFDL | IGKV1-39*01 | IGKJ4*01 | 10 | QQSFSIPPT |  |  |  |  |
|  | 69 | IGHV3-33*01 | IGHD2-15*01 | IGHJ6*02 | 3 | ARETTFGRFCSGGSCYSDYYYGMDV | IGKV2-24*01 | IGKJ1*01 | 4 | MQAAQFPWT |  |  |  |  |
|  | 69 | IGHV3-33*01 | IGHD6-13*01 | IGHJ4*02 | 13 | AREGGIVAADK |  |  |  |  | IGLV1-51*01 | IGLJ1*01 | 6 | XXXX |
| **H014** | 146 | IGHV3-33*01 | IGHD4/OR15-4a*01 | IGHJ4*02 | 15 | AREARVAAPASYDY |  |  |  |  | IGLV3-21*02 | IGLJ3*02 | 14 | QVWDNGSNHVV |
|  | 146 | IGHV3-33*01 | IGHD6-13*01 | IGHJ4*02 | 7 | AREALIAAPATFDY |  |  |  |  | IGLV3-21*02 | IGLJ2*01 | 10 | QVWDNNSRHVV |
|  | 146 | IGHV3-33*01 | IGHD6-13*01 | IGHJ5*02 | 14 | AREGHIAAPAALDL |  |  |  |  | IGLV3-21*02 | IGLJ2*01 | 7 | QVWDSSSEHVV |
| **H013** | 146 | IGHV3-33*01 | IGHD6-13*01 | IGHJ4*02 | 13 | AREANIAAPAIYDH |  |  |  |  | IGLV3-21*02 | IGLJ2*01 | 5 | QVWDSYSDHVV |
|  | 146 | IGHV3-33*01 | IGHD2-15*01 | IGHJ5*02 | 8 | AREGHVATPILDL | IGKV1-39*01 | IGKJ4*01 | 8 | QQSYSMPTLT |  |  |  |  |
|  | 146 | IGHV3-33*01 | IGHD3-10*01 | IGHJ3*01 | 12 | VRDNFGLNAFDV | IGKV3-11*01 | IGKJ2*01 | 9 | QHRSNWPYT |  |  |  |  |
|  | 146 | IGHV3-33*01 | IGHD6-13*01 | IGHJ4*02 | 7 | ATERRIAAPGCLDY |  |  |  |  | IGLV5-48*02 | IGLJ2*01 | 22 | MIXHSSAMW |
|  | 146 | IGHV3-33*01 | IGHD6-13*01 | IGHJ4*02 | 5 | AREALIAAPATFDY | IGKV1-5*03 | IGKJ1*01 | 11 | QYYSVYST | IGLV3-21*02 | IGLJ3*02 | 6 | QVWDNNSDHVV |
|  | 60 | IGHV3-33*02 | IGHD5-12*01 | IGHJ6*02 | 12 | AGGGYSSRGYYNYGLDV | IGKV1-33*01 | IGKJ4*01 | 2 | QQYDNLPPLT | IGLV3-10*01 | IGLJ2*01 | 18 | YSTDRSGDQRV |
|  | 99 | IGHV3-33*06 | IGHD2-15*01 | IGHJ4*02 | 3 | AKATCGDGSCGLYYFDY |  |  |  |  | IGLV1-40*01 | IGLJ3*02 | 4 | QSYDSNLSGWV |
|  | 55 | IGHV3-43*02 | IGHD2-21*01 | IGHJ5*02 | 8 | AKDIWIFDGRRWIAGSPDA | IGKV2-30*01 | IGKJ1*01 | 5 | MQQTHWPWA |  |  |  |  |
|  | 49 | IGHV3-48*01 | IGHD1-1*01 | IGHJ6*02 | 16 | ARAGPPSPPNYGMDV | IGKV4-1*01 | IGKJ4*01 | 9 | QQYYTAPLL |  |  |  |  |
| **H016** | 69 | IGHV3-48*02 | IGHD3/OR15-3a*01 | IGHJ2*01 | 16 | ASVGLDSKISGYWYFDL | IGKV3-15*01 | IGKJ4*01 | 11 | QQYDHWPLT |  |  |  |  |
|  | 69 | IGHV3-48*02 | IGHD3/OR15-3a*01 | IGHJ2*01 | 7 | ARVGLALTISGYWYFDL | IGKV3-15*01 | IGKJ4*01 | 4 | QQYNDWPLT |  |  |  |  |
|  | 146 | IGHV3-48*02 | IGHD7-27*01 | IGHJ2*01 | 13 | ARAKLGSGSYWYFDL | IGKV3-15*01 | IGKJ4*01 | 12 | QQYNNWPLT |  |  |  |  |
|  | 13 | IGHV3-48*03 | IGHD1-1*01 | IGHJ3*02 | 15 | ARDTGIWNGAYDAFDI |  |  |  |  | IGLV5-45*02 | IGLJ1*01 | 5 | MIWHSTAYV |
|  | 13 | IGHV3-48*03 |  | IGHJ4*02 | 12 | VGFDH | IGKV4-1*01 | IGKJ5*01 | 7 | QQYYSPPIT |  |  |  |  |
|  | 55 | IGHV3-48*03 | IGHD2-2*02 | IGHJ4*02 | 9 | AGHCSSNKCYKY |  |  |  |  | IGLV1-40*01 | IGLJ2*01 | 3 | QSYDSSLSGSI |
|  | 69 | IGHV3-49*03 | IGHD3-22*01 | IGHJ4*02 | 15 | ARDLTINKIIVANDF | IGKV4-1*01 | IGKJ5*01 | 4 | QQYYSTPIT |  |  |  |  |
|  | 146 | IGHV3-49*03 | IGHD6-13*01 | IGHJ4*02 | 6 | ARVPYSSSWYVAWADY | IGKV4-1*01 | IGKJ4*01 | 3 | QQYYSTPLT |  |  |  |  |
|  | 99 | IGHV3-49*04 | IGHD3-10*01 | IGHJ4*02 | 9 | TRGDYYGSRNSYFWLFDY | IGKV2D-29*01 | IGKJ1*01 | 5 | MQSIQLRT |  |  |  |  |
|  | 69 | IGHV3-53*01 | IGHD6-19*01 | IGHJ4*02 | 5 | ARVIAVAGTNRGGPRWRSTYYFDY |  |  |  |  | IGLV7-43*01 | IGLJ3*02 | 14 | LLYCSGVRV |
|  | 9 | IGHV3-7*01 | IGHD3-3*01 | IGHJ4*02 | 23 | ASELWTAFNKDWSGYNDY |  |  |  |  | IGLV1-51*01 | IGLJ3*02 | 12 | GTWDSSLKVVV |
|  | 9 | IGHV3-7*01 | IGHD1-26*01 | IGHJ4*02 | 8 | TRDTWVDS | IGKV1-17*03 | IGKJ3*01 | 8 | LQHQSYPFT |  |  |  |  |
|  | 13 | IGHV3-7*01 | IGHD2-21*01 | IGHJ4*02 | 8 | ASSHYSAGDVSYNFDY |  |  |  |  | IGLV10-54*01 | IGLJ3*02 | 9 | SAWDFSLRAWV |
|  | 55 | IGHV3-7*03 | IGHD6-13*01 | IGHJ5*02 | 4 | ARSHVAAGVTRWFDP | IGKV3-11*01 | IGKJ2*01 | 2 | QHRSK |  |  |  |  |
|  | 55 | IGHV3-7*03 | IGHD6-13*01 | IGHJ5*01 | 10 | ARSHVAAGGTRWIDS | IGKV3-11*01 | IGKJ2*01 | 10 | XXXX |  |  |  |  |
|  | 55 | IGHV3-7*03 | IGHD3-16*02 | IGHJ3*01 | 10 | ARLDRGTGESGYRSSDV |  |  |  |  | IGLV1-44*01 | IGLJ2*01 | 6 | AAWDDSLNGVV |
|  | 55 | IGHV3-7*03 | IGHD1-7*01 | IGHJ5*02 | 7 | ARSHVAASGTRWFDP | IGKV3-11*01 | IGKJ2*01 | 3 | XXXX |  |  |  |  |
|  | 55 | IGHV3-72*01 | IGHD3-9*01 | IGHJ4*02 | 3 | ARDGYDILNHFVRFDF | IGKV3-11*01 | IGKJ2*01 | 3 | XXXX |  |  |  |  |
|  | 60 | IGHV3-72*01 | IGHD3-16*01 | IGHJ3*02 | 7 | AREGLGSPTSDAFDI |  |  |  |  | IGLV2-14*01 | IGLJ3*02 | 9 | SSYTTSSTLV |
|  | 60 | IGHV3-73*02 | IGHD4-17*01 | IGHJ6*02 | 5 | TSQYGDGYYYAMDV | IGKV3-15*01 | IGKJ3*01 | 7 | QHYNNWPLFT |  |  |  |  |
|  | 9 | IGHV3-74*01 |  | IGHJ6*02 | 13 | VRGLNHAMDV |  |  |  |  | IGLV2-14*01 | IGLJ3*02 | 13 | CSFTTRNTWV |
|  | 13 | IGHV3-74*01 | IGHD3-22*01 | IGHJ4*02 | 11 | ARVYRDSRDGSDFRHFDS | IGKV4-1*01 | IGKJ4*01 | 8 | QQYYDIPYT |  |  |  |  |
| **H017** | 55 | IGHV3-74*01 | IGHD3-10*01 | IGHJ4*02 | 5 | ARGSTYYFGSGSVDY | IGKV4-1*01 | IGKJ3*01 | 0 | QQYYSTPLT | IGLV2-14*01 | IGLJ1*01 | 9 | SSYRGSSTPYV |
|  | 55 | IGHV3-74*01 | IGHD1-26*01 | IGHJ4*02 | 7 | ARGSLDF |  |  |  |  | IGLV2-14*01 | IGLJ2*01 | 8 | YSYTTSNTLV |
|  | 55 | IGHV3-74*01 | IGHD1-26*01 | IGHJ4*02 | 10 | SRGSLDY |  |  |  |  | IGLV2-14*01 | IGLJ2*01 | 9 | YSYTTSNTLV |
|  | 99 | IGHV3-74*01 | IGHD4/OR15-4a*01 | IGHJ4*02 | 6 | ACLRVPDRN |  |  |  |  | IGLV7-46*01 | IGLJ2*01 | 1 | LLSYSGAQV |
|  | 13 | IGHV3-9*01 | IGHD2-2*01 | IGHJ4*02 | 14 | VKANVKKGSTSCFDY | IGKV3-20*01 | IGKJ1*01 | 5 | QQYSGSSPRT |  |  |  |  |
|  | 60 | IGHV3-9*01 | IGHD6-19*01 | IGHJ4*01 | 13 | VKDKSQGIPVAGLEY |  |  |  |  | IGLV1-51*01 | IGLJ2*01 | 5 | GTWDSSLSAA |
|  | 60 | IGHV3-9*01 | IGHD6-19*01 | IGHJ4*02 | 12 | VKDKSQGIPLAGLEY |  |  |  |  | IGLV1-51*01 | IGLJ2*01 | 7 | ATWDSSLTAA |
| **H018** | 60 | IGHV3-9*01 | IGHD6-19*01 | IGHJ4*01 | 14 | VKDKSQGIPVAGLEY |  |  |  |  | IGLV1-51*01 | IGLJ2*01 | 7 | GTWDSSLSAA |
|  | 69 | IGHV4-30-4*01 | IGHD2-2*03 | IGHJ3*02 | 11 | AIYMDEAWAFEI | IGKV1-39*01 | IGKJ2*01 | 7 | QQSYTISLFT |  |  |  |  |
| **H019** | 69 | IGHV4-30-4*01 | IGHD2-2*03 | IGHJ3*02 | 11 | AIYMDEAWAFEI | IGKV1-39*01 | IGKJ2*01 | 9 | QQSYTISLFT |  |  |  |  |
|  | 146 | IGHV4-30-4*01 | IGHD3-10*01 | IGHJ4*02 | 7 | VRENYITSPLSR |  |  |  |  | IGLV2-8*01 | IGLJ3*02 | 4 | SSYAGSNDVV |
|  | 146 | IGHV4-30-4*01 | IGHD4-17*01 | IGHJ4*02 | 10 | ASYTVTTWGGFDY |  |  |  |  | IGLV3-10*01 | IGLJ3*02 | 11 | YSTDSSGNYRV |
|  | 99 | IGHV4-31*02 | IGHD4/OR15-4a*01 | IGHJ4*02 | 4 | ARGVLH | IGKV3-20*01 | IGKJ2*01 | 3 | QQYGSSPYT |  |  |  |  |
|  | 13 | IGHV4-31*03 | IGHD3-22*01 | IGHJ4*02 | 39 | ARVVSSGHRHYYFDY | IGKV1-27*01 | IGKJ1*01 | 7 | QKYWT |  |  |  |  |
|  | 60 | IGHV4-31*03 | IGHD1-1*01 | IGHJ5*02 | 36 | AQSRRLVGPFVS | IGKV3-15*01 | IGKJ2*03 | 11 | QQYDKWPRS | IGLV6-57*01 | IGLJ3*02 | 0 | QSYDSSNHGV |
|  | 99 | IGHV4-31*03 | IGHD2-8*02 | IGHJ4*02 | 5 | ARGVLV | IGKV3-20*01 | IGKJ2*01 | 3 | QQYDSSPYT |  |  |  |  |
|  | 146 | IGHV4-31*03 | IGHD3-10*01 | IGHJ3*01 | 15 | ARVVHASANAFDV | IGKV1-6*01 | IGKJ4*01 | 1 | LQDYNYPLT | IGLV1-51*01 | IGLJ3*02 | 5 | GTWDSSLNGWV |
|  | 146 | IGHV4-31*03 | IGHD3-10*01 | IGHJ3*01 | 14 | ARVVHASANAFDV | IGKV1-39*01 | IGKJ5*01 | 13 | QQTYSTPT |  |  |  |  |
|  | 146 | IGHV4-31*03 | IGHD4-11*01 | IGHJ3*02 | 23 | ARVPLRDFYSNYSPSAFDI | IGKV3-15*01 | IGKJ4*01 | 7 | QQYKNWPPLT |  |  |  |  |
|  | 9 | IGHV4-34*01 | IGHD3-16*02 | IGHJ5*02 | 10 | AGGRFTNDFVWGSYRYES | IGKV2-28*01 | IGKJ4*01 | 2 | MQALQTLLLT |  |  |  |  |
|  | 60 | IGHV4-34*01 | IGHD6-19*01 | IGHJ4*02 | 14 | VRGGYSSAPYPREWRY |  |  |  |  | IGLV1-44*01 | IGLJ3*02 | 0 | AAWDDSLNGWV |
|  | 69 | IGHV4-34*01 | IGHD5-24*01 | IGHJ6*03 | 10 | ARGRDGYNYVGYYYYYYMDV | IGKV3-11*01 | IGKJ1*01 | 5 | QQRSNWQWT |  |  |  |  |
|  | 146 | IGHV4-34*01 | IGHD3-3*01 | IGHJ6*02 | 11 | ARGIFEVVIIPYYSYRVDV | IGKV3-15*01 | IGKJ3*01 | 3 | QQYNNWPPFT | IGLV3-19*01 | IGLJ2*01 | 13 | SSRSGNRLV |
|  | 9 | IGHV4-34*02 | IGHD6-13*01 | IGHJ5*02 | 3 | GRGLGREYSSSWYGGRRFDP |  |  |  |  | IGLV2-14*01 | IGLJ3*02 | 40 | RSYISNNXXWV |
|  | 9 | IGHV4-38-2*02 | IGHD3-22*01 | IGHJ3*02 | 7 | ARDRSGYVFFYDAFDI | IGKV3-20*01 | IGKJ4*01 | 2 | QQYGSSPLT |  |  |  |  |
|  | 146 | IGHV4-38-2*02 | IGHD2-21*02 | IGHJ4*02 | 15 | GGGVTRADY |  |  |  |  | IGLV1-47*02 | IGLJ3*02 | 7 | AVWDDNLSAWE |
|  | 60 | IGHV4-39*01 | IGHD2-8*01 | IGHJ4*02 | 8 | ARRIQLMVFDF | IGKV1-5*03 | IGKJ1*01 | 6 | HQYNTYPWT |  |  |  |  |
|  | 60 | IGHV4-39*01 | IGHD3-3*01 | IGHJ2*01 | 13 | ARHPYYNFWIYWYFDL |  |  |  |  | IGLV1-44*01 | IGLJ2*01 | 3 | ASWDDSLNGLVV |
|  | 99 | IGHV4-39*01 | IGHD3-10*01 | IGHJ3*01 | 19 | TRHWLGGDKWSQSPFLAV |  |  |  |  | IGLV1-44*01 | IGLJ3*02 | 9 | AVWDDSLNTWV |
|  | 99 | IGHV4-39*01 | IGHD5-12*01 | IGHJ3*02 | 6 | AKGRYSGYNDYNAFDI |  |  |  |  | IGLV1-47*02 | IGLJ3*02 | 3 | AAWDDSPEWLG |
|  | 146 | IGHV4-39*01 | IGHD3-10*01 | IGHJ3*01 | 9 | TRPASGAHDYVSRSYYPGQGAFGV | IGKV3-20*01 | IGKJ5*01 | 1 | QQYGSSSIT |  |  |  |  |
|  | 146 | IGHV4-39*01 | IGHD6-13*01 | IGHJ4*02 | 15 | ARLLGIAATGHFDS |  |  |  |  | IGLV10-54*01 | IGLJ3*02 | 7 | STWDSSLSTWL |
|  | 146 | IGHV4-39*01 | IGHD3-10*01 | IGHJ3*02 | 10 | TRPASGAHDYASRSYYPGLGAFGI | IGKV3-20*01 | IGKJ5*01 | 4 | QQYGSSSTT |  |  |  |  |
|  | 146 | IGHV4-39*01 | IGHD3-3*01 | IGHJ3*02 | 15 | ARPLLNPMTLYGVTPGIGPFEI | IGKV3-20*01 | IGKJ4*01 | 10 | QQHDNSLS |  |  |  |  |
|  | 146 | IGHV4-39*01 | IGHD6-6*01 | IGHJ4*02 | 18 | AAHRVSSSYPADY | IGKV1-39*01 | IGKJ1*01 | 14 | QQSYNTPT |  |  |  |  |
|  | 146 | IGHV4-39*01 | IGHD3-3*01 | IGHJ3*02 | 12 | ARPLLNPSTIYGVTPGIGPFEM | IGKV3D-15*01 | IGKJ1*01 | 14 | QQYINWPPWT |  |  |  |  |
|  | 146 | IGHV4-4*02 | IGHD3-16*02 | IGHJ6*02 | 8 | ARGNYDYVWGSYRSDQGYGLDV | IGKV1-39*01 | IGKJ2*01 | 12 | QRSYSTPYT |  |  |  |  |
|  | 49 | IGHV4-4*07 | IGHD6-13*01 | IGHJ6*03 | 17 | AREGGSSYYYYYYMDV |  |  |  |  | IGLV2-8*01 | IGLJ2*01 | 6 | SSYAGINSYVI |
|  | 13 | IGHV4-59*01 | IGHD2-2*01 | IGHJ5*02 | 5 | ARSKNQLLLFDP | IGKV3-15*01 | IGKJ4*01 | 5 | QQYNDWPPLT |  |  |  |  |
|  | 13 | IGHV4-59*01 | IGHD4-23*01 | IGHJ5*01 | 15 | ARSKNQLLLFEF | IGKV1-33*01 | IGKJ4*01 | 3 | QQYDNLPLT |  |  |  |  |
|  | 55 | IGHV4-59*01 | IGHD7-27*01 | IGHJ3*01 | 18 | ARTNWAYDPFNV | IGKV1-5*03 | IGKJ1*01 | 21 | QQYYSYST | IGLV1-36*01 | IGLJ2*01 | 68 | SAWDFSLSVQV |
|  | 60 | IGHV4-59*01 | IGHD2-15*01 | IGHJ6*02 | 1 | ARDRGYCSGGSCLGGMDV | IGKV1-5*03 | IGKJ4*01 | 2 | QQYNSYFPLT |  |  |  |  |
|  | 60 | IGHV4-59*01 | IGHD2-8*02 | IGHJ6*02 | 8 | VRDRGFCTGKSCLGGMDV | IGKV1-5*03 | IGKJ1*01 | 0 | QQYNSYRT |  |  |  |  |
|  | 60 | IGHV4-59*01 | IGHD2-8*02 | IGHJ4*02 | 9 | ARLRRRGLTGTDFDY |  |  |  |  | IGLV1-44*01 | IGLJ2*01 | 8 | AAWDDSLNGPNVV |
|  | 69 | IGHV4-59*01 | IGHD3-22*01 | IGHJ6*03 | 2 | ARSYYYDSSGYRPSFYYYYMDV |  |  |  |  | IGLV3-1*01 | IGLJ2*01 | 2 | QAWDSSVV |
|  | 146 | IGHV4-59*01 | IGHD1-26*01 | IGHJ6*02 | 6 | ARGILGSTWYYYYGLDV | IGKV1-39*01 | IGKJ2*01 | 6 | HQSYSSPYT |  |  |  |  |
|  | 55 | IGHV4-59*08 | IGHD5-18*01 | IGHJ4*02 | 11 | ARHLYRYGYRNYFDY | IGKV1-12*01 | IGKJ4*01 | 5 | QQANSFPLT |  |  |  |  |
| **H020** | 55 | IGHV4-59*08 | IGHD5-18*01 | IGHJ4*02 | 11 | ARHLYRYGYRNYFDY | IGKV1-12*01 | IGKJ4*01 | 5 | QQANSFPLT |  |  |  |  |
|  | 69 | IGHV4-61*08 | IGHD6-13*01 | IGHJ3*02 | 22 | ARMTSFKQSGGWYRGRHDGFDI | IGKV1-5*01 | IGKJ2*01 | 14 | QEYNSYSYT |  |  |  |  |
|  | 69 | IGHV4-61*08 | IGHD6-19*01 | IGHJ3*02 | 9 | ARLTSYKQRGGWYRGRHDAFDI | IGKV1-5*01 | IGKJ2*01 | 7 | QEYSSYSYT |  |  |  |  |
|  | 9 | IGHV5-51*01 | IGHD3-22*01 | IGHJ4*02 | 7 | ARRARNVGNYGTSDFYPYFDH |  |  |  |  | IGLV2-14*01 | IGLJ3*02 | 7 | SSYISSNTLWV |
|  | 60 | IGHV5-51*01 | IGHD1-14*01 | IGHJ3*02 | 16 | ARRRVSVTGTDAFDI |  |  |  |  | IGLV2-8*01 | IGLJ2*01 | 4 | SSYAGTNTFVV |
|  | 60 | IGHV5-51*01 | IGHD1-14*01 | IGHJ3*02 | 16 | ARRRVSVTGTDAFDI |  |  |  |  | IGLV2-8*01 | IGLJ2*01 | 4 | SSYAGTNTFVV |
|  | 69 | IGHV5-51*01 | IGHD1-1*01 | IGHJ4*02 | 15 | ARATPGNYYFDS |  |  |  |  | IGLV2-14*01 | IGLJ1*01 | 9 | SSYTDSSPNCV |
|  | 99 | IGHV5-51*01 | IGHD2-2*01 | IGHJ1*01 | 12 | ARPSRSRDINKWYLSTSEYFHY | IGKV3-20*01 | IGKJ4*01 | 7 | QQYANSPLT |  |  |  |  |
|  | 9 | IGHV6-1*01 | IGHD1-1*01 | IGHJ4*02 | 8 | AVGHHWHFKY |  |  |  |  | IGLV2-8*01 | IGLJ1*01 | 5 | SSHAGSNYGV |
|  | 9 | IGHV6-1*01 | IGHD1-1*01 | IGHJ4*02 | 6 | AVGHHWHFKY |  |  |  |  | IGLV2-8*01 | IGLJ1*01 | 5 | SSHAGSNYGV |
|  | 55 | IGHV6-1*01 | IGHD3-10*01 | IGHJ4*02 | 9 | ARDTYYYTSASYYNVDY | IGKV1-39*01 | IGKJ5*01 | 7 | QQSYRTPIT |  |  |  |  |
|  | 55 | IGHV7-4-1*02 | IGHD1-7*01 | IGHJ4*02 | 14 | ARLGEYSWNSIGYFDY |  |  |  |  | IGLV1-40*01 | IGLJ2*01 | 11 | QSYDRSLILVV |
|  | 99 | IGHV7-4-1*02 | IGHD3-10*01 | IGHJ4*02 | 12 | ARSYAYGDF | IGKV1-39*01 | IGKJ1*01 | 19 | QQSDTLPWT |  |  |  |  |
|  | 99 | IGHV7-4-1*02 | IGHD3-22*01 | IGHJ4*02 | 2 | ARGARSYYDSSGYYSWSDY | IGKV4-1*01 | IGKJ1*01 | 2 | QQYYNTLTWA |  |  |  |  |
