## Supplemental Table 3 for "Public broadly neutralizing antibodies against hepatitis B virus in individuals with elite serologic activity"

Table S3. Primers and synthesized nucleotide sequences.

| Single-Cell Antibody Cloning Primers | |
| --- | --- |
| F1-HC | 5'-ACAGGTGCCCACTCCCAGGTGCAG |
| F2-HC | 5'-AAGGTGTCCAGTGTGARGTGCAG |
| F3-HC | 5'-CCCAGATGGGTCCTGTCCCAGGTGCAG |
| F4-HC | 5'-CAAGGAGTCTGTTCCGAGGTGCAG |
| R-1st-HC | 5'-GGAAGGTGTGCACGCCGCTGGTC |
| R-2nd-HC | 5'-GTTCGGGGAAGTAGTCCTTGAC |
| F1-Kappa | 5'-ATGAGGSTCCCYGCTCAGCTGCTGG |
| F2-Kappa | 5'-CTCTTCCTCCTGCTACTCTGGCTCCCAG |
| F3-Kappa | 5'-ATTTCTCTGTTGCTCTGGATCTCTG |
| F4-Kappa | 5'-ATGACCCAGWCTCCABYCWCCCTG |
| R-1st-Kappa | 5'-GTTTCTCGTAGTCTGCTTTGCTCA |
| R-2nd-Kappa | 5'-GTGCTGTCCTTGCTGTCCTGCT |
| F1-Lambda | 5'-GGTCCTGGGCCCAGTCTGTGCTG |
| F2-Lambda | 5'-GGTCCTGGGCCCAGTCTGCCCTG |
| F3-Lambda | 5'-GCTCTGTGACCTCCTATGAGCTG |
| F4-Lambda | 5'-GGTCTCTCTCSCAGCYTGTGCTG |
| F5-Lambda | 5'-GTTCTTGGGCCAATTTTATGCTG |
| F6-Lambda | 5'-GGTCCAATTCYCAGGCTGTGGTG |
| F7-Lambda | 5'-GAGTGGATTCTCAGACTGTGGTG |
| R-1st-Lambda | 5'-CACCAGTGTGGCCTTGTTGGCTTG |
| R-2nd-Lambda | 5'-CTCCTCACTCGAGGGYGGGAACAGAGTG |
| F-Vector Seq Primer | 5'-GCTTCGTTAGAACGCGGCTAC |
| Antibody Cloning Primers for Each Antibody | |
| F-H001-HC | 5'-CTAGTAGCAACTGCAACCGGTGTACATTCT CAGGTCCAGCTTGTGCAGTCTGG |
| R-H001-HC | 5'-CCGATGGGCCCTTGGTCGACGC TGAAGAGACGGTGACCATTGTCCCTT |
| F-H001-Kappa | 5'-CTAGTAGCAACTGCAACCGGTGTACATTCT GACATCCAGATGACCCAGTCTCCA |
| R-H001-Kappa | 5'-GAAGACAGATGGTGCAGCCACCGTACG TTTGATCTCCACCTTGGTCCCTCC |
| F-H002-HC | 5'-CTAGTAGCAACTGCAACCGGTGTACATTCT CAGGTCACCTTGAAGGAGTCTGG |
| R-H002-HC | 5'-CCGATGGGCCCTTGGTCGACGC TGAGGAGACGGTGACCAGGGTG |
| F-H002-Lambda | 5'-CTAGTAGCAACTGCAACCGGTTCCTGGGCC TCCTATGTGCTGACCCAGGCGC |
| F-H003-HC | 5'-CTAGTAGCAACTGCAACCGGTGTACATTCT GAGATGCAGCTGCTGGAGTCTGG |
| R-H003-HC | 5'-CCGATGGGCCCTTGGTCGACGC TGAGGAGACAGTGACCAGGGTGC |
| F-H003-Lambda | 5'-CTAGTAGCAACTGCAACCGGTTCCTGGGCC TCCTATATGCTGACTCAGGCACCC |
| F-H004-HC | 5'-CTAGTAGCAACTGCAACCGGTGTACATTCT GAGATGCAACTGGTGGAGTCTGG |
| R-H004-HC | 5'-CCGATGGGCCCTTGGTCGACGC TGAGGAGACGATGACCGTGGTCC |
| F-H004-Lambda | 5'-CTAGTAGCAACTGCAACCGGTTCCTGGGCC TCCTATGTGCTGACTCAGCCACC |
| F-H005-HC | 5'-CTAGTAGCAACTGCAACCGGTGTACATTCT CAGATGCGTCTGGTGGAATCTGG |
| R-H005-HC | 5'-CCGATGGGCCCTTGGTCGACGC TGAGGAGACGGTGACCGGGGT |
| F-H005-Lambda | 5'-CTAGTAGCAACTGCAACCGGTTCCTGGGCC TCCTATGTGCTGACTCAGCCACC |
| F-H006-HC | 5'-CTAGTAGCAACTGCAACCGGTGTACATTCT CAGATGCGTCTGGTGGAATCGGG |
| R-H006-HC | 5'-CCGATGGGCCCTTGGTCGACGC TGAGGAGACGGTGACCGGGATC |
| F-H006-Lambda | 5'-CTAGTAGCAACTGCAACCGGTTCCTGGGCC TCCTATGTGCTGACTCAGCCACC |
| F-H007-HC | 5'-CTAGTAGCAACTGCAACCGGTGTACATTCT CAGGTGCACCTGGTGGAGTCTG |
| R-H007-HC | 5'-CCGATGGGCCCTTGGTCGACGC TGAGGAGACGGTGACCGTGGTC |
| F-H007-Kappa | 5'-CTAGTAGCAACTGCAACCGGTGTACATTCT GAAATTGTGTTGACGCAGTCTCCAG |
| R-H007-Kappa | 5'-GAAGACAGATGGTGCAGCCACCGTACG TTTGATCTCCAACTTGGTCCCCTGG |
| F-H008-HC | 5'-CTAGTAGCAACTGCAACCGGTGTACATTCT CAGATGCACCTATTGGAGTCTGGG |
| R-H008-HC | 5'-CCGATGGGCCCTTGGTCGACGC TGACGAGACGGTGACCCTGGTC |
| F-H008-Lambda | 5'-CTAGTAGCAACTGCAACCGGTTCCTGGGCC TCCTATGTGCTGACTCAGCCACC |
| F-H009-HC | 5'-CTAGTAGCAACTGCAACCGGTGTACATTCT CAGATGAAGTTGGTGGAGTCTGGG |
| R-H009-HC | 5'-CCGATGGGCCCTTGGTCGACGC TGAGGAGACGGTGACCGTGGTC |
| F-H009-Lambda | 5'-CTAGTAGCAACTGCAACCGGTTCCTGGGCC TCCTATGTGCTGACTCAGCCACC |
| F-H010-HC | 5'-CTAGTAGCAACTGCAACCGGTGTACATTCT CAGGTGCAGCTGGTGGAGTCTG |
| R-H010-HC | 5'-CCGATGGGCCCTTGGTCGACGC TGAGGAGACGGTGACCAGGGTT |
| F-H010-Lambda | 5'-CTAGTAGCAACTGCAACCGGTTCCTGGGCC TCCTATGTGCTGACTCAGCCACC |
| F-H011-HC | 5'-CTAGTAGCAACTGCAACCGGTGTACATTCT CAGGTTCACTTGGCGGAGTCTGG |
| R-H011-HC | 5'-CCGATGGGCCCTTGGTCGACGC TGAAGAGACGGTGACCAATGTCCC |
| F-H011-Kappa | 5'-CTAGTAGCAACTGCAACCGGTGTACATTCT GAAGTTGTGTTGACACAGTCTCCAGC |
| R-H011-Kappa | 5'-GAAGACAGATGGTGCAGCCACCGTACG TTTGATCTCCAGCTTGGTCCCCTG |
| F-H012-HC | 5'-CTAGTAGCAACTGCAACCGGTGTACATTCT CAGCTGCAGCTGGTGGAGTCTG |
| R-H012-HC | 5'-CCGATGGGCCCTTGGTCGACGC TGAGGAGACGGTGACCAGGGTTC |
| F-H012-Lambda | 5'-CTAGTAGCAACTGCAACCGGTTCCTGGGCC TCCTATGTGCTGACTCAGCCACC |
| F-H013-HC | 5'-CTAGTAGCAACTGCAACCGGTGTACATTCT CAGGTGCAGCTGGTGGAGTCTG |
| R-H013-HC | 5'-CCGATGGGCCCTTGGTCGACGC TGAGGAGACGGTGACCAGGGCT |
| F-H013-Lambda | 5'-CTAGTAGCAACTGCAACCGGTTCCTGGGCC TCCTATGTGCTGACTCAGCCACC |
| F-H014-HC | 5'-CTAGTAGCAACTGCAACCGGTGTACATTCT CAGGTACAACTGATGGAGTCTGGG |
| R-H014-HC | 5'-CCGATGGGCCCTTGGTCGACGC TGAGGAGACGGTGACCAGGGCT |
| F-H014-Lambda | 5'-CTAGTAGCAACTGCAACCGGTTCCTGGGCC TCCTATGTGCTGACTCAGACACCC |
| F-H015-HC | 5'-CTAGTAGCAACTGCAACCGGTGTACATTCT CAGGTGCAGCTGGTGGAGTCTG |
| R-H015-HC | 5'-CCGATGGGCCCTTGGTCGACGC TGAGGAGACGGTGACCAGGGTTC |
| F-H015-Kappa | 5'-CTAGTAGCAACTGCAACCGGTGTACATTCT GACATCCAGGTGACCCAGTCAC |
| R-H015-Kappa | 5'-GAAGACAGATGGTGCAGCCACCGTACG TTTGATGTCCACCTTGGTCCCTCC |
| F-H016-HC | 5'-CTAGTAGCAACTGCAACCGGTGTACATTCT GAGATGCACCTGGTGGAGTCTGG |
| R-H016-HC | 5'-CCGATGGGCCCTTGGTCGACGC TGAGGAGACAGTGACCAGGGTGC |
| F-H016-Kappa | 5'-CTAGTAGCAACTGCAACCGGTGTACATTCT GAAATACTGCTGACGCAGTCTCCAG |
| R-H016-Kappa | 5'-GAAGACAGATGGTGCAGCCACCGTACG TTTGATCTCCACCTTGGTCCCTCC |
| F-H017-HC | 5'-CTAGTAGCAACTGCAACCGGTGTACATTCT GAGGTGCAGCTGGTGGAGTCC |
| R-H017-HC | 5'-CCGATGGGCCCTTGGTCGACGC TGAGGAGACGGTGACCAGGGTTC |
| F-H017-Lambda | 5'-CTAGTAGCAACTGCAACCGGTTCCTGGGCC CAGTCTGCCCTGACTCAGCCTG |
| F-H018-HC | 5'-CTAGTAGCAACTGCAACCGGTGTACATTCT GAAGGACAGCTGGTGGAGTCTGG |
| R-H018-HC | 5'-CCGATGGGCCCTTGGTCGACGC TGAGGAGACGGTGACCAGGGTTC |
| F-H018-Lambda | 5'-CTAGTAGCAACTGCAACCGGTTCCTGGGCC CAGTCTGTGTTGACGCAGCCGC |
| F-H019-HC | 5'-CTAGTAGCAACTGCAACCGGTGTACATTCT CAGGTGGTGCTGCAGGAGTCG |
| R-H019-HC | 5'-CCGATGGGCCCTTGGTCGACGC TGAAGAGACGGCGACCAGTGTCC |
| F-H019-Kappa | 5'-CTAGTAGCAACTGCAACCGGTGTACATTCT GACATCCAGATGACCCAGTCTCCG |
| R-H019-Kappa | 5'-GAAGACAGATGGTGCAGCCACCGTACG TTTGATCTCCAGCTTGGTCCCCTG |
| F-H020-HC | 5'-CTAGTAGCAACTGCAACCGGTGTACATTCT CAGGTGCAGCTGCAGGAGTCG |
| R-H020-HC | 5'-CCGATGGGCCCTTGGTCGACGC TGAGGAGACGGTGACCAGGTTTCC |
| F-H020-Kappa | 5'-CTAGTAGCAACTGCAACCGGTGTACATTCT GACATCCAGATGACCCAGTCTCCA |
| R-H020-Kappa | 5'-GAAGACAGATGGTGCAGCCACCGTACG TTTGAACTCCACCTTGGTCCCTCC |
| R-Lambda | 5'-GGCTTGAAGCTCCTCACTCGAGGGYGGGAACAGAGTG |
| gBlock Synthesis for Alanine Mutations | |
| 101Q | 5'-ACAAGAATCCTCACAATACCGCAGAGTCTAGACTCGTGGTGGACTTCTCTCAATTTTCTAGGGGGATCTCCCGTGTGTCTTGGCCAAAATTCGCAGTCCCCAACCTCCAATCACTCACCAACCTCCTGTCCTCCAATTTGTCCTGGTTATCGCTGGATGTGTCTGCGGCGTTTTATCATATTCCTCTTCATCCTGCTGCTATGCCTCATCTTCTTATTGGTTCTTCTGGATTATGCAGGTATGTTGCCCGTTTGTCCTCTAATTCCAGGATCAACAACAACCAGTACGGGACCATGCAAAACCTGCACGACTCCTGCTCAAGGCAACTCTATGTTTCCCTCATGTTGCTGTACAAAACCTACGGATGGAAATTGCACCTGTATTCCCATCCCATCGTCCTGGGCTTTCGCAAAATACCTATGGGAGTGGGCCTCAGTCCGTTTCTCTTGGCTCAGTTTACTAGTGCCATTTGTTCAGTGGTTCGTAGGG |
| 102G | 5'-ACAAGAATCCTCACAATACCGCAGAGTCTAGACTCGTGGTGGACTTCTCTCAATTTTCTAGGGGGATCTCCCGTGTGTCTTGGCCAAAATTCGCAGTCCCCAACCTCCAATCACTCACCAACCTCCTGTCCTCCAATTTGTCCTGGTTATCGCTGGATGTGTCTGCGGCGTTTTATCATATTCCTCTTCATCCTGCTGCTATGCCTCATCTTCTTATTGGTTCTTCTGGATTATCAAGCAATGTTGCCCGTTTGTCCTCTAATTCCAGGATCAACAACAACCAGTACGGGACCATGCAAAACCTGCACGACTCCTGCTCAAGGCAACTCTATGTTTCCCTCATGTTGCTGTACAAAACCTACGGATGGAAATTGCACCTGTATTCCCATCCCATCGTCCTGGGCTTTCGCAAAATACCTATGGGAGTGGGCCTCAGTCCGTTTCTCTTGGCTCAGTTTACTAGTGCCATTTGTTCAGTGGTTCGTAGGG |
| 103M | 5'-ACAAGAATCCTCACAATACCGCAGAGTCTAGACTCGTGGTGGACTTCTCTCAATTTTCTAGGGGGATCTCCCGTGTGTCTTGGCCAAAATTCGCAGTCCCCAACCTCCAATCACTCACCAACCTCCTGTCCTCCAATTTGTCCTGGTTATCGCTGGATGTGTCTGCGGCGTTTTATCATATTCCTCTTCATCCTGCTGCTATGCCTCATCTTCTTATTGGTTCTTCTGGATTATCAAGGTGCATTGCCCGTTTGTCCTCTAATTCCAGGATCAACAACAACCAGTACGGGACCATGCAAAACCTGCACGACTCCTGCTCAAGGCAACTCTATGTTTCCCTCATGTTGCTGTACAAAACCTACGGATGGAAATTGCACCTGTATTCCCATCCCATCGTCCTGGGCTTTCGCAAAATACCTATGGGAGTGGGCCTCAGTCCGTTTCTCTTGGCTCAGTTTACTAGTGCCATTTGTTCAGTGGTTCGTAGGG |
| 104L | 5'-ACAAGAATCCTCACAATACCGCAGAGTCTAGACTCGTGGTGGACTTCTCTCAATTTTCTAGGGGGATCTCCCGTGTGTCTTGGCCAAAATTCGCAGTCCCCAACCTCCAATCACTCACCAACCTCCTGTCCTCCAATTTGTCCTGGTTATCGCTGGATGTGTCTGCGGCGTTTTATCATATTCCTCTTCATCCTGCTGCTATGCCTCATCTTCTTATTGGTTCTTCTGGATTATCAAGGTATGGCACCCGTTTGTCCTCTAATTCCAGGATCAACAACAACCAGTACGGGACCATGCAAAACCTGCACGACTCCTGCTCAAGGCAACTCTATGTTTCCCTCATGTTGCTGTACAAAACCTACGGATGGAAATTGCACCTGTATTCCCATCCCATCGTCCTGGGCTTTCGCAAAATACCTATGGGAGTGGGCCTCAGTCCGTTTCTCTTGGCTCAGTTTACTAGTGCCATTTGTTCAGTGGTTCGTAGGG |
| 105P | 5'-ACAAGAATCCTCACAATACCGCAGAGTCTAGACTCGTGGTGGACTTCTCTCAATTTTCTAGGGGGATCTCCCGTGTGTCTTGGCCAAAATTCGCAGTCCCCAACCTCCAATCACTCACCAACCTCCTGTCCTCCAATTTGTCCTGGTTATCGCTGGATGTGTCTGCGGCGTTTTATCATATTCCTCTTCATCCTGCTGCTATGCCTCATCTTCTTATTGGTTCTTCTGGATTATCAAGGTATGTTGGCAGTTTGTCCTCTAATTCCAGGATCAACAACAACCAGTACGGGACCATGCAAAACCTGCACGACTCCTGCTCAAGGCAACTCTATGTTTCCCTCATGTTGCTGTACAAAACCTACGGATGGAAATTGCACCTGTATTCCCATCCCATCGTCCTGGGCTTTCGCAAAATACCTATGGGAGTGGGCCTCAGTCCGTTTCTCTTGGCTCAGTTTACTAGTGCCATTTGTTCAGTGGTTCGTAGGG |
| 106V | 5'-ACAAGAATCCTCACAATACCGCAGAGTCTAGACTCGTGGTGGACTTCTCTCAATTTTCTAGGGGGATCTCCCGTGTGTCTTGGCCAAAATTCGCAGTCCCCAACCTCCAATCACTCACCAACCTCCTGTCCTCCAATTTGTCCTGGTTATCGCTGGATGTGTCTGCGGCGTTTTATCATATTCCTCTTCATCCTGCTGCTATGCCTCATCTTCTTATTGGTTCTTCTGGATTATCAAGGTATGTTGCCCGCATGTCCTCTAATTCCAGGATCAACAACAACCAGTACGGGACCATGCAAAACCTGCACGACTCCTGCTCAAGGCAACTCTATGTTTCCCTCATGTTGCTGTACAAAACCTACGGATGGAAATTGCACCTGTATTCCCATCCCATCGTCCTGGGCTTTCGCAAAATACCTATGGGAGTGGGCCTCAGTCCGTTTCTCTTGGCTCAGTTTACTAGTGCCATTTGTTCAGTGGTTCGTAGGG |
| 108P | 5'-ACAAGAATCCTCACAATACCGCAGAGTCTAGACTCGTGGTGGACTTCTCTCAATTTTCTAGGGGGATCTCCCGTGTGTCTTGGCCAAAATTCGCAGTCCCCAACCTCCAATCACTCACCAACCTCCTGTCCTCCAATTTGTCCTGGTTATCGCTGGATGTGTCTGCGGCGTTTTATCATATTCCTCTTCATCCTGCTGCTATGCCTCATCTTCTTATTGGTTCTTCTGGATTATCAAGGTATGTTGCCCGTTTGTGCACTAATTCCAGGATCAACAACAACCAGTACGGGACCATGCAAAACCTGCACGACTCCTGCTCAAGGCAACTCTATGTTTCCCTCATGTTGCTGTACAAAACCTACGGATGGAAATTGCACCTGTATTCCCATCCCATCGTCCTGGGCTTTCGCAAAATACCTATGGGAGTGGGCCTCAGTCCGTTTCTCTTGGCTCAGTTTACTAGTGCCATTTGTTCAGTGGTTCGTAGGG |
| 109L | 5'-ACAAGAATCCTCACAATACCGCAGAGTCTAGACTCGTGGTGGACTTCTCTCAATTTTCTAGGGGGATCTCCCGTGTGTCTTGGCCAAAATTCGCAGTCCCCAACCTCCAATCACTCACCAACCTCCTGTCCTCCAATTTGTCCTGGTTATCGCTGGATGTGTCTGCGGCGTTTTATCATATTCCTCTTCATCCTGCTGCTATGCCTCATCTTCTTATTGGTTCTTCTGGATTATCAAGGTATGTTGCCCGTTTGTCCTGCAATTCCAGGATCAACAACAACCAGTACGGGACCATGCAAAACCTGCACGACTCCTGCTCAAGGCAACTCTATGTTTCCCTCATGTTGCTGTACAAAACCTACGGATGGAAATTGCACCTGTATTCCCATCCCATCGTCCTGGGCTTTCGCAAAATACCTATGGGAGTGGGCCTCAGTCCGTTTCTCTTGGCTCAGTTTACTAGTGCCATTTGTTCAGTGGTTCGTAGGG |
| 110I | 5'-ACAAGAATCCTCACAATACCGCAGAGTCTAGACTCGTGGTGGACTTCTCTCAATTTTCTAGGGGGATCTCCCGTGTGTCTTGGCCAAAATTCGCAGTCCCCAACCTCCAATCACTCACCAACCTCCTGTCCTCCAATTTGTCCTGGTTATCGCTGGATGTGTCTGCGGCGTTTTATCATATTCCTCTTCATCCTGCTGCTATGCCTCATCTTCTTATTGGTTCTTCTGGATTATCAAGGTATGTTGCCCGTTTGTCCTCTAGCACCAGGATCAACAACAACCAGTACGGGACCATGCAAAACCTGCACGACTCCTGCTCAAGGCAACTCTATGTTTCCCTCATGTTGCTGTACAAAACCTACGGATGGAAATTGCACCTGTATTCCCATCCCATCGTCCTGGGCTTTCGCAAAATACCTATGGGAGTGGGCCTCAGTCCGTTTCTCTTGGCTCAGTTTACTAGTGCCATTTGTTCAGTGGTTCGTAGGG |
| 111P | 5'-ACAAGAATCCTCACAATACCGCAGAGTCTAGACTCGTGGTGGACTTCTCTCAATTTTCTAGGGGGATCTCCCGTGTGTCTTGGCCAAAATTCGCAGTCCCCAACCTCCAATCACTCACCAACCTCCTGTCCTCCAATTTGTCCTGGTTATCGCTGGATGTGTCTGCGGCGTTTTATCATATTCCTCTTCATCCTGCTGCTATGCCTCATCTTCTTATTGGTTCTTCTGGATTATCAAGGTATGTTGCCCGTTTGTCCTCTAATTGCAGGATCAACAACAACCAGTACGGGACCATGCAAAACCTGCACGACTCCTGCTCAAGGCAACTCTATGTTTCCCTCATGTTGCTGTACAAAACCTACGGATGGAAATTGCACCTGTATTCCCATCCCATCGTCCTGGGCTTTCGCAAAATACCTATGGGAGTGGGCCTCAGTCCGTTTCTCTTGGCTCAGTTTACTAGTGCCATTTGTTCAGTGGTTCGTAGGG |
| 112G | 5'-ACAAGAATCCTCACAATACCGCAGAGTCTAGACTCGTGGTGGACTTCTCTCAATTTTCTAGGGGGATCTCCCGTGTGTCTTGGCCAAAATTCGCAGTCCCCAACCTCCAATCACTCACCAACCTCCTGTCCTCCAATTTGTCCTGGTTATCGCTGGATGTGTCTGCGGCGTTTTATCATATTCCTCTTCATCCTGCTGCTATGCCTCATCTTCTTATTGGTTCTTCTGGATTATCAAGGTATGTTGCCCGTTTGTCCTCTAATTCCAGCATCAACAACAACCAGTACGGGACCATGCAAAACCTGCACGACTCCTGCTCAAGGCAACTCTATGTTTCCCTCATGTTGCTGTACAAAACCTACGGATGGAAATTGCACCTGTATTCCCATCCCATCGTCCTGGGCTTTCGCAAAATACCTATGGGAGTGGGCCTCAGTCCGTTTCTCTTGGCTCAGTTTACTAGTGCCATTTGTTCAGTGGTTCGTAGGG |
| 113S | 5'-ACAAGAATCCTCACAATACCGCAGAGTCTAGACTCGTGGTGGACTTCTCTCAATTTTCTAGGGGGATCTCCCGTGTGTCTTGGCCAAAATTCGCAGTCCCCAACCTCCAATCACTCACCAACCTCCTGTCCTCCAATTTGTCCTGGTTATCGCTGGATGTGTCTGCGGCGTTTTATCATATTCCTCTTCATCCTGCTGCTATGCCTCATCTTCTTATTGGTTCTTCTGGATTATCAAGGTATGTTGCCCGTTTGTCCTCTAATTCCAGGAGCAACAACAACCAGTACGGGACCATGCAAAACCTGCACGACTCCTGCTCAAGGCAACTCTATGTTTCCCTCATGTTGCTGTACAAAACCTACGGATGGAAATTGCACCTGTATTCCCATCCCATCGTCCTGGGCTTTCGCAAAATACCTATGGGAGTGGGCCTCAGTCCGTTTCTCTTGGCTCAGTTTACTAGTGCCATTTGTTCAGTGGTTCGTAGGG |
| 114T | 5'-ACAAGAATCCTCACAATACCGCAGAGTCTAGACTCGTGGTGGACTTCTCTCAATTTTCTAGGGGGATCTCCCGTGTGTCTTGGCCAAAATTCGCAGTCCCCAACCTCCAATCACTCACCAACCTCCTGTCCTCCAATTTGTCCTGGTTATCGCTGGATGTGTCTGCGGCGTTTTATCATATTCCTCTTCATCCTGCTGCTATGCCTCATCTTCTTATTGGTTCTTCTGGATTATCAAGGTATGTTGCCCGTTTGTCCTCTAATTCCAGGATCAGCAACAACCAGTACGGGACCATGCAAAACCTGCACGACTCCTGCTCAAGGCAACTCTATGTTTCCCTCATGTTGCTGTACAAAACCTACGGATGGAAATTGCACCTGTATTCCCATCCCATCGTCCTGGGCTTTCGCAAAATACCTATGGGAGTGGGCCTCAGTCCGTTTCTCTTGGCTCAGTTTACTAGTGCCATTTGTTCAGTGGTTCGTAGGG |
| 115T | 5'-ACAAGAATCCTCACAATACCGCAGAGTCTAGACTCGTGGTGGACTTCTCTCAATTTTCTAGGGGGATCTCCCGTGTGTCTTGGCCAAAATTCGCAGTCCCCAACCTCCAATCACTCACCAACCTCCTGTCCTCCAATTTGTCCTGGTTATCGCTGGATGTGTCTGCGGCGTTTTATCATATTCCTCTTCATCCTGCTGCTATGCCTCATCTTCTTATTGGTTCTTCTGGATTATCAAGGTATGTTGCCCGTTTGTCCTCTAATTCCAGGATCAACAGCAACCAGTACGGGACCATGCAAAACCTGCACGACTCCTGCTCAAGGCAACTCTATGTTTCCCTCATGTTGCTGTACAAAACCTACGGATGGAAATTGCACCTGTATTCCCATCCCATCGTCCTGGGCTTTCGCAAAATACCTATGGGAGTGGGCCTCAGTCCGTTTCTCTTGGCTCAGTTTACTAGTGCCATTTGTTCAGTGGTTCGTAGGG |
| 116T | 5'-ACAAGAATCCTCACAATACCGCAGAGTCTAGACTCGTGGTGGACTTCTCTCAATTTTCTAGGGGGATCTCCCGTGTGTCTTGGCCAAAATTCGCAGTCCCCAACCTCCAATCACTCACCAACCTCCTGTCCTCCAATTTGTCCTGGTTATCGCTGGATGTGTCTGCGGCGTTTTATCATATTCCTCTTCATCCTGCTGCTATGCCTCATCTTCTTATTGGTTCTTCTGGATTATCAAGGTATGTTGCCCGTTTGTCCTCTAATTCCAGGATCAACAACAGCAAGTACGGGACCATGCAAAACCTGCACGACTCCTGCTCAAGGCAACTCTATGTTTCCCTCATGTTGCTGTACAAAACCTACGGATGGAAATTGCACCTGTATTCCCATCCCATCGTCCTGGGCTTTCGCAAAATACCTATGGGAGTGGGCCTCAGTCCGTTTCTCTTGGCTCAGTTTACTAGTGCCATTTGTTCAGTGGTTCGTAGGG |
| 117S | 5'-ACAAGAATCCTCACAATACCGCAGAGTCTAGACTCGTGGTGGACTTCTCTCAATTTTCTAGGGGGATCTCCCGTGTGTCTTGGCCAAAATTCGCAGTCCCCAACCTCCAATCACTCACCAACCTCCTGTCCTCCAATTTGTCCTGGTTATCGCTGGATGTGTCTGCGGCGTTTTATCATATTCCTCTTCATCCTGCTGCTATGCCTCATCTTCTTATTGGTTCTTCTGGATTATCAAGGTATGTTGCCCGTTTGTCCTCTAATTCCAGGATCAACAACAACCGCAACGGGACCATGCAAAACCTGCACGACTCCTGCTCAAGGCAACTCTATGTTTCCCTCATGTTGCTGTACAAAACCTACGGATGGAAATTGCACCTGTATTCCCATCCCATCGTCCTGGGCTTTCGCAAAATACCTATGGGAGTGGGCCTCAGTCCGTTTCTCTTGGCTCAGTTTACTAGTGCCATTTGTTCAGTGGTTCGTAGGG |
| 118T | 5'-ACAAGAATCCTCACAATACCGCAGAGTCTAGACTCGTGGTGGACTTCTCTCAATTTTCTAGGGGGATCTCCCGTGTGTCTTGGCCAAAATTCGCAGTCCCCAACCTCCAATCACTCACCAACCTCCTGTCCTCCAATTTGTCCTGGTTATCGCTGGATGTGTCTGCGGCGTTTTATCATATTCCTCTTCATCCTGCTGCTATGCCTCATCTTCTTATTGGTTCTTCTGGATTATCAAGGTATGTTGCCCGTTTGTCCTCTAATTCCAGGATCAACAACAACCAGTGCAGGACCATGCAAAACCTGCACGACTCCTGCTCAAGGCAACTCTATGTTTCCCTCATGTTGCTGTACAAAACCTACGGATGGAAATTGCACCTGTATTCCCATCCCATCGTCCTGGGCTTTCGCAAAATACCTATGGGAGTGGGCCTCAGTCCGTTTCTCTTGGCTCAGTTTACTAGTGCCATTTGTTCAGTGGTTCGTAGGG |
| 119G | 5'-ACAAGAATCCTCACAATACCGCAGAGTCTAGACTCGTGGTGGACTTCTCTCAATTTTCTAGGGGGATCTCCCGTGTGTCTTGGCCAAAATTCGCAGTCCCCAACCTCCAATCACTCACCAACCTCCTGTCCTCCAATTTGTCCTGGTTATCGCTGGATGTGTCTGCGGCGTTTTATCATATTCCTCTTCATCCTGCTGCTATGCCTCATCTTCTTATTGGTTCTTCTGGATTATCAAGGTATGTTGCCCGTTTGTCCTCTAATTCCAGGATCAACAACAACCAGTACGGCACCATGCAAAACCTGCACGACTCCTGCTCAAGGCAACTCTATGTTTCCCTCATGTTGCTGTACAAAACCTACGGATGGAAATTGCACCTGTATTCCCATCCCATCGTCCTGGGCTTTCGCAAAATACCTATGGGAGTGGGCCTCAGTCCGTTTCTCTTGGCTCAGTTTACTAGTGCCATTTGTTCAGTGGTTCGTAGGG |
| 120P | 5'-ACAAGAATCCTCACAATACCGCAGAGTCTAGACTCGTGGTGGACTTCTCTCAATTTTCTAGGGGGATCTCCCGTGTGTCTTGGCCAAAATTCGCAGTCCCCAACCTCCAATCACTCACCAACCTCCTGTCCTCCAATTTGTCCTGGTTATCGCTGGATGTGTCTGCGGCGTTTTATCATATTCCTCTTCATCCTGCTGCTATGCCTCATCTTCTTATTGGTTCTTCTGGATTATCAAGGTATGTTGCCCGTTTGTCCTCTAATTCCAGGATCAACAACAACCAGTACGGGAGCATGCAAAACCTGCACGACTCCTGCTCAAGGCAACTCTATGTTTCCCTCATGTTGCTGTACAAAACCTACGGATGGAAATTGCACCTGTATTCCCATCCCATCGTCCTGGGCTTTCGCAAAATACCTATGGGAGTGGGCCTCAGTCCGTTTCTCTTGGCTCAGTTTACTAGTGCCATTTGTTCAGTGGTTCGTAGGG |
| 122K | 5'-ACAAGAATCCTCACAATACCGCAGAGTCTAGACTCGTGGTGGACTTCTCTCAATTTTCTAGGGGGATCTCCCGTGTGTCTTGGCCAAAATTCGCAGTCCCCAACCTCCAATCACTCACCAACCTCCTGTCCTCCAATTTGTCCTGGTTATCGCTGGATGTGTCTGCGGCGTTTTATCATATTCCTCTTCATCCTGCTGCTATGCCTCATCTTCTTATTGGTTCTTCTGGATTATCAAGGTATGTTGCCCGTTTGTCCTCTAATTCCAGGATCAACAACAACCAGTACGGGACCATGCGCAACCTGCACGACTCCTGCTCAAGGCAACTCTATGTTTCCCTCATGTTGCTGTACAAAACCTACGGATGGAAATTGCACCTGTATTCCCATCCCATCGTCCTGGGCTTTCGCAAAATACCTATGGGAGTGGGCCTCAGTCCGTTTCTCTTGGCTCAGTTTACTAGTGCCATTTGTTCAGTGGTTCGTAGGG |
| 123T | 5'-ACAAGAATCCTCACAATACCGCAGAGTCTAGACTCGTGGTGGACTTCTCTCAATTTTCTAGGGGGATCTCCCGTGTGTCTTGGCCAAAATTCGCAGTCCCCAACCTCCAATCACTCACCAACCTCCTGTCCTCCAATTTGTCCTGGTTATCGCTGGATGTGTCTGCGGCGTTTTATCATATTCCTCTTCATCCTGCTGCTATGCCTCATCTTCTTATTGGTTCTTCTGGATTATCAAGGTATGTTGCCCGTTTGTCCTCTAATTCCAGGATCAACAACAACCAGTACGGGACCATGCAAAGCATGCACGACTCCTGCTCAAGGCAACTCTATGTTTCCCTCATGTTGCTGTACAAAACCTACGGATGGAAATTGCACCTGTATTCCCATCCCATCGTCCTGGGCTTTCGCAAAATACCTATGGGAGTGGGCCTCAGTCCGTTTCTCTTGGCTCAGTTTACTAGTGCCATTTGTTCAGTGGTTCGTAGGG |
| 125T | 5'-ACAAGAATCCTCACAATACCGCAGAGTCTAGACTCGTGGTGGACTTCTCTCAATTTTCTAGGGGGATCTCCCGTGTGTCTTGGCCAAAATTCGCAGTCCCCAACCTCCAATCACTCACCAACCTCCTGTCCTCCAATTTGTCCTGGTTATCGCTGGATGTGTCTGCGGCGTTTTATCATATTCCTCTTCATCCTGCTGCTATGCCTCATCTTCTTATTGGTTCTTCTGGATTATCAAGGTATGTTGCCCGTTTGTCCTCTAATTCCAGGATCAACAACAACCAGTACGGGACCATGCAAAACCTGCGCAACTCCTGCTCAAGGCAACTCTATGTTTCCCTCATGTTGCTGTACAAAACCTACGGATGGAAATTGCACCTGTATTCCCATCCCATCGTCCTGGGCTTTCGCAAAATACCTATGGGAGTGGGCCTCAGTCCGTTTCTCTTGGCTCAGTTTACTAGTGCCATTTGTTCAGTGGTTCGTAGGG |
| 126T | 5'-ACAAGAATCCTCACAATACCGCAGAGTCTAGACTCGTGGTGGACTTCTCTCAATTTTCTAGGGGGATCTCCCGTGTGTCTTGGCCAAAATTCGCAGTCCCCAACCTCCAATCACTCACCAACCTCCTGTCCTCCAATTTGTCCTGGTTATCGCTGGATGTGTCTGCGGCGTTTTATCATATTCCTCTTCATCCTGCTGCTATGCCTCATCTTCTTATTGGTTCTTCTGGATTATCAAGGTATGTTGCCCGTTTGTCCTCTAATTCCAGGATCAACAACAACCAGTACGGGACCATGCAAAACCTGCACGGCACCTGCTCAAGGCAACTCTATGTTTCCCTCATGTTGCTGTACAAAACCTACGGATGGAAATTGCACCTGTATTCCCATCCCATCGTCCTGGGCTTTCGCAAAATACCTATGGGAGTGGGCCTCAGTCCGTTTCTCTTGGCTCAGTTTACTAGTGCCATTTGTTCAGTGGTTCGTAGGG |
| 127P | 5'-ACAAGAATCCTCACAATACCGCAGAGTCTAGACTCGTGGTGGACTTCTCTCAATTTTCTAGGGGGATCTCCCGTGTGTCTTGGCCAAAATTCGCAGTCCCCAACCTCCAATCACTCACCAACCTCCTGTCCTCCAATTTGTCCTGGTTATCGCTGGATGTGTCTGCGGCGTTTTATCATATTCCTCTTCATCCTGCTGCTATGCCTCATCTTCTTATTGGTTCTTCTGGATTATCAAGGTATGTTGCCCGTTTGTCCTCTAATTCCAGGATCAACAACAACCAGTACGGGACCATGCAAAACCTGCACGACTGCAGCTCAAGGCAACTCTATGTTTCCCTCATGTTGCTGTACAAAACCTACGGATGGAAATTGCACCTGTATTCCCATCCCATCGTCCTGGGCTTTCGCAAAATACCTATGGGAGTGGGCCTCAGTCCGTTTCTCTTGGCTCAGTTTACTAGTGCCATTTGTTCAGTGGTTCGTAGGG |
| 129Q | 5'-ACAAGAATCCTCACAATACCGCAGAGTCTAGACTCGTGGTGGACTTCTCTCAATTTTCTAGGGGGATCTCCCGTGTGTCTTGGCCAAAATTCGCAGTCCCCAACCTCCAATCACTCACCAACCTCCTGTCCTCCAATTTGTCCTGGTTATCGCTGGATGTGTCTGCGGCGTTTTATCATATTCCTCTTCATCCTGCTGCTATGCCTCATCTTCTTATTGGTTCTTCTGGATTATCAAGGTATGTTGCCCGTTTGTCCTCTAATTCCAGGATCAACAACAACCAGTACGGGACCATGCAAAACCTGCACGACTCCTGCTGCAGGCAACTCTATGTTTCCCTCATGTTGCTGTACAAAACCTACGGATGGAAATTGCACCTGTATTCCCATCCCATCGTCCTGGGCTTTCGCAAAATACCTATGGGAGTGGGCCTCAGTCCGTTTCTCTTGGCTCAGTTTACTAGTGCCATTTGTTCAGTGGTTCGTAGGG |
| 130G | 5'-ACAAGAATCCTCACAATACCGCAGAGTCTAGACTCGTGGTGGACTTCTCTCAATTTTCTAGGGGGATCTCCCGTGTGTCTTGGCCAAAATTCGCAGTCCCCAACCTCCAATCACTCACCAACCTCCTGTCCTCCAATTTGTCCTGGTTATCGCTGGATGTGTCTGCGGCGTTTTATCATATTCCTCTTCATCCTGCTGCTATGCCTCATCTTCTTATTGGTTCTTCTGGATTATCAAGGTATGTTGCCCGTTTGTCCTCTAATTCCAGGATCAACAACAACCAGTACGGGACCATGCAAAACCTGCACGACTCCTGCTCAAGCAAACTCTATGTTTCCCTCATGTTGCTGTACAAAACCTACGGATGGAAATTGCACCTGTATTCCCATCCCATCGTCCTGGGCTTTCGCAAAATACCTATGGGAGTGGGCCTCAGTCCGTTTCTCTTGGCTCAGTTTACTAGTGCCATTTGTTCAGTGGTTCGTAGGG |
| 131N | 5'-ACAAGAATCCTCACAATACCGCAGAGTCTAGACTCGTGGTGGACTTCTCTCAATTTTCTAGGGGGATCTCCCGTGTGTCTTGGCCAAAATTCGCAGTCCCCAACCTCCAATCACTCACCAACCTCCTGTCCTCCAATTTGTCCTGGTTATCGCTGGATGTGTCTGCGGCGTTTTATCATATTCCTCTTCATCCTGCTGCTATGCCTCATCTTCTTATTGGTTCTTCTGGATTATCAAGGTATGTTGCCCGTTTGTCCTCTAATTCCAGGATCAACAACAACCAGTACGGGACCATGCAAAACCTGCACGACTCCTGCTCAAGGCGCATCTATGTTTCCCTCATGTTGCTGTACAAAACCTACGGATGGAAATTGCACCTGTATTCCCATCCCATCGTCCTGGGCTTTCGCAAAATACCTATGGGAGTGGGCCTCAGTCCGTTTCTCTTGGCTCAGTTTACTAGTGCCATTTGTTCAGTGGTTCGTAGGG |
| 132S | 5'-ACAAGAATCCTCACAATACCGCAGAGTCTAGACTCGTGGTGGACTTCTCTCAATTTTCTAGGGGGATCTCCCGTGTGTCTTGGCCAAAATTCGCAGTCCCCAACCTCCAATCACTCACCAACCTCCTGTCCTCCAATTTGTCCTGGTTATCGCTGGATGTGTCTGCGGCGTTTTATCATATTCCTCTTCATCCTGCTGCTATGCCTCATCTTCTTATTGGTTCTTCTGGATTATCAAGGTATGTTGCCCGTTTGTCCTCTAATTCCAGGATCAACAACAACCAGTACGGGACCATGCAAAACCTGCACGACTCCTGCTCAAGGCAACGCAATGTTTCCCTCATGTTGCTGTACAAAACCTACGGATGGAAATTGCACCTGTATTCCCATCCCATCGTCCTGGGCTTTCGCAAAATACCTATGGGAGTGGGCCTCAGTCCGTTTCTCTTGGCTCAGTTTACTAGTGCCATTTGTTCAGTGGTTCGTAGGG |
| 133M | 5'-ACAAGAATCCTCACAATACCGCAGAGTCTAGACTCGTGGTGGACTTCTCTCAATTTTCTAGGGGGATCTCCCGTGTGTCTTGGCCAAAATTCGCAGTCCCCAACCTCCAATCACTCACCAACCTCCTGTCCTCCAATTTGTCCTGGTTATCGCTGGATGTGTCTGCGGCGTTTTATCATATTCCTCTTCATCCTGCTGCTATGCCTCATCTTCTTATTGGTTCTTCTGGATTATCAAGGTATGTTGCCCGTTTGTCCTCTAATTCCAGGATCAACAACAACCAGTACGGGACCATGCAAAACCTGCACGACTCCTGCTCAAGGCAACTCTGCATTTCCCTCATGTTGCTGTACAAAACCTACGGATGGAAATTGCACCTGTATTCCCATCCCATCGTCCTGGGCTTTCGCAAAATACCTATGGGAGTGGGCCTCAGTCCGTTTCTCTTGGCTCAGTTTACTAGTGCCATTTGTTCAGTGGTTCGTAGGG |
| 134F | 5'-ACAAGAATCCTCACAATACCGCAGAGTCTAGACTCGTGGTGGACTTCTCTCAATTTTCTAGGGGGATCTCCCGTGTGTCTTGGCCAAAATTCGCAGTCCCCAACCTCCAATCACTCACCAACCTCCTGTCCTCCAATTTGTCCTGGTTATCGCTGGATGTGTCTGCGGCGTTTTATCATATTCCTCTTCATCCTGCTGCTATGCCTCATCTTCTTATTGGTTCTTCTGGATTATCAAGGTATGTTGCCCGTTTGTCCTCTAATTCCAGGATCAACAACAACCAGTACGGGACCATGCAAAACCTGCACGACTCCTGCTCAAGGCAACTCTATGGCACCCTCATGTTGCTGTACAAAACCTACGGATGGAAATTGCACCTGTATTCCCATCCCATCGTCCTGGGCTTTCGCAAAATACCTATGGGAGTGGGCCTCAGTCCGTTTCTCTTGGCTCAGTTTACTAGTGCCATTTGTTCAGTGGTTCGTAGGG |
| 135P | 5'-ACAAGAATCCTCACAATACCGCAGAGTCTAGACTCGTGGTGGACTTCTCTCAATTTTCTAGGGGGATCTCCCGTGTGTCTTGGCCAAAATTCGCAGTCCCCAACCTCCAATCACTCACCAACCTCCTGTCCTCCAATTTGTCCTGGTTATCGCTGGATGTGTCTGCGGCGTTTTATCATATTCCTCTTCATCCTGCTGCTATGCCTCATCTTCTTATTGGTTCTTCTGGATTATCAAGGTATGTTGCCCGTTTGTCCTCTAATTCCAGGATCAACAACAACCAGTACGGGACCATGCAAAACCTGCACGACTCCTGCTCAAGGCAACTCTATGTTTGCATCATGTTGCTGTACAAAACCTACGGATGGAAATTGCACCTGTATTCCCATCCCATCGTCCTGGGCTTTCGCAAAATACCTATGGGAGTGGGCCTCAGTCCGTTTCTCTTGGCTCAGTTTACTAGTGCCATTTGTTCAGTGGTTCGTAGGG |
| 136S | 5'-ACAAGAATCCTCACAATACCGCAGAGTCTAGACTCGTGGTGGACTTCTCTCAATTTTCTAGGGGGATCTCCCGTGTGTCTTGGCCAAAATTCGCAGTCCCCAACCTCCAATCACTCACCAACCTCCTGTCCTCCAATTTGTCCTGGTTATCGCTGGATGTGTCTGCGGCGTTTTATCATATTCCTCTTCATCCTGCTGCTATGCCTCATCTTCTTATTGGTTCTTCTGGATTATCAAGGTATGTTGCCCGTTTGTCCTCTAATTCCAGGATCAACAACAACCAGTACGGGACCATGCAAAACCTGCACGACTCCTGCTCAAGGCAACTCTATGTTTCCCGCATGTTGCTGTACAAAACCTACGGATGGAAATTGCACCTGTATTCCCATCCCATCGTCCTGGGCTTTCGCAAAATACCTATGGGAGTGGGCCTCAGTCCGTTTCTCTTGGCTCAGTTTACTAGTGCCATTTGTTCAGTGGTTCGTAGGG |
| 140T | 5'-ACAAGAATCCTCACAATACCGCAGAGTCTAGACTCGTGGTGGACTTCTCTCAATTTTCTAGGGGGATCTCCCGTGTGTCTTGGCCAAAATTCGCAGTCCCCAACCTCCAATCACTCACCAACCTCCTGTCCTCCAATTTGTCCTGGTTATCGCTGGATGTGTCTGCGGCGTTTTATCATATTCCTCTTCATCCTGCTGCTATGCCTCATCTTCTTATTGGTTCTTCTGGATTATCAAGGTATGTTGCCCGTTTGTCCTCTAATTCCAGGATCAACAACAACCAGTACGGGACCATGCAAAACCTGCACGACTCCTGCTCAAGGCAACTCTATGTTTCCCTCATGTTGCTGTGCAAAACCTACGGATGGAAATTGCACCTGTATTCCCATCCCATCGTCCTGGGCTTTCGCAAAATACCTATGGGAGTGGGCCTCAGTCCGTTTCTCTTGGCTCAGTTTACTAGTGCCATTTGTTCAGTGGTTCGTAGGG |
| 141K | 5'-ACAAGAATCCTCACAATACCGCAGAGTCTAGACTCGTGGTGGACTTCTCTCAATTTTCTAGGGGGATCTCCCGTGTGTCTTGGCCAAAATTCGCAGTCCCCAACCTCCAATCACTCACCAACCTCCTGTCCTCCAATTTGTCCTGGTTATCGCTGGATGTGTCTGCGGCGTTTTATCATATTCCTCTTCATCCTGCTGCTATGCCTCATCTTCTTATTGGTTCTTCTGGATTATCAAGGTATGTTGCCCGTTTGTCCTCTAATTCCAGGATCAACAACAACCAGTACGGGACCATGCAAAACCTGCACGACTCCTGCTCAAGGCAACTCTATGTTTCCCTCATGTTGCTGTACAGCACCTACGGATGGAAATTGCACCTGTATTCCCATCCCATCGTCCTGGGCTTTCGCAAAATACCTATGGGAGTGGGCCTCAGTCCGTTTCTCTTGGCTCAGTTTACTAGTGCCATTTGTTCAGTGGTTCGTAGGG |
| 142P | 5'-ACAAGAATCCTCACAATACCGCAGAGTCTAGACTCGTGGTGGACTTCTCTCAATTTTCTAGGGGGATCTCCCGTGTGTCTTGGCCAAAATTCGCAGTCCCCAACCTCCAATCACTCACCAACCTCCTGTCCTCCAATTTGTCCTGGTTATCGCTGGATGTGTCTGCGGCGTTTTATCATATTCCTCTTCATCCTGCTGCTATGCCTCATCTTCTTATTGGTTCTTCTGGATTATCAAGGTATGTTGCCCGTTTGTCCTCTAATTCCAGGATCAACAACAACCAGTACGGGACCATGCAAAACCTGCACGACTCCTGCTCAAGGCAACTCTATGTTTCCCTCATGTTGCTGTACAAAAGCAACGGATGGAAATTGCACCTGTATTCCCATCCCATCGTCCTGGGCTTTCGCAAAATACCTATGGGAGTGGGCCTCAGTCCGTTTCTCTTGGCTCAGTTTACTAGTGCCATTTGTTCAGTGGTTCGTAGGG |
| 143T | 5'-ACAAGAATCCTCACAATACCGCAGAGTCTAGACTCGTGGTGGACTTCTCTCAATTTTCTAGGGGGATCTCCCGTGTGTCTTGGCCAAAATTCGCAGTCCCCAACCTCCAATCACTCACCAACCTCCTGTCCTCCAATTTGTCCTGGTTATCGCTGGATGTGTCTGCGGCGTTTTATCATATTCCTCTTCATCCTGCTGCTATGCCTCATCTTCTTATTGGTTCTTCTGGATTATCAAGGTATGTTGCCCGTTTGTCCTCTAATTCCAGGATCAACAACAACCAGTACGGGACCATGCAAAACCTGCACGACTCCTGCTCAAGGCAACTCTATGTTTCCCTCATGTTGCTGTACAAAACCTGCAGATGGAAATTGCACCTGTATTCCCATCCCATCGTCCTGGGCTTTCGCAAAATACCTATGGGAGTGGGCCTCAGTCCGTTTCTCTTGGCTCAGTTTACTAGTGCCATTTGTTCAGTGGTTCGTAGGG |
| 144D | 5'-ACAAGAATCCTCACAATACCGCAGAGTCTAGACTCGTGGTGGACTTCTCTCAATTTTCTAGGGGGATCTCCCGTGTGTCTTGGCCAAAATTCGCAGTCCCCAACCTCCAATCACTCACCAACCTCCTGTCCTCCAATTTGTCCTGGTTATCGCTGGATGTGTCTGCGGCGTTTTATCATATTCCTCTTCATCCTGCTGCTATGCCTCATCTTCTTATTGGTTCTTCTGGATTATCAAGGTATGTTGCCCGTTTGTCCTCTAATTCCAGGATCAACAACAACCAGTACGGGACCATGCAAAACCTGCACGACTCCTGCTCAAGGCAACTCTATGTTTCCCTCATGTTGCTGTACAAAACCTACGGCAGGAAATTGCACCTGTATTCCCATCCCATCGTCCTGGGCTTTCGCAAAATACCTATGGGAGTGGGCCTCAGTCCGTTTCTCTTGGCTCAGTTTACTAGTGCCATTTGTTCAGTGGTTCGTAGGG |
| 145G | 5'-ACAAGAATCCTCACAATACCGCAGAGTCTAGACTCGTGGTGGACTTCTCTCAATTTTCTAGGGGGATCTCCCGTGTGTCTTGGCCAAAATTCGCAGTCCCCAACCTCCAATCACTCACCAACCTCCTGTCCTCCAATTTGTCCTGGTTATCGCTGGATGTGTCTGCGGCGTTTTATCATATTCCTCTTCATCCTGCTGCTATGCCTCATCTTCTTATTGGTTCTTCTGGATTATCAAGGTATGTTGCCCGTTTGTCCTCTAATTCCAGGATCAACAACAACCAGTACGGGACCATGCAAAACCTGCACGACTCCTGCTCAAGGCAACTCTATGTTTCCCTCATGTTGCTGTACAAAACCTACGGATGCAAATTGCACCTGTATTCCCATCCCATCGTCCTGGGCTTTCGCAAAATACCTATGGGAGTGGGCCTCAGTCCGTTTCTCTTGGCTCAGTTTACTAGTGCCATTTGTTCAGTGGTTCGTAGGG |
| 146N | 5'-ACAAGAATCCTCACAATACCGCAGAGTCTAGACTCGTGGTGGACTTCTCTCAATTTTCTAGGGGGATCTCCCGTGTGTCTTGGCCAAAATTCGCAGTCCCCAACCTCCAATCACTCACCAACCTCCTGTCCTCCAATTTGTCCTGGTTATCGCTGGATGTGTCTGCGGCGTTTTATCATATTCCTCTTCATCCTGCTGCTATGCCTCATCTTCTTATTGGTTCTTCTGGATTATCAAGGTATGTTGCCCGTTTGTCCTCTAATTCCAGGATCAACAACAACCAGTACGGGACCATGCAAAACCTGCACGACTCCTGCTCAAGGCAACTCTATGTTTCCCTCATGTTGCTGTACAAAACCTACGGATGGAGCATGCACCTGTATTCCCATCCCATCGTCCTGGGCTTTCGCAAAATACCTATGGGAGTGGGCCTCAGTCCGTTTCTCTTGGCTCAGTTTACTAGTGCCATTTGTTCAGTGGTTCGTAGGG |
| 148T | 5'-ACAAGAATCCTCACAATACCGCAGAGTCTAGACTCGTGGTGGACTTCTCTCAATTTTCTAGGGGGATCTCCCGTGTGTCTTGGCCAAAATTCGCAGTCCCCAACCTCCAATCACTCACCAACCTCCTGTCCTCCAATTTGTCCTGGTTATCGCTGGATGTGTCTGCGGCGTTTTATCATATTCCTCTTCATCCTGCTGCTATGCCTCATCTTCTTATTGGTTCTTCTGGATTATCAAGGTATGTTGCCCGTTTGTCCTCTAATTCCAGGATCAACAACAACCAGTACGGGACCATGCAAAACCTGCACGACTCCTGCTCAAGGCAACTCTATGTTTCCCTCATGTTGCTGTACAAAACCTACGGATGGAAATTGCGCATGTATTCCCATCCCATCGTCCTGGGCTTTCGCAAAATACCTATGGGAGTGGGCCTCAGTCCGTTTCTCTTGGCTCAGTTTACTAGTGCCATTTGTTCAGTGGTTCGTAGGG |
| 150I | 5'-ACAAGAATCCTCACAATACCGCAGAGTCTAGACTCGTGGTGGACTTCTCTCAATTTTCTAGGGGGATCTCCCGTGTGTCTTGGCCAAAATTCGCAGTCCCCAACCTCCAATCACTCACCAACCTCCTGTCCTCCAATTTGTCCTGGTTATCGCTGGATGTGTCTGCGGCGTTTTATCATATTCCTCTTCATCCTGCTGCTATGCCTCATCTTCTTATTGGTTCTTCTGGATTATCAAGGTATGTTGCCCGTTTGTCCTCTAATTCCAGGATCAACAACAACCAGTACGGGACCATGCAAAACCTGCACGACTCCTGCTCAAGGCAACTCTATGTTTCCCTCATGTTGCTGTACAAAACCTACGGATGGAAATTGCACCTGTGCACCCATCCCATCGTCCTGGGCTTTCGCAAAATACCTATGGGAGTGGGCCTCAGTCCGTTTCTCTTGGCTCAGTTTACTAGTGCCATTTGTTCAGTGGTTCGTAGGG |
| 151P | 5'-ACAAGAATCCTCACAATACCGCAGAGTCTAGACTCGTGGTGGACTTCTCTCAATTTTCTAGGGGGATCTCCCGTGTGTCTTGGCCAAAATTCGCAGTCCCCAACCTCCAATCACTCACCAACCTCCTGTCCTCCAATTTGTCCTGGTTATCGCTGGATGTGTCTGCGGCGTTTTATCATATTCCTCTTCATCCTGCTGCTATGCCTCATCTTCTTATTGGTTCTTCTGGATTATCAAGGTATGTTGCCCGTTTGTCCTCTAATTCCAGGATCAACAACAACCAGTACGGGACCATGCAAAACCTGCACGACTCCTGCTCAAGGCAACTCTATGTTTCCCTCATGTTGCTGTACAAAACCTACGGATGGAAATTGCACCTGTATTGCAATCCCATCGTCCTGGGCTTTCGCAAAATACCTATGGGAGTGGGCCTCAGTCCGTTTCTCTTGGCTCAGTTTACTAGTGCCATTTGTTCAGTGGTTCGTAGGG |
| 152I | 5'-ACAAGAATCCTCACAATACCGCAGAGTCTAGACTCGTGGTGGACTTCTCTCAATTTTCTAGGGGGATCTCCCGTGTGTCTTGGCCAAAATTCGCAGTCCCCAACCTCCAATCACTCACCAACCTCCTGTCCTCCAATTTGTCCTGGTTATCGCTGGATGTGTCTGCGGCGTTTTATCATATTCCTCTTCATCCTGCTGCTATGCCTCATCTTCTTATTGGTTCTTCTGGATTATCAAGGTATGTTGCCCGTTTGTCCTCTAATTCCAGGATCAACAACAACCAGTACGGGACCATGCAAAACCTGCACGACTCCTGCTCAAGGCAACTCTATGTTTCCCTCATGTTGCTGTACAAAACCTACGGATGGAAATTGCACCTGTATTCCCGCACCATCGTCCTGGGCTTTCGCAAAATACCTATGGGAGTGGGCCTCAGTCCGTTTCTCTTGGCTCAGTTTACTAGTGCCATTTGTTCAGTGGTTCGTAGGG |
| 153P | 5'-ACAAGAATCCTCACAATACCGCAGAGTCTAGACTCGTGGTGGACTTCTCTCAATTTTCTAGGGGGATCTCCCGTGTGTCTTGGCCAAAATTCGCAGTCCCCAACCTCCAATCACTCACCAACCTCCTGTCCTCCAATTTGTCCTGGTTATCGCTGGATGTGTCTGCGGCGTTTTATCATATTCCTCTTCATCCTGCTGCTATGCCTCATCTTCTTATTGGTTCTTCTGGATTATCAAGGTATGTTGCCCGTTTGTCCTCTAATTCCAGGATCAACAACAACCAGTACGGGACCATGCAAAACCTGCACGACTCCTGCTCAAGGCAACTCTATGTTTCCCTCATGTTGCTGTACAAAACCTACGGATGGAAATTGCACCTGTATTCCCATCGCATCGTCCTGGGCTTTCGCAAAATACCTATGGGAGTGGGCCTCAGTCCGTTTCTCTTGGCTCAGTTTACTAGTGCCATTTGTTCAGTGGTTCGTAGGG |
| 154S | 5'-ACAAGAATCCTCACAATACCGCAGAGTCTAGACTCGTGGTGGACTTCTCTCAATTTTCTAGGGGGATCTCCCGTGTGTCTTGGCCAAAATTCGCAGTCCCCAACCTCCAATCACTCACCAACCTCCTGTCCTCCAATTTGTCCTGGTTATCGCTGGATGTGTCTGCGGCGTTTTATCATATTCCTCTTCATCCTGCTGCTATGCCTCATCTTCTTATTGGTTCTTCTGGATTATCAAGGTATGTTGCCCGTTTGTCCTCTAATTCCAGGATCAACAACAACCAGTACGGGACCATGCAAAACCTGCACGACTCCTGCTCAAGGCAACTCTATGTTTCCCTCATGTTGCTGTACAAAACCTACGGATGGAAATTGCACCTGTATTCCCATCCCAGCATCCTGGGCTTTCGCAAAATACCTATGGGAGTGGGCCTCAGTCCGTTTCTCTTGGCTCAGTTTACTAGTGCCATTTGTTCAGTGGTTCGTAGGG |
| 155S | 5'-ACAAGAATCCTCACAATACCGCAGAGTCTAGACTCGTGGTGGACTTCTCTCAATTTTCTAGGGGGATCTCCCGTGTGTCTTGGCCAAAATTCGCAGTCCCCAACCTCCAATCACTCACCAACCTCCTGTCCTCCAATTTGTCCTGGTTATCGCTGGATGTGTCTGCGGCGTTTTATCATATTCCTCTTCATCCTGCTGCTATGCCTCATCTTCTTATTGGTTCTTCTGGATTATCAAGGTATGTTGCCCGTTTGTCCTCTAATTCCAGGATCAACAACAACCAGTACGGGACCATGCAAAACCTGCACGACTCCTGCTCAAGGCAACTCTATGTTTCCCTCATGTTGCTGTACAAAACCTACGGATGGAAATTGCACCTGTATTCCCATCCCATCGGCATGGGCTTTCGCAAAATACCTATGGGAGTGGGCCTCAGTCCGTTTCTCTTGGCTCAGTTTACTAGTGCCATTTGTTCAGTGGTTCGTAGGG |
| 156W | 5'-ACAAGAATCCTCACAATACCGCAGAGTCTAGACTCGTGGTGGACTTCTCTCAATTTTCTAGGGGGATCTCCCGTGTGTCTTGGCCAAAATTCGCAGTCCCCAACCTCCAATCACTCACCAACCTCCTGTCCTCCAATTTGTCCTGGTTATCGCTGGATGTGTCTGCGGCGTTTTATCATATTCCTCTTCATCCTGCTGCTATGCCTCATCTTCTTATTGGTTCTTCTGGATTATCAAGGTATGTTGCCCGTTTGTCCTCTAATTCCAGGATCAACAACAACCAGTACGGGACCATGCAAAACCTGCACGACTCCTGCTCAAGGCAACTCTATGTTTCCCTCATGTTGCTGTACAAAACCTACGGATGGAAATTGCACCTGTATTCCCATCCCATCGTCCGCAGCTTTCGCAAAATACCTATGGGAGTGGGCCTCAGTCCGTTTCTCTTGGCTCAGTTTACTAGTGCCATTTGTTCAGTGGTTCGTAGGG |
| 158F | 5'-ACAAGAATCCTCACAATACCGCAGAGTCTAGACTCGTGGTGGACTTCTCTCAATTTTCTAGGGGGATCTCCCGTGTGTCTTGGCCAAAATTCGCAGTCCCCAACCTCCAATCACTCACCAACCTCCTGTCCTCCAATTTGTCCTGGTTATCGCTGGATGTGTCTGCGGCGTTTTATCATATTCCTCTTCATCCTGCTGCTATGCCTCATCTTCTTATTGGTTCTTCTGGATTATCAAGGTATGTTGCCCGTTTGTCCTCTAATTCCAGGATCAACAACAACCAGTACGGGACCATGCAAAACCTGCACGACTCCTGCTCAAGGCAACTCTATGTTTCCCTCATGTTGCTGTACAAAACCTACGGATGGAAATTGCACCTGTATTCCCATCCCATCGTCCTGGGCTGCAGCAAAATACCTATGGGAGTGGGCCTCAGTCCGTTTCTCTTGGCTCAGTTTACTAGTGCCATTTGTTCAGTGGTTCGTAGGG |
| 160K | 5'-ACAAGAATCCTCACAATACCGCAGAGTCTAGACTCGTGGTGGACTTCTCTCAATTTTCTAGGGGGATCTCCCGTGTGTCTTGGCCAAAATTCGCAGTCCCCAACCTCCAATCACTCACCAACCTCCTGTCCTCCAATTTGTCCTGGTTATCGCTGGATGTGTCTGCGGCGTTTTATCATATTCCTCTTCATCCTGCTGCTATGCCTCATCTTCTTATTGGTTCTTCTGGATTATCAAGGTATGTTGCCCGTTTGTCCTCTAATTCCAGGATCAACAACAACCAGTACGGGACCATGCAAAACCTGCACGACTCCTGCTCAAGGCAACTCTATGTTTCCCTCATGTTGCTGTACAAAACCTACGGATGGAAATTGCACCTGTATTCCCATCCCATCGTCCTGGGCTTTCGCAGCATACCTATGGGAGTGGGCCTCAGTCCGTTTCTCTTGGCTCAGTTTACTAGTGCCATTTGTTCAGTGGTTCGTAGGG |
| 161Y | 5'-ACAAGAATCCTCACAATACCGCAGAGTCTAGACTCGTGGTGGACTTCTCTCAATTTTCTAGGGGGATCTCCCGTGTGTCTTGGCCAAAATTCGCAGTCCCCAACCTCCAATCACTCACCAACCTCCTGTCCTCCAATTTGTCCTGGTTATCGCTGGATGTGTCTGCGGCGTTTTATCATATTCCTCTTCATCCTGCTGCTATGCCTCATCTTCTTATTGGTTCTTCTGGATTATCAAGGTATGTTGCCCGTTTGTCCTCTAATTCCAGGATCAACAACAACCAGTACGGGACCATGCAAAACCTGCACGACTCCTGCTCAAGGCAACTCTATGTTTCCCTCATGTTGCTGTACAAAACCTACGGATGGAAATTGCACCTGTATTCCCATCCCATCGTCCTGGGCTTTCGCAAAAGCACTATGGGAGTGGGCCTCAGTCCGTTTCTCTTGGCTCAGTTTACTAGTGCCATTTGTTCAGTGGTTCGTAGGG |
| 162L | 5'-ACAAGAATCCTCACAATACCGCAGAGTCTAGACTCGTGGTGGACTTCTCTCAATTTTCTAGGGGGATCTCCCGTGTGTCTTGGCCAAAATTCGCAGTCCCCAACCTCCAATCACTCACCAACCTCCTGTCCTCCAATTTGTCCTGGTTATCGCTGGATGTGTCTGCGGCGTTTTATCATATTCCTCTTCATCCTGCTGCTATGCCTCATCTTCTTATTGGTTCTTCTGGATTATCAAGGTATGTTGCCCGTTTGTCCTCTAATTCCAGGATCAACAACAACCAGTACGGGACCATGCAAAACCTGCACGACTCCTGCTCAAGGCAACTCTATGTTTCCCTCATGTTGCTGTACAAAACCTACGGATGGAAATTGCACCTGTATTCCCATCCCATCGTCCTGGGCTTTCGCAAAATACGCATGGGAGTGGGCCTCAGTCCGTTTCTCTTGGCTCAGTTTACTAGTGCCATTTGTTCAGTGGTTCGTAGGG |
| 163W | 5'-ACAAGAATCCTCACAATACCGCAGAGTCTAGACTCGTGGTGGACTTCTCTCAATTTTCTAGGGGGATCTCCCGTGTGTCTTGGCCAAAATTCGCAGTCCCCAACCTCCAATCACTCACCAACCTCCTGTCCTCCAATTTGTCCTGGTTATCGCTGGATGTGTCTGCGGCGTTTTATCATATTCCTCTTCATCCTGCTGCTATGCCTCATCTTCTTATTGGTTCTTCTGGATTATCAAGGTATGTTGCCCGTTTGTCCTCTAATTCCAGGATCAACAACAACCAGTACGGGACCATGCAAAACCTGCACGACTCCTGCTCAAGGCAACTCTATGTTTCCCTCATGTTGCTGTACAAAACCTACGGATGGAAATTGCACCTGTATTCCCATCCCATCGTCCTGGGCTTTCGCAAAATACCTAGCAGAGTGGGCCTCAGTCCGTTTCTCTTGGCTCAGTTTACTAGTGCCATTTGTTCAGTGGTTCGTAGGG |
| 164E | 5'-ACAAGAATCCTCACAATACCGCAGAGTCTAGACTCGTGGTGGACTTCTCTCAATTTTCTAGGGGGATCTCCCGTGTGTCTTGGCCAAAATTCGCAGTCCCCAACCTCCAATCACTCACCAACCTCCTGTCCTCCAATTTGTCCTGGTTATCGCTGGATGTGTCTGCGGCGTTTTATCATATTCCTCTTCATCCTGCTGCTATGCCTCATCTTCTTATTGGTTCTTCTGGATTATCAAGGTATGTTGCCCGTTTGTCCTCTAATTCCAGGATCAACAACAACCAGTACGGGACCATGCAAAACCTGCACGACTCCTGCTCAAGGCAACTCTATGTTTCCCTCATGTTGCTGTACAAAACCTACGGATGGAAATTGCACCTGTATTCCCATCCCATCGTCCTGGGCTTTCGCAAAATACCTATGGGCATGGGCCTCAGTCCGTTTCTCTTGGCTCAGTTTACTAGTGCCATTTGTTCAGTGGTTCGTAGGG |
| 165W | 5'-ACAAGAATCCTCACAATACCGCAGAGTCTAGACTCGTGGTGGACTTCTCTCAATTTTCTAGGGGGATCTCCCGTGTGTCTTGGCCAAAATTCGCAGTCCCCAACCTCCAATCACTCACCAACCTCCTGTCCTCCAATTTGTCCTGGTTATCGCTGGATGTGTCTGCGGCGTTTTATCATATTCCTCTTCATCCTGCTGCTATGCCTCATCTTCTTATTGGTTCTTCTGGATTATCAAGGTATGTTGCCCGTTTGTCCTCTAATTCCAGGATCAACAACAACCAGTACGGGACCATGCAAAACCTGCACGACTCCTGCTCAAGGCAACTCTATGTTTCCCTCATGTTGCTGTACAAAACCTACGGATGGAAATTGCACCTGTATTCCCATCCCATCGTCCTGGGCTTTCGCAAAATACCTATGGGAGGCAGCCTCAGTCCGTTTCTCTTGGCTCAGTTTACTAGTGCCATTTGTTCAGTGGTTCGTAGGG |
| 167S | 5'-ACAAGAATCCTCACAATACCGCAGAGTCTAGACTCGTGGTGGACTTCTCTCAATTTTCTAGGGGGATCTCCCGTGTGTCTTGGCCAAAATTCGCAGTCCCCAACCTCCAATCACTCACCAACCTCCTGTCCTCCAATTTGTCCTGGTTATCGCTGGATGTGTCTGCGGCGTTTTATCATATTCCTCTTCATCCTGCTGCTATGCCTCATCTTCTTATTGGTTCTTCTGGATTATCAAGGTATGTTGCCCGTTTGTCCTCTAATTCCAGGATCAACAACAACCAGTACGGGACCATGCAAAACCTGCACGACTCCTGCTCAAGGCAACTCTATGTTTCCCTCATGTTGCTGTACAAAACCTACGGATGGAAATTGCACCTGTATTCCCATCCCATCGTCCTGGGCTTTCGCAAAATACCTATGGGAGTGGGCCGCAGTCCGTTTCTCTTGGCTCAGTTTACTAGTGCCATTTGTTCAGTGGTTCGTAGGG |
| 168V | 5'-ACAAGAATCCTCACAATACCGCAGAGTCTAGACTCGTGGTGGACTTCTCTCAATTTTCTAGGGGGATCTCCCGTGTGTCTTGGCCAAAATTCGCAGTCCCCAACCTCCAATCACTCACCAACCTCCTGTCCTCCAATTTGTCCTGGTTATCGCTGGATGTGTCTGCGGCGTTTTATCATATTCCTCTTCATCCTGCTGCTATGCCTCATCTTCTTATTGGTTCTTCTGGATTATCAAGGTATGTTGCCCGTTTGTCCTCTAATTCCAGGATCAACAACAACCAGTACGGGACCATGCAAAACCTGCACGACTCCTGCTCAAGGCAACTCTATGTTTCCCTCATGTTGCTGTACAAAACCTACGGATGGAAATTGCACCTGTATTCCCATCCCATCGTCCTGGGCTTTCGCAAAATACCTATGGGAGTGGGCCTCAGCACGTTTCTCTTGGCTCAGTTTACTAGTGCCATTTGTTCAGTGGTTCGTAGGG |
| 169R | 5'-ACAAGAATCCTCACAATACCGCAGAGTCTAGACTCGTGGTGGACTTCTCTCAATTTTCTAGGGGGATCTCCCGTGTGTCTTGGCCAAAATTCGCAGTCCCCAACCTCCAATCACTCACCAACCTCCTGTCCTCCAATTTGTCCTGGTTATCGCTGGATGTGTCTGCGGCGTTTTATCATATTCCTCTTCATCCTGCTGCTATGCCTCATCTTCTTATTGGTTCTTCTGGATTATCAAGGTATGTTGCCCGTTTGTCCTCTAATTCCAGGATCAACAACAACCAGTACGGGACCATGCAAAACCTGCACGACTCCTGCTCAAGGCAACTCTATGTTTCCCTCATGTTGCTGTACAAAACCTACGGATGGAAATTGCACCTGTATTCCCATCCCATCGTCCTGGGCTTTCGCAAAATACCTATGGGAGTGGGCCTCAGTCGCATTCTCTTGGCTCAGTTTACTAGTGCCATTTGTTCAGTGGTTCGTAGGG |
| 170F | 5'-ACAAGAATCCTCACAATACCGCAGAGTCTAGACTCGTGGTGGACTTCTCTCAATTTTCTAGGGGGATCTCCCGTGTGTCTTGGCCAAAATTCGCAGTCCCCAACCTCCAATCACTCACCAACCTCCTGTCCTCCAATTTGTCCTGGTTATCGCTGGATGTGTCTGCGGCGTTTTATCATATTCCTCTTCATCCTGCTGCTATGCCTCATCTTCTTATTGGTTCTTCTGGATTATCAAGGTATGTTGCCCGTTTGTCCTCTAATTCCAGGATCAACAACAACCAGTACGGGACCATGCAAAACCTGCACGACTCCTGCTCAAGGCAACTCTATGTTTCCCTCATGTTGCTGTACAAAACCTACGGATGGAAATTGCACCTGTATTCCCATCCCATCGTCCTGGGCTTTCGCAAAATACCTATGGGAGTGGGCCTCAGTCCGTGCATCTTGGCTCAGTTTACTAGTGCCATTTGTTCAGTGGTTCGTAGGG |
| 171S | 5'-ACAAGAATCCTCACAATACCGCAGAGTCTAGACTCGTGGTGGACTTCTCTCAATTTTCTAGGGGGATCTCCCGTGTGTCTTGGCCAAAATTCGCAGTCCCCAACCTCCAATCACTCACCAACCTCCTGTCCTCCAATTTGTCCTGGTTATCGCTGGATGTGTCTGCGGCGTTTTATCATATTCCTCTTCATCCTGCTGCTATGCCTCATCTTCTTATTGGTTCTTCTGGATTATCAAGGTATGTTGCCCGTTTGTCCTCTAATTCCAGGATCAACAACAACCAGTACGGGACCATGCAAAACCTGCACGACTCCTGCTCAAGGCAACTCTATGTTTCCCTCATGTTGCTGTACAAAACCTACGGATGGAAATTGCACCTGTATTCCCATCCCATCGTCCTGGGCTTTCGCAAAATACCTATGGGAGTGGGCCTCAGTCCGTTTCGCATGGCTCAGTTTACTAGTGCCATTTGTTCAGTGGTTCGTAGGG |
| 172W | 5'-ACAAGAATCCTCACAATACCGCAGAGTCTAGACTCGTGGTGGACTTCTCTCAATTTTCTAGGGGGATCTCCCGTGTGTCTTGGCCAAAATTCGCAGTCCCCAACCTCCAATCACTCACCAACCTCCTGTCCTCCAATTTGTCCTGGTTATCGCTGGATGTGTCTGCGGCGTTTTATCATATTCCTCTTCATCCTGCTGCTATGCCTCATCTTCTTATTGGTTCTTCTGGATTATCAAGGTATGTTGCCCGTTTGTCCTCTAATTCCAGGATCAACAACAACCAGTACGGGACCATGCAAAACCTGCACGACTCCTGCTCAAGGCAACTCTATGTTTCCCTCATGTTGCTGTACAAAACCTACGGATGGAAATTGCACCTGTATTCCCATCCCATCGTCCTGGGCTTTCGCAAAATACCTATGGGAGTGGGCCTCAGTCCGTTTCTCTGCACTCAGTTTACTAGTGCCATTTGTTCAGTGGTTCGTAGGG |
| gBlock Synthesis for Germline Antibodies | |
| H006-HC_GL | 5'-CTAGTAGCAACTGCAACCGGTGTACATTCTCAGGTGCAGCTGGTGGAGTCTGGGGGAGGCGTGGTCCAGCCTGGGAGGTCCCTGAGACTCTCCTGTGCAGCCTCTGGATTCACCTTCAGTAGCTATGGCATGCACTGGGTCCGCCAGGCTCCAGGCAAGGGGCTGGAGTGGGTGGCAGTTATATCATATGATGGAAGTAATAAATACTATGCAGACTCCGTGAAGGGCCGATTCACCATCTCCAGAGACAATTCCAAGAACACGCTGTATCTGCAAATGAACAGCCTGAGAGCTGAGGACACGGCTGTGTATTACTGTGCGAAAGATGCTTATCTTTCTGCAGCGAGAGGATACGGTATGGACGTCTGGGGCCAAGGGACCACGGTCACCGTCTCCTCAGCGTCGACCAAGGGCCCATCGG |
| H006-Lambda_GL | 5'-CTAGTAGCAACTGCAACCGGTTCCTGGGCCTCCTATGTGCTGACTCAGCCACCCTCGGTGTCAGTGGCCCCAGGACAGACGGCCAGGATTACCTGTGGGGGAAACAACATTGGAAGTAAAAGTGTGCACTGGTACCAGCAGAAGCCAGGCCAGGCCCCTGTGCTGGTCGTCTATGATGATAGCGACCGGCCCTCAGGGATCCCTGAGCGATTCTCTGGCTCCAACTCTGGGAACACGGCCACCCTGACCATCAGCAGGGTCGAAGCCGGGGATGAGGCCGACTATTACTGTCAGGTGTGGGATAGTAGTAGTGATCATGTGGTATTCGGCGGAGGGACCAAGCTGACCGTCCTAGGTCAGCCCAAGGCTGCCCCCTCGGTCACTCTGTTCCCACCCTCGAGTGAGGAGCTTCAAGCC |
| H019-HC_GL | 5'-CTAGTAGCAACTGCAACCGGTGTACATTCTCAGGTGCAGCTGCAGGAGTCGGGCCCAGGACTGGTGAAGCCTTCACAGACCCTGTCCCTCACCTGCACTGTCTCTGGTGGCTCCATCAGCAGTGGTGATTACTACTGGAGTTGGATCCGCCAGCCCCCAGGGAAGGGCCTGGAGTGGATTGGGTACATCTATTACAGTGGGAGCACCTACTACAACCCGTCCCTCAAGAGTCGAGTTACCATATCAGTAGACACGTCCAAGAACCAGTTCTCCCTGAAGCTGAGCTCTGTGACTGCCGCAGACACGGCCGTGTATTACTGTGCCATCTACATGGATGAGGCCTGGGCTTTTGATATCTGGGGCCAAGGGACAATGGTCACCGTCTCTTCAGCGTCGACCAAGGGCCCATCGG |
| H019-Kappa_GL | 5'-CTAGTAGCAACTGCAACCGGTGTACATTCTGACATCCAGATGACCCAGTCTCCATCCTCCCTGTCTGCATCTGTAGGAGACAGAGTCACCATCACTTGCCGGGCAAGTCAGAGCATTAGCAGCTATTTAAATTGGTATCAGCAGAAACCAGGGAAAGCCCCTAAGCTCCTGATCTATGCTGCATCCAGTTTGCAAAGTGGGGTCCCATCAAGGTTCAGTGGCAGTGGATCTGGGACAGATTTCACTCTCACCATCAGCAGTCTGCAACCTGAAGATTTTGCAACTTACTACTGTCAACAGAGTTACAGTATTTCCTTATTCACTTTTGGCCAGGGGACCAAGCTGGAGATCAAACGTACGGTGGCTGCACCATCTGTCTTC |
| H020-HC_GL | 5'-CTAGTAGCAACTGCAACCGGTGTACATTCTCAGGTGCAGCTGCAGGAGTCGGGCCCAGGACTGGTGAAGCCTTCGGAGACCCTGTCCCTCACCTGCACTGTCTCTGGTGGCTCCATCAGTAGTTACTACTGGAGCTGGATCCGGCAGCCCCCAGGGAAGGGACTGGAGTGGATTGGGTATATCTATTACAGTGGGAGCACCAACTACAACCCCTCCCTCAAGAGTCGAGTCACCATATCAGTAGACACGTCCAAGAACCAGTTCTCCCTGAAGCTGAGCTCTGTGACCGCCGCAGACACGGCCGTGTATTACTGTGCGAGACACCTTTATCGCTATGGTTATAGGAACTACTTTGACTACTGGGGCCAGGGAACCCTGGTCACCGTCTCCTCAGCGTCGACCAAGGGCCCATCGG |
| H020-Kappa_GL | 5'-CTAGTAGCAACTGCAACCGGTGTACATTCTGACATCCAGATGACCCAGTCTCCATCTTCCGTGTCTGCATCTGTAGGAGACAGAGTCACCATCACTTGTCGGGCGAGTCAGGGTATTAGCAGCTGGTTAGCCTGGTATCAGCAGAAACCAGGGAAAGCCCCTAAGCTCCTGATCTATGCTGCATCCAGTTTGCAAAGTGGGGTCCCATCAAGGTTCAGCGGCAGTGGATCTGGGACAGATTTCACTCTCACCATCAGCAGCCTGCAGCCTGAAGATTTTGCAACTTACTATTGTCAACAGGCTAACAGTTTCCCGCTCACTTTCGGCGGAGGGACCAAGGTGGAGATCAAACGTACGGTGGCTGCACCATCTGTCTTC |
| HBV TaqMan PCR | |
| F-sense | 5'-CCGTCTGTGCCTTCTCATCTG |
| R-anti-sense | 5'-AGTCCAAGAGTCCTCTTATGTAAGACCTT |
| Probe | 5-/56 FAM/ CCGTGTGCA/ZEN/CTTCGCTTC ACCTCT GC/3IABkFQ/-3 |
| S-protein PCR | |
| F-S-protein | 5'-CCCTGCGCTGAACATGGAGAACA |
| R-S-protein | 5'-AAATGTATACCCAAAGACAAAAGAAAA |
