## Supplemental Figures and Methods for "Public broadly neutralizing antibodies against hepatitis B virus in individuals with elite serologic activity"

**SUPPLEMENTARY FIGURES**

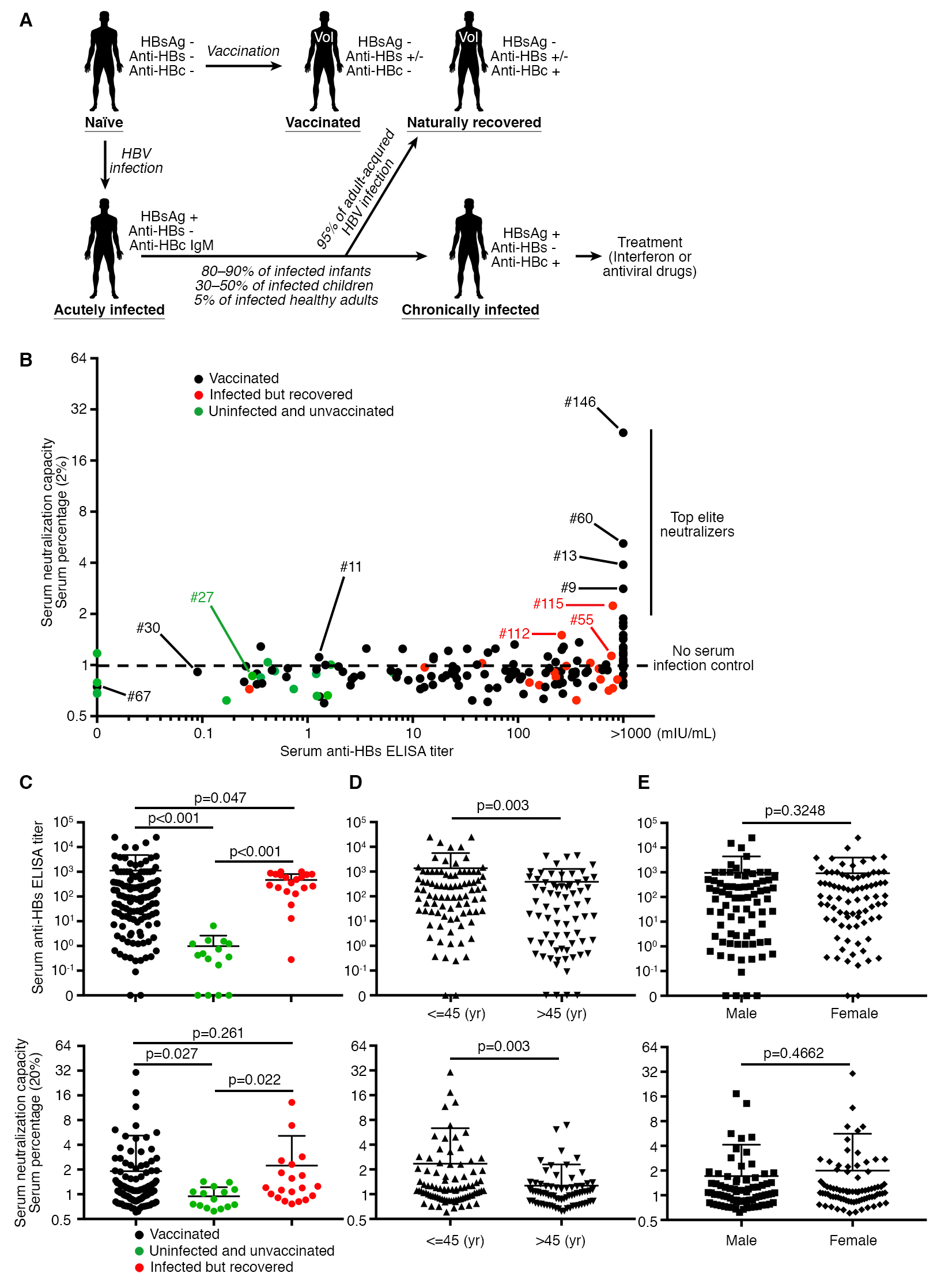

Figure S1. Characterization of Antibody Immune Response Against HBV, Related to Figure 1

(A) Schematic representation of different stages of HBV infection. Vaccinated or infected naturally recovered individuals were recruited for this study. (B) Sera (1:50 dilution in the final assay volume) from 159 volunteers were screened, see also Figure 1A. (C-E) Comparison of anti-HBs ELISA titers (upper panel) and their serum neutralization capacity (lower panel) between different groups of individuals. Vaccinated or recovered individuals show statistically higher anti-HBs titers (upper panel, C) and more potent neutralizing activity (lower panel, C) than the uninfected unvaccinated individuals. Younger individuals (≤45 years old) showed slightly higher antibody immune response against HBsAg (D). No difference was found between genders (E).

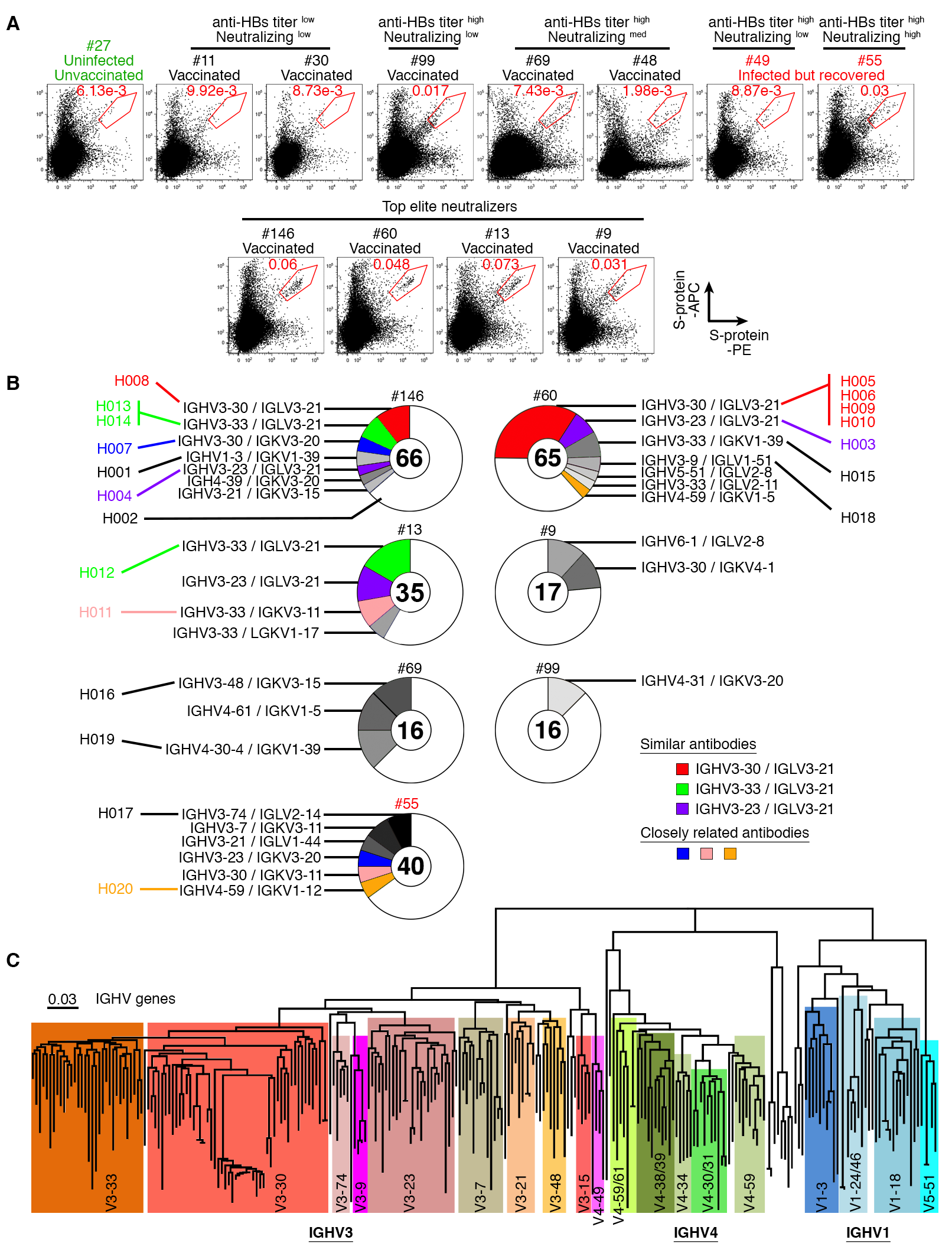

Figure S2. Antibody Cloning and Sequence Analysis of Anti-HBs, Related to Figure 2

(A) Frequency of S-protein-specific memory B cells in peripheral blood mononuclear cells of all twelve donors. Details are similar to Figure 2A. (B) Pie charts show the distribution of anti-HBs antibodies. Figure legends are similar to Figure 2C. VH and VL genes for each slice are shown and the 20 chosen anti-HBs antibodies are labeled. (C) Phylogenetic tree of all cloned anti-HBs antibodies based on IGH Fab region. IGH Fab regions of 244 antibodies were aligned followed by tree construction.

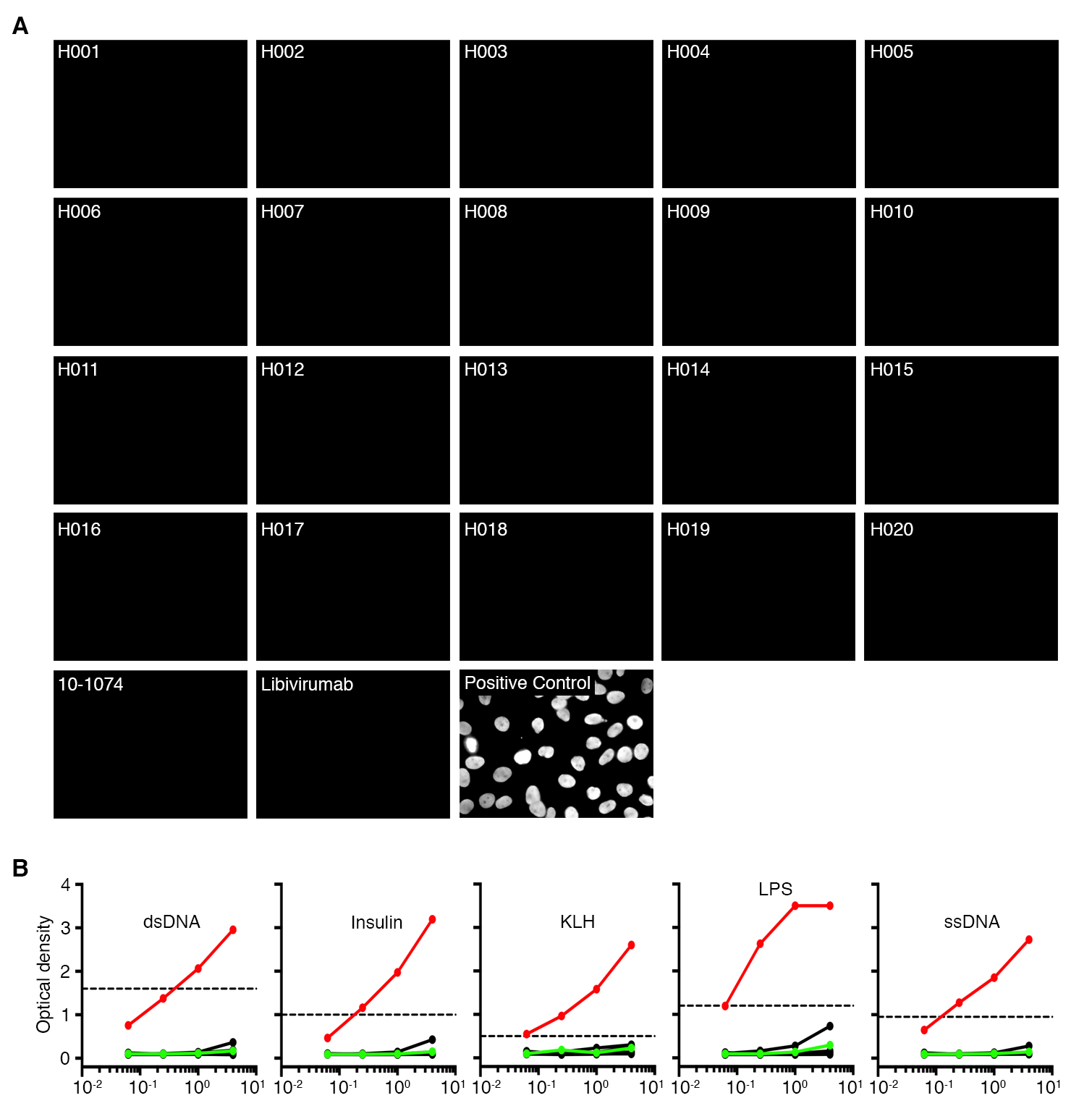

Figure S3. Autoreactivity of 20 anti-HBs antibodies, Related to Figure 3

(A) Autoreactivity of monoclonal antibodies. Positive control antibody efficiently stained the nucleus of HEp-2 cells. Twenty anti-HBs antibodies, as well as anti-HBs antibody libivirumab and anti-HIV antibody 10-1074, were also tested. (B) Polyreactivity profiles of 20 anti-HBs antibodies. ELISA measures antibody binding to the following antigens: double-stranded DNA (dsDNA), insulin, keyhole limpet hemocyanin (KLH), lipopolysaccharides (LPS), and single-stranded DNA (ssDNA). Red and green lines represent positive control antibody ED38 and negative control antibody mGO53 respectively, while dashed lines show cut-off values for positive reactivity (Gitlin et al., 2016).

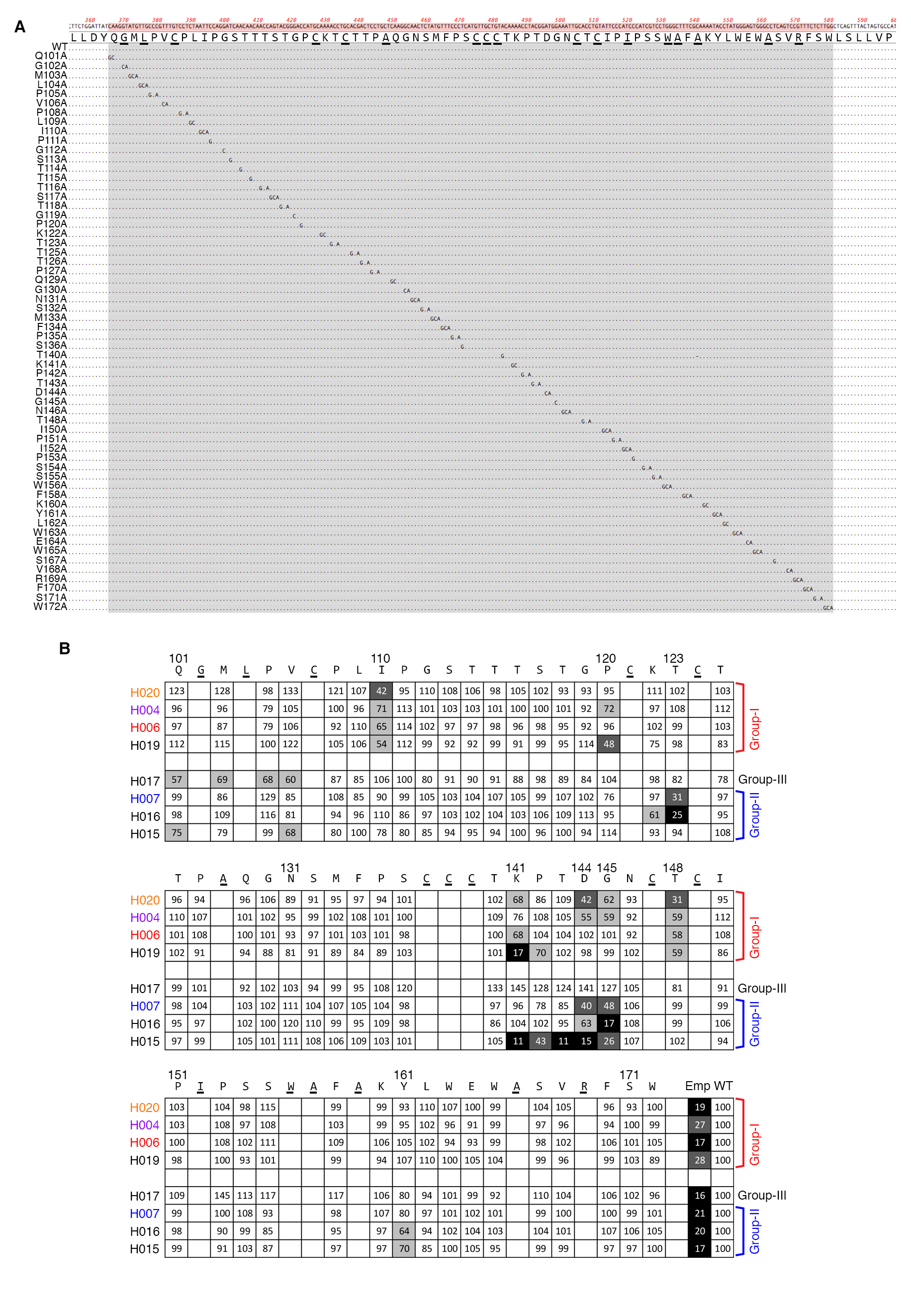

Figure S4. Alanine Scanning and Peptide Screening, Related to Figure 4

(A) Alignment of nucleotide sequences shows each alanine mutation in the antigenic loops. (B) Results of ELISA on alanine scanning mutants of HBsAg. Binding to mutants was normalized to wild-type S-protein: black, 0-25%; dark grey, 26-50%; light grey, 51-75%; white, >76%. Experiments were performed three times. Underlined cysteines, alanines, and amino acids known to be critical for S-protein production were not mutated (Salisse and Sureau, 2009).

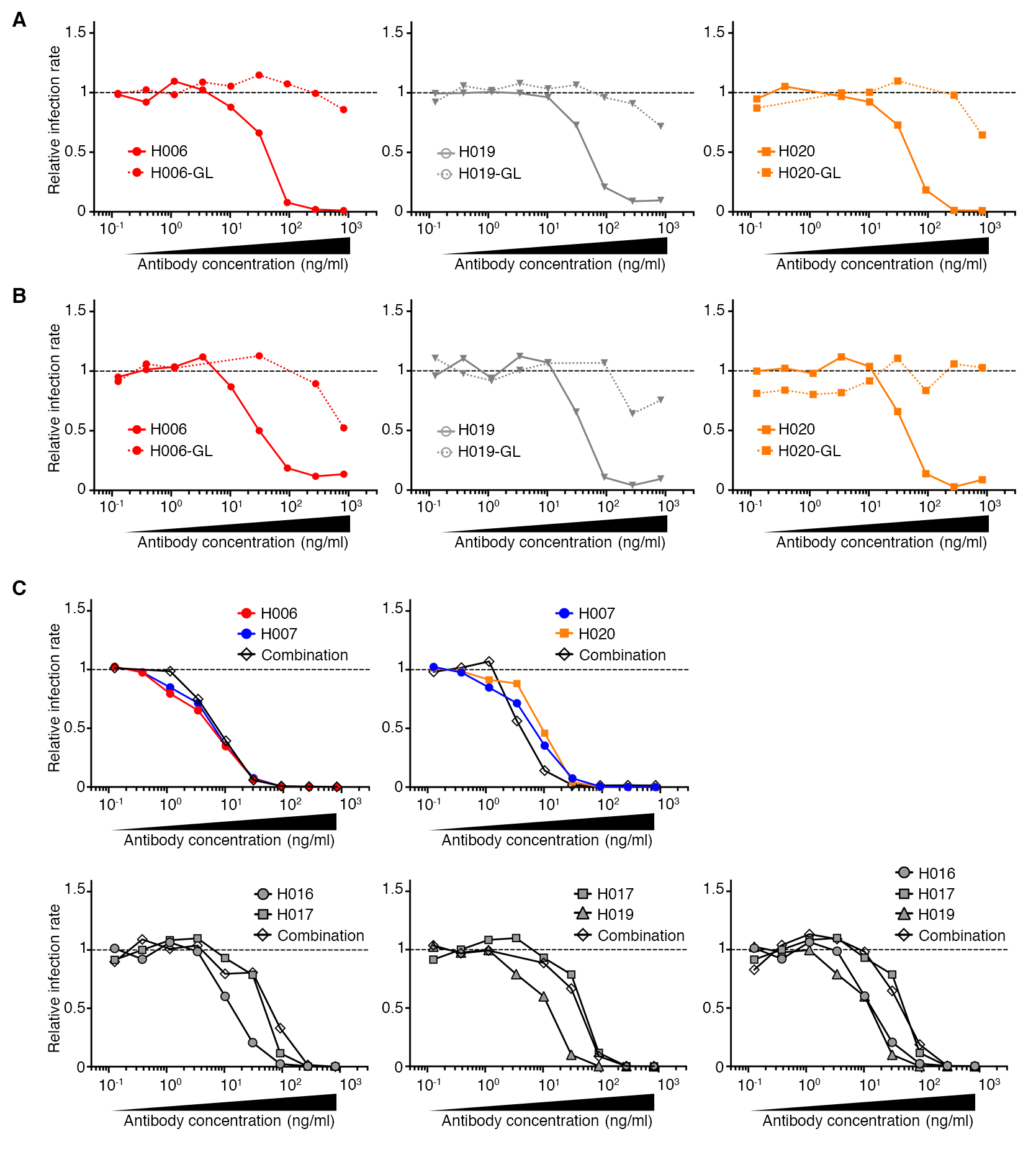

Figure S5. *In Vitro* Neutralization Assay of anti-HBs bNAb Germline Antibodies or Combinations, Related to Figure 5

(A-B) *In vitro* neutralization assay of anti-HBs bNAbs and their corresponding germline antibodies. The relative infection rates were calculated based on either HBsAg protein level in culture medium (A) or HBcAg staining intracellularly (B). (C) *In vitro* neutralization assay of anti-HBs bNAbs recognizing different epitopes and the same total amount of antibody combination at 1:1 or 1:1:1 ratio.

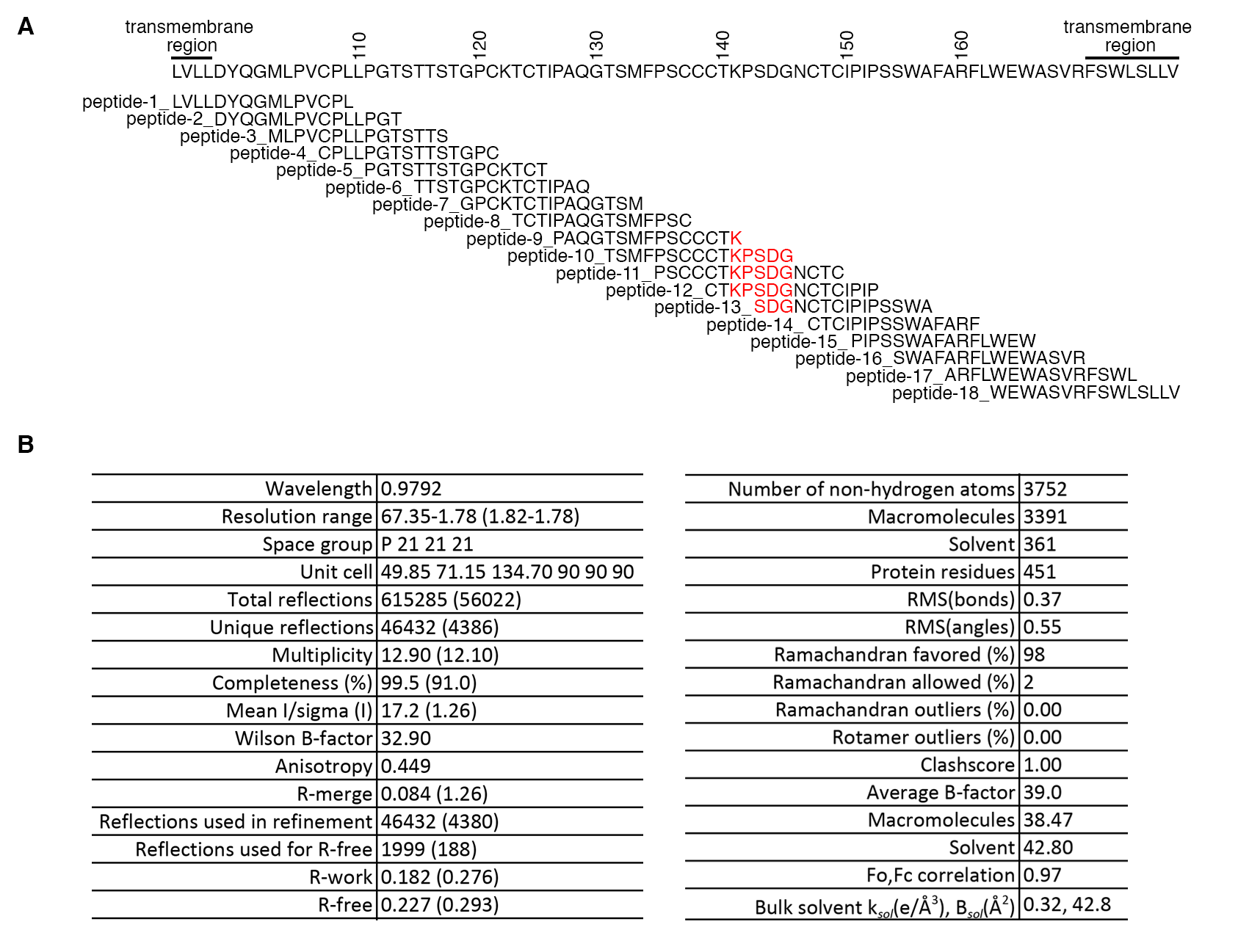

Figure S6. Detailed Information of Crystal Structure of H015 and Its Linear Epitope, Related to Figure 6

(A) Synthesized peptides for antigenic loop region were subjected to ELISA for antibody binding. Among the tested antibodies, only H015 binds peptide-11 and -12. (B) Data collection and refinement statistics for H015 Fab are summarized. Statistics for the highest-resolution shell are shown in parentheses. Refinement program PHENIX 1.16.

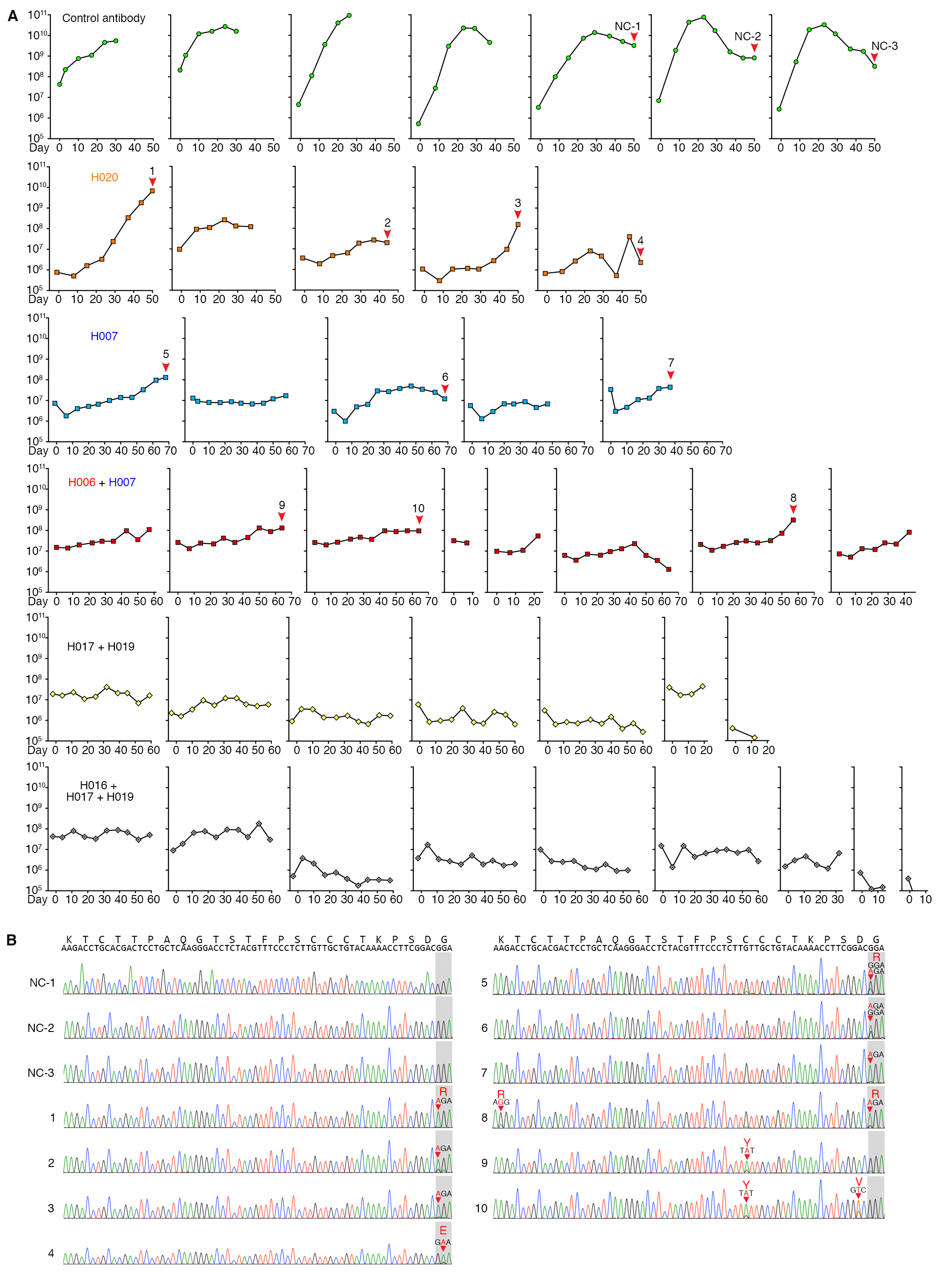

Figure S7. HBV DNA levels and S-protein Sequences in Antibody-Treated huFNRG Mice, Related to Figure 7

(A) HBV DNA levels in representative individual huFNRG mice treated by control antibody 10-1074, anti-HBs bNAb H020, anti-HBs bNAb H007, combination of anti-HBs bNAb (H006 + H007), (H017 + H019), and (H016 + H017 + H019). HBV DNA levels in mouse sera were monitored on a weekly basis. The mice without red arrows bear no escape mutations at the last time point.

(B) Part of the S-protein sequences from the indicated mice (red arrows and numbers) are shown below as chromatograms, with mutations marked by red arrowheads.

**KEY RESOURCES TABLE**

| REAGENT or RESOURCE | SOURCE | IDENTIFIER |
| --- | --- | --- |
| **Experimental Models: Cell Lines** |  |  |
| Human Hepatocytes, Cryopreserved, Plateable and Interaction Qualified | Lonza Bioscience | Cat#HUCPI |
| Hepatocyte Defined Medium | Corning | Cat#05449 |
| HepG2-NTCP | (Michailidis et al., 2017) | N/A |
| HEK293-6E | National Research Council of Canada | NRC file 11565 |
| HepDE19 cells | (Cai et al., 2012) | N/A |
| Huh-7.5 | (Robbiani et al., 2017) | N/A |
| **Exprimental Models: Mouse Strains** |  |  |
| *Fah*^-/-^NOD*Rag*1^-/-^*IL2rg*^-/-^ mouse  (huFNRG) | (de Jong et al., 2014) | N/A |
| **Bacteria and Viruses** |  |  |
| Subcloning Efficiency™ DH5α™ Competent Cells | Thermo Fisher Scientific | Cat#18265017 |
| HBV viruses | (Cai et al., 2012) | N |
| **Antibodies** |  |  |
| Human recombinant 10-1074 | (Mouquet et al., 2012) | N/A |
| Human recombinant ED38 | (Wardemann et al., 2003) | N/A |
| Human recombinant mG053 | (Yurasov et al., 2005) | N/A |
| Goat anti-Human IgG (H+L) Secondary Antibody, HRP | Thermo Fisher Scientific | Cat#31410 |
| Alexa Fluor 488 Mouse anti-Human CD19 | BD Biosciences | Cat#557697 |
| BV421 Mouse Anti-Human CD19 | BD Biosciences | Cat#562440 |
| anti-CD20-PECy7 | BD Biosciences | Cat#335811 |
| Anti-CD27-PE | BD Biosciences | Cat#555441 |
| APC Mouse Anti-Human IgG | BD Pharmingen | Cat#550931 |
| Bv421 Mouse Anti-Human IgG | BD Biosciences | Cat#562581 |
| Anti-Hepatitis B virus core antigen IgG | AUSTRAL Biologicals | Cat#HBP-023-9 |
| Alexa Fluor® 488 AffiniPure Goat Anti-Human IgG, F(ab')2 fragment specific | Jackson ImmunoResearch | Cat#109-545-006 |
| Goat anti-Rabbit IgG (H+L) Alexa Fluor 594 | Thermo Fisher Scientific | Cat#A11037 |
| **Chemicals and Proteins** |  |  |
| Streptavidin HRP | BD Biosciences | Cat#554066 |
| APC Streptavidin | BD Biosciences | Cat#554067 |
| Strep-PE | eBioscience | Cat#12-4317-87 |
| Streptavidin, Alexa Fluor™ 488 | Thermo Fisher Scientific | Cat#S32354 |
| Human BD Fc Block™ | BD Biosciences | Cat#564220 |
| Ovalbumin (257-264) chicken | Sigma-Aldrich | Cat#S7951 |
| HBsAg adr CHO | ProSpec | Cat#HBS-875 |
| HBsAg adw | ProSpec | Cat#HBS-872 |
| HBsAg protein adr | Fitzgerald | Cat#30-AH37 |
| HBsAg protein ay | Fitzgerald | Cat#30-1816 |
| HBsAg protein ad | Fitzgerald | Cat#30-AH15 |
| HBsAg protein ayw | Fitzgerald | Cat#30R-AH018 |
| RNAsin Plus RNAse inhibitor | Promega | Cat#N2615 |
| Random Primers | Thermo Fisher Scientific | Cat#48190011 |
| Lipopolysaccharides from *E. coli* O55:B5 | Sigma-Aldrich | Cat#L2637 |
| Insulin solution human | Sigma-Aldrich | Cat#I1927 |
| Deoxyribonucleic acid from calf thymus | Sigma-Aldrich | Cat#4522 |
| Hemocyanin from Megathura crenulata (keyhole limpet) | Sigma-Aldrich | Cat#H8283 |
| Poly(ethylene glycol) | Sigma-Aldrich | Cat#81268 |
| Normal Goat Serum | Jackson ImmunoResearch | Cat#005-000-121 |
| DAPI, FluoroPure™ grade | Thermo Fisher Scientific | Cat#D21490 |
| Paraformaldehyde 4% Aqueous Solution | Electron Microscopy Sciences | Cat#157-4 |
| Phusion High-Fidelity DNA Polymerase | Thermo Fisher Scientific | Cat#F-530L |
| **Critical Commercial Assays** |  |  |
| ARCHITECT Anti-HBs | Abbott Laboratories | Cat#B7C180 |
| ARCHITECT HBsAg Qualitative | Abbott Laboratories | Cat#B1P970 |
| ARCHITECT Anti-HBc II | Abbott Laboratories | Cat#B8L440 |
| HBsAg CLIA kit | Autobio Diagnostics Co. | Cat#CL0310-2 |
| HBeAg CLIA kit | Autobio Diagnostics Co. | Cat#CL0312-2 |
| Anti-HBs CLIA kit | Autobio Diagnostics Co. | Cat#CL0311-2 |
| LS magnetic columns | Miltenyi Biotech | Cat#130-042-401 |
| CD19 MicroBeads, human | Miltenyi Biotech | Cat#130-097-055 |
| EZ-Link™ Micro NHS-PEG4-Biotinylation Kit | Thermo Fisher Scientific | Cat#21955 |
| Superscript III Reverse Transcriptase | Thermo Fisher Scientific | Cat#18080044 |
| QIAamp DNA blood mini kit | Qiagen | Cat#51104 |
| TaqMan Universal PCR Master Mix | Applied Biosystems | Cat#4304437 |
| Antinuclear antibodies (HEp-2) Kit | MBL International | Cat#ANK-120 |
| Centricon Plus-70 Ultracel PL-100 | Millipore Sigma | Cat#UFC710008 |
| X-tremeGENE 9 DNA Transfection Reagent | Sigma-Aldrich | Cat#6365787001 |
| **Plasmids** |  |  |
| IGγ1 expression vector | (von Boehmer et al., 2016) | N/A |
| IGκ expression vector | (von Boehmer et al., 2016) | N/A |
| IGλ expression vector | (von Boehmer et al., 2016) | N/A |
| p1.3xHBV-WT | Laboratory of Charles M. Rice | N/A |
| **Softwares and Websites** |  |  |
| PRISM | GraphPad | https://www.graphpad.com |
| IgBlast | (Ye et al., 2013) | http://www.ncbi.nlm.nih.gov/igblast/ |
| IMGT/V-QUEST | (Lefranc et al., 2015) | http://www.imgt.org/IMGT_vquest/vquest |
| Geneious Prime | Geneious | https://www.geneious.com/ |

**CONTACT FOR REAGENT AND RESOURSE SHARING**

Further information and requests for reagents should be directed to and will be fulfilled by the contacts, Qiao Wang or Ype P. de Jong. Sharing of antibodies and other reagents with academic researchers may require UBMTA agreements.

**EXPERIMENTAL MODELS AND SUBJECTS**

**Human Subjects**

Volunteer recruitment and blood draws were performed at the Rockefeller University Hospital under a protocol approved by the institutional review board (IRB QWA-0947). Study participants ranged in age from 22-65 with a mean of 43, the female:male ratio was 81:78 (Figure S1D and S1E; Table S1). Additional information on the sex and age of study participants can be obtained upon request.

**Mice**

*Fah*^-/-^NOD*Rag*1^-/-^*IL2rg*^-/-^ (FNRG) mice were produced as reported (de Jong et al., 2014) and maintained in the AAALAC-certified facility of the Rockefeller University. Animal protocols were in accordance with NIH guidelines and approved by the Rockefeller University Institutional Animal Care and Use Committee under protocol #18063.

**Cell Lines**

HepG2-NTCP cells (Michailidis et al., 2017) and HepDE19 cells (Cai et al., 2012) were maintained in collagen-coated flasks in Dulbecco's Modified Eagle Medium (DMEM) supplemented with 10% or 3% fetal bovine serum (FBS) and 0.1 mM non-essential amino acids (NEAA). Huh7.5-NTCP cells were maintained in DMEM supplemented with 10% FBS and 0.1 mM NEAA. All liver cell lines were cultured at 37°C in 5% CO_2_. Human embryonic kidney HEK293-6E suspension cells were cultured at 37°C in 8% CO_2_ with shaking at 120 rpm.

**Viruses**

HBV-containing supernatant from HepDE19 cells was collected and concentrated as previously described (Michailidis et al., 2017). The concentrated virus stock was aliquoted and stored at -80°C.

**Bacteria**

*E. coli* DH5-alpha were cultured at 37°C with shaking at 230 rpm.

**METHODS**

**Collection of Human Samples**

Samples of peripheral blood were collected from volunteers at the Rockefeller University Hospital. Serum was isolated by centrifugation of coagulated whole blood, and aliquoted for storage at -80°C. PBMCs were isolated using a cell separation tube with frit barrier and cryopreserved in liquid nitrogen in 90% heat-inactivated FBS supplemented with 10% dimethylsulfoxide (DMSO).

**HBV Stock**

HepDE19 cells (Cai et al., 2012) were cultured in the absence of tetracycline to induce HBV replication. After seven days, supernatant was collected every other day for two weeks and fresh medium was added. After each collection, medium was spun down to remove cell debris, passed through a 0.22 μm filter, and kept at 4°C. Collected medium was concentrated 100-fold via centrifugation using Centricon Plus-70 centrifugal filter devices (Millipore-Sigma, Billerica, MA).

***In Vitro* HBV Neutralization Assay**

*In vitro* HBV infection was performed as previously described (Michailidis et al., 2017). Briefly, HepG2-NTCP cells were seeded in 96-well collagen-coated plates in DMEM supplemented with 10% FBS and 0.1 mM NEAA. The medium was changed to DMEM with 3% FBS, 0.1 mM NEAA, and 2% DMSO the next day and cultured for an additional 24 hours before infection. The inoculation was in DMEM supplemented with 3% FBS and 0.1 mM NEAA 4% PEG and 2% DMSO. Antibodies or serum samples were incubated with the virus in the inoculation medium for one hour at 37°C before adding to cells. Serum neutralization capacity was calculated by the reverse of the percentage of infected HepG2-NTCP cells immunostained by rabbit anti-HBV core antibody (AUSTRAL Biologicals). For the blocking neutralization assay, S-protein antigen at different concentration was incubated with purified polyclonal antibodies for one hour at 37°C before incubation with HBV virus. The cells were then spinoculated for one hour by centrifugation at 1,000 g at 37°C. After a 24-hour incubation, supernatant was removed, cells were washed five times with PBS, and 100 μl of fresh DMEM supplemented with 3% FBS, 0.1 mM NEAA, and 2% DMSO. Both supernatant and cells were harvested 7 days after infection for analysis. Neutralization assays in primary human hepatocytes were performed as above using hepatocytes from livers of highly humanized mice that were harvested and seeded on collagen-coated plates in hepatocyte defined medium (Corning) (Michailidis et al., 2020).

**Chemiluminescence Immunoassay**

For quantitative analysis of secreted antigen HBsAg or HBeAg, 50 μl of the collected supernatant was loaded into 96-well plates of a chemiluminescence immunoassay (CLIA) kit (Autobio Diagnostics Co., Zhengzhou, China) according to the manufacturer’s instructions. Plates were read using a FLUOstar Omega luminometer (BMG Labtech). The absolute concentrations were measured and the relative values were calculated by normalizing to the saturation level of infection in the same lane.

**Immunofluorescence**

Cells were fixed in 4% paraformaldehyde for 20 minutes at room temperature, washed with PBS and permeabilized with 0.1% Triton X-100 in PBS. After blocking with 5% goat serum, the cells were incubated with rabbit anti-HBV core antibody (AUSTRAL Biologicals) overnight at 4°C and visualized with goat anti-rabbit Alexa Fluor 594 (Thermo Fisher Scientific). Nuclei were stained with DAPI. Cells were imaged using a Nikon Eclipse TE300 fluorescent microscope and processed with ImageJ. For high-content imaging analysis ImageXpress Micro XLS (Molecular Devices, Sunnyvale, CA) was used. The absolute HBc^+^ percentages were obtained and the relative infection rates were calculated by normalizing to the saturation level of infection in the same lane.

**ELISA Assays**

Blood samples were submitted to Memorial Sloan Kettering Cancer Center for clinical testing. The presence of HBsAg protein and anti-HBc antibody, as well as anti-HBs titers, were determined by ELISA (Abbott Laboratories) as per the manufacturer's instructions.

The binding of serum or recombinant IgG antibodies to HBsAg proteins (see KEY RESOURCES TABLE) was measured by coating ELISA plates with 10 μg/ml of antigen in PBS. Plates were blocked with 2% BSA in PBS and incubated with antibody for one hour at room temperature. Visualization was with HRP-conjugated goat anti-human IgG (Thermo Fisher Scientific). The 50% effective concentration (EC_50_) needed for maximal binding was determined by non-linear regression analysis in software PRISM.

For competition ELISAs plates were coated with 0.12 μg/ml HBsAg (adr CHO) and incubated with 16.7 μg/ml primary antibody for two hours, followed by directly adding 0.25 μg/ml biotinylated secondary antibody and incubation for 30 minutes all at room temperature. Detection was with streptavidin-HRP (BD Biosciences).

**Autoreactivity and Polyreactivity**

Autoreactivity and polyreactivity assays were performed as described (Gitlin et al., 2016; Mayer et al., 2017; Robbiani et al., 2017). For the autoreactivity assays, monoclonal antibodies were tested with the Antinuclear antibodies (HEp-2) Kit (MBL International). Antibodies were incubated at 100 μg/ml and were detected with Alexa Fluor 488 AffiniPure F(ab')₂ Fragment Goat Anti-Human IgG (H+L) (Jackson ImmunoResearch) at 10 μg/ml. Fluorescence images were taken with a wide-field fluorescence microscope (Axioplan 2, Zeiss), a 40x dry objective and a Hamamatsu Orca ER B/W digital camera. Images were analyzed with Image J. Human serum containing antinuclear antibodies (MBL International) was used as a positive control. For the polyreactivity ELISA assays, antibody binding to five different antigens, double-stranded DNA (dsDNA), insulin, keyhole limpet hemocyanin (KLH), lipopolysaccharides (LPS), and single-stranded DNA (ssDNA), were measured. ED38 (Wardemann et al., 2003) and mG053 (Yurasov et al., 2005) antibodies were used as positive and negative controls, respectively.

**Synthetic Peptides**

Eighteen peptides spanning the antigenic loop region of S-protein antigen were synthesized at the Proteomics Resource Center of The Rockefeller University. For peptide ELISAs plates were coated with 10 μg/ml peptide in PBS.

**HBsAg-Binding Memory B cells**

S-protein (adr serotype) expressed and purified from Chinese hamster ovary (CHO) cells (ProSpec) and ovalbumin (Sigma-Aldrich) were biotinylated using EZ-Link™ Micro NHS-PEG4-Biotinylation kit (Thermo Fisher Scientific). S-protein-PE and S-protein-APC were prepared by incubating 2-3 μg of biotin-S-protein with streptavidin-PE (eBioscience) or streptavidin-APC (BD Biosciences) in PBS respectively overnight at 4°C in the dark. Ovalbumin-Alexa Fluor 488 was generated by incubating biotin-ovalbumin with streptavidin-Alexa Fluor 488 (Thermo Fisher Scientific).

B cell purification, labeling, and sorting were as previously described (Escolano et al., 2019; Robbiani et al., 2017; Tiller et al., 2008; von Boehmer et al., 2016). Briefly, PBMCs were thawed and washed with RPMI medium at 37°C. B lymphocytes were positively selected using CD19 MicroBeads (Miltenyi Biotec) followed by incubation with human Fc block (BD Biosciences) and anti-CD20-PECy7 (BD Biosciences), anti-IgG-Bv421 (BD Biosciences), S-protein-PE at 10 μg/ml, S-protein-APC at 10 μg/ml, and ovalbumin-Alexa Fluor 488 at 10 μg/ml at 4°C for 20 minutes. Single CD20^+^ IgG^+^ S-protein-PE^+^ S-protein-APC^+^ Ova-Alexa Fluor 488^−^ memory B cells were sorted into 96-well plates using a FACSAriaII (Becton Dickinson) and stored at -80°C.

**Antibody Cloning, Sequencing and Production**

Antibody cloning, sequencing and production were done as previously reported (Robbiani et al., 2017; Tiller et al., 2008; von Boehmer et al., 2016). Primers are listed in Table S3. Germline antibody sequences of H006, H019 and H020 were synthesized by gBlock IDT (Table S3) and were inserted into antibody vectors for expression.

**S-protein Mutagenesis**

Oligonucleotides fragments with the target point mutations were synthesized by gBlock IDT (Table S3), and were substituted into the antigenic loop region in plasmid p1.3xHBV-WT by Sequence and Ligation-Independent Cloning (SLIC) (Jeong et al., 2012). Mutant plasmids were transfected into Huh-7.5-NTCP cells using X-tremeGENE 9 DNA Transfection Reagent (Sigma-Aldrich) and the culture medium was changed to serum-free DMEM after 24 hours. Supernatants were collected 2 days later and stored at -80°C. Serum-free medium (50 μl) was directly used to coat ELISA plates.

**Crystallization, X-ray Data Collection, Structure Determination and Refinement**

Antibody Fab (25 mg/ml) in 50 mM Tris 8.0, 50 mM NaCl was mixed with peptide (5 mg/ml) in the same buffer at 5:1 v/v. Crystals were obtained upon substitution of all peptide-11 cysteine residues with serine in the peptide synthesis (Proteomics Resource Center, RU). The crystallization condition for Fab15/peptide-11Ser was identified from a commercial screen (Morpheus by Molecular Dimensions) by the sitting-drop vapor-diffusion method at room temperature. The crystal used for data collection was obtained directly from the initial setup (position E1) in a precipitant solution consisting of 0.12 M Ethylene glycols (Di, Tri, Tetra and Penta-ethylene glycol), 0.1 M Buffer Mix 1 (Imidazole/MES) at pH 6.5 and 30% Precipitant Mix 1 (20% v/v PEG 500* MME; 10 % w/v PEG 20000). The crystals were flash-cooled in liquid nitrogen directly from the mother liquor without additional cryoprotectant. X-ray diffraction data were collected from a single crystal on the Advanced Photon Source (APS) beamline 24-ID-E to 1.78 Å resolution. The data were integrated and scaled with the program XDS (Kabsch, 2010a, b) and other data processing utilities from the CCP4 suite (Collaborative Computational Project, 1994) using RAPD, the software available at the beam-line. Initial phase estimates and electron-density maps were obtained by molecular replacement with Phaser (McCoy et al., 2007) using a single FAB molecule from (PDB: 5GGU) as an initial search model in Phenix (Adams et al., 2010). Iterative model building and structural refinement were manually performed using COOT (Emsley et al., 2010) and Phenix, respectively. The peptide density was well defined, and refined to 90% occupancy, for residues STKPSDGNST. All other residues were not visible and the area where they would be is fully solvent, with no crystal contacts involving any of the peptide atoms. The quality of the final model was good as noted in a Ramachandran of 96% of the observed residues within the allowable region. Data-collection and refinement statistics are summarized (Figure S6B). All molecular graphics were prepared with PyMOL (Version 2.0 Schrödinger, LLC). Atomic coordinates and experimental structure factors have been deposited in the PDB under accession code 6VJT.

**Humanized Mice and *In Vivo* Studies**

Six to eight week old FNRG female mice were transplanted with one million human hepatocytes from a pediatric female donor HUM4188 (Lonza Bioscience) as previously described (de Jong et al., 2014). Briefly, during isoflurane anesthesia mice underwent skin and peritoneal incision, exposing the spleen. One million hepatocytes were injected in the spleen using a 28-gauge needle. The peritoneum was then approximated using 4.0 VICRYL sutures (Johnson & Johnson), and skin was closed using MikRon Autoclip surgical clips (Becton Dickinson). Mice were cycled off the drug nitisinone (Yecuris) on the basis of weight loss and overall health. Humanization was monitored by human albumin quantification in mouse serum using a human-specific ELISA (Bethyl Labs). Humanized FNRG mice with human albumin values greater than 1 mg/ml were used for infection experiments. Mice were challenged intravenously with 1x10^4^ genome equivalent (GE) of mouse-passaged HBV-DE19 viruses diluted in PBS.

For prophylaxis experiments, 500 μg of monoclonal antibody was administered intraperitoneally at 20 and again at 6 hours before infection. For therapy experiments, huFNRG mice with established HBV infections (<10^8^ DNA copies per ml of serum) were injected with 500 μg of each monoclonal antibody intraperitoneally 3 times per week.

DNA in mouse serum collected weekly was extracted using a QIAamp DNA Blood Mini Kit (Qiagen). Total HBV DNA was determined by quantitative PCR (Michailidis et al., 2017). PCR was performed using a TaqMan Universal PCR Master Mix (Applied Biosystems), primers and probe (Table S3).

To obtain HBV DNA from serum for sequence analysis the S domain was amplified using primers (Table S3), and Phusion DNA polymerase (Thermo Fisher Scientific). Initial denaturation was at 98°C for 30 s, followed by 40 amplification cycles (98°C for 10 s, 60°C for 30 s, and 72°C for 30 s), followed by one cycle at 72°C for 5 min. A ~700 bp fragment was gel extracted for Sanger sequencing. Sequence alignments were performed using MacVector.

**QUANTIFICATION AND STATISTICAL ANALYSIS**

The detailed information of statistical analysis could be found in the Result and Figure Legends. Correlation was evaluated by Spearman’s rank correlation method (Figure 1A and 2B). Statistical significance was calculated by Dunn's Kruskal-Wallis multiple comparisons with p values corrected with the Benjamini-Hochberg procedure (Figure S1C). The 50% effective concentration (EC_50_) values by ELISA assays (Figure 3A and 3C) and 50% inhibitory concentration (IC_50_) values by neutralization assays (Figure 5C) were calculated by nonlinear regression analysis in PRISM software.

Adams, P.D., Afonine, P.V., Bunkoczi, G., Chen, V.B., Davis, I.W., Echols, N., Headd, J.J., Hung, L.W., Kapral, G.J., Grosse-Kunstleve, R.W.*, et al.* (2010). PHENIX: a comprehensive Python-based system for macromolecular structure solution. Acta Crystallogr D Biol Crystallogr *66*, 213-221.

Cai, D., Mills, C., Yu, W., Yan, R., Aldrich, C.E., Saputelli, J.R., Mason, W.S., Xu, X., Guo, J.T., Block, T.M.*, et al.* (2012). Identification of disubstituted sulfonamide compounds as specific inhibitors of hepatitis B virus covalently closed circular DNA formation. Antimicrob Agents Chemother *56*, 4277-4288.

Collaborative Computational Project, N. (1994). The CCP4 suite: programs for protein crystallography. Acta Crystallogr D Biol Crystallogr *50*, 760-763.

de Jong, Y.P., Dorner, M., Mommersteeg, M.C., Xiao, J.W., Balazs, A.B., Robbins, J.B., Winer, B.Y., Gerges, S., Vega, K., Labitt, R.N.*, et al.* (2014). Broadly neutralizing antibodies abrogate established hepatitis C virus infection. Sci Transl Med *6*, 254ra129.

Emsley, P., Lohkamp, B., Scott, W.G., and Cowtan, K. (2010). Features and development of Coot. Acta Crystallogr D Biol Crystallogr *66*, 486-501.

Escolano, A., Gristick, H.B., Abernathy, M.E., Merkenschlager, J., Gautam, R., Oliveira, T.Y., Pai, J., West, A.P., Jr., Barnes, C.O., Cohen, A.A.*, et al.* (2019). Immunization expands B cells specific to HIV-1 V3 glycan in mice and macaques. Nature *570*, 468-473.

Gitlin, A.D., von Boehmer, L., Gazumyan, A., Shulman, Z., Oliveira, T.Y., and Nussenzweig, M.C. (2016). Independent Roles of Switching and Hypermutation in the Development and Persistence of B Lymphocyte Memory. Immunity *44*, 769-781.

Jeong, J.Y., Yim, H.S., Ryu, J.Y., Lee, H.S., Lee, J.H., Seen, D.S., and Kang, S.G. (2012). One-step sequence- and ligation-independent cloning as a rapid and versatile cloning method for functional genomics studies. Appl Environ Microbiol *78*, 5440-5443.

Kabsch, W. (2010a). Integration, scaling, space-group assignment and post-refinement. Acta Crystallogr D Biol Crystallogr *66*, 133-144.

Kabsch, W. (2010b). Xds. Acta Crystallogr D Biol Crystallogr *66*, 125-132.

Lefranc, M.P., Giudicelli, V., Duroux, P., Jabado-Michaloud, J., Folch, G., Aouinti, S., Carillon, E., Duvergey, H., Houles, A., Paysan-Lafosse, T.*, et al.* (2015). IMGT(R), the international ImMunoGeneTics information system(R) 25 years on. Nucleic Acids Res *43*, D413-422.

Mayer, C.T., Gazumyan, A., Kara, E.E., Gitlin, A.D., Golijanin, J., Viant, C., Pai, J., Oliveira, T.Y., Wang, Q., Escolano, A.*, et al.* (2017). The microanatomic segregation of selection by apoptosis in the germinal center. Science *358*.

McCoy, A.J., Grosse-Kunstleve, R.W., Adams, P.D., Winn, M.D., Storoni, L.C., and Read, R.J. (2007). Phaser crystallographic software. J Appl Crystallogr *40*, 658-674.

Michailidis, E., Pabon, J., Xiang, K., Park, P., Ramanan, V., Hoffmann, H.H., Schneider, W.M., Bhatia, S.N., de Jong, Y.P., Shlomai, A.*, et al.* (2017). A robust cell culture system supporting the complete life cycle of hepatitis B virus. Sci Rep *7*, 16616.

Michailidis, E., Vercauteren, K., Mancio-Silva, L., Andrus, L., Jahan, C., Ricardo-Lax, I., Zou, C., Kabbani, M., Park, P., Quirk, C.*, et al.* (2020). Expansion, in vivo-ex vivo cycling, and genetic manipulation of primary human hepatocytes. Proc Natl Acad Sci U S A.

Mouquet, H., Scharf, L., Euler, Z., Liu, Y., Eden, C., Scheid, J.F., Halper-Stromberg, A., Gnanapragasam, P.N., Spencer, D.I., Seaman, M.S.*, et al.* (2012). Complex-type N-glycan recognition by potent broadly neutralizing HIV antibodies. Proc Natl Acad Sci U S A *109*, E3268-3277.

Robbiani, D.F., Bozzacco, L., Keeffe, J.R., Khouri, R., Olsen, P.C., Gazumyan, A., Schaefer-Babajew, D., Avila-Rios, S., Nogueira, L., Patel, R.*, et al.* (2017). Recurrent Potent Human Neutralizing Antibodies to Zika Virus in Brazil and Mexico. Cell *169*, 597-609 e511.

Salisse, J., and Sureau, C. (2009). A function essential to viral entry underlies the hepatitis B virus "a" determinant. J Virol *83*, 9321-9328.

Tiller, T., Meffre, E., Yurasov, S., Tsuiji, M., Nussenzweig, M.C., and Wardemann, H. (2008). Efficient generation of monoclonal antibodies from single human B cells by single cell RT-PCR and expression vector cloning. J Immunol Methods *329*, 112-124.

von Boehmer, L., Liu, C., Ackerman, S., Gitlin, A.D., Wang, Q., Gazumyan, A., and Nussenzweig, M.C. (2016). Sequencing and cloning of antigen-specific antibodies from mouse memory B cells. Nat Protoc *11*, 1908-1923.

Wardemann, H., Yurasov, S., Schaefer, A., Young, J.W., Meffre, E., and Nussenzweig, M.C. (2003). Predominant autoantibody production by early human B cell precursors. Science *301*, 1374-1377.

Ye, J., Ma, N., Madden, T.L., and Ostell, J.M. (2013). IgBLAST: an immunoglobulin variable domain sequence analysis tool. Nucleic Acids Res *41*, W34-40.

Yurasov, S., Wardemann, H., Hammersen, J., Tsuiji, M., Meffre, E., Pascual, V., and Nussenzweig, M.C. (2005). Defective B cell tolerance checkpoints in systemic lupus erythematosus. J Exp Med *201*, 703-711.
